# A Dynamical Digital Twin Unmasks Hidden Neuromotor Control Policies and Catastrophic Tipping Points in Parkinson’s Disease

**DOI:** 10.64898/2026.03.09.710685

**Authors:** Kazuki Matsui, Yasuyuki Suzuki, Charles E. Smith, Toru Nakamura, Takuyuki Endo, Saburo Sakoda, Pietro G. Morasso, Taishin Nomura

## Abstract

How the central nervous system maintains upright stance—and why this capability collapses in Parkinson’s disease (PD)—remains a fundamental open question in computational neuroscience. A primary barrier is a severe topological degeneracy between observable sway kinematics (*z*-space) and latent neural control policies (*μ*-space), causing radically distinct control strategies to mask as identical sway patterns. Here, we establish a dynamical digital twin framework bridging empirical sway time series with mechanistic physical models. Assimilating a large cohort (*N* = 1, 038) into an intermittent control model via Bayesian inference, we construct a bidirectional mapping (*z* ↔ *μ*) that unmasks these hidden dynamics. We prove that healthy stance is universally governed by flexible intermittent control near an optimal intermittency ratio (*ρ* ≈ 0.5), whereas PD progression reflects a structural regression toward rigid continuous control (*ρ →* 1). Crucially, parameter topology in *μ*-space reveals that neural control policies reside on a folded low-dimensional manifold separated by a distinct parameter gap; attractor bifurcation analysis demonstrates that a catastrophic tipping point is embedded within this void, proving that parkinsonian postural breakdown is driven by a discontinuous dynamical phase transition rather than continuous control decay. By unmasking hidden disease severity beneath degenerate sway phenotypes, our framework reframes parkinsonian motor failure as an attractor bifurcation on a neural control manifold, providing a modern computational realization of the classical “dynamical disease” paradigm.

**Significance:** Postural collapse in Parkinson’s disease presents a profound neurobiological paradox: cellular degeneration progresses continuously over decades, yet clinical motor failure strikes as a sudden, catastrophic collapse. We resolve this paradox by demonstrating that motor breakdown is not a passive decay of control resources, but a discontinuous dynamical phase transition (attractor bifurcation) triggered when underlying neural control policies cross a latent parameter gap. By unmasking “phenotypic degeneracy”—where radically different neural control strategies produce identical body sway—our digital twin framework redefines neurodegenerative motor failure as a qualitative regime shift in complex control dynamics, offering a universal paradigm for predicting catastrophic transitions in neurological disorders.

---

How the central nervous system maintains human upright stance—and why this fundamental capability abruptly collapses in Parkinson’s disease (PD)—remains a central open question in computational neuroscience and non-equilibrium motor control (1, 2). While clinical management relies predominantly on observational scales (3), the underlying motor failure presents a profound paradox: neurodegenerative processes progress gradually over time at the cellular level (4, 5), yet clinical postural instability often manifests as sudden, non-smooth, and catastrophic deterioration (6–9). Furthermore, observable sway phenotypes display striking heterogeneity, ranging from pathologically enlarged sway areas to paradoxically restricted, rigid sway that equals or is even smaller than that of healthy individuals (10–13). Conventional sway metrics operate strictly on observable kinematics, leaving the latent, nonlinear neural control policies mechanistically unobservable and clinical heterogeneity unresolved (14, 15).

A fundamental barrier to resolving this paradox has been the lack of a mechanistic framework capable of mathematically mapping empirical behavioral phenotypes to underlying physical control policies across large populations (16–18). Human upright stance is inherently unstable due to gravitational toppling torque, sensory feedback delays, and intrinsic neural noise (19, 20). To achieve robust stabilization, theoretical neuroscience posits that the brain employs an intermittent control policy—a state-dependent switching mechanism between active feedback control and unassisted drift along stable manifolds (21–23). Although intermittent control provides an energy-efficient solution for balance (24, 25), whether this principle universally governs real-world human stance across large population cohorts, and how this control policy dynamically degenerates under neurodegenerative stress, remains unproven. Crucially, it remains unknown whether parkinsonian postural collapse reflects a continuous, quantitative degradation of control efficacy or a qualitative, catastrophic dynamical phase transition driven by underlying bifurcation mechanisms.

To address these challenges, we establish a dynamical digital twin framework (Fig. 1A) that assimilates empirical postural sway time series into a mechanistic intermittent control model via Bayesian inference across a large-scale digital twin dataset (*N* = 1, 038). This framework maps each individual into two mathematically coupled domains: an observable phenotypic sway space (***z***-space) capturing high-dimensional wave morphology, and a latent control policy space (***μ***-space) defining mechanistic neural parameters, including control intermittency (*ρ*), feedback gains, and neural delay (Fig. 1A). Through this dual-space representation, we resolve the fundamental origin of clinical ambiguity: observable sway phenotypes in ***z***-space suffer from severe topological degeneracy, where radically distinct neural control policies collapse onto overlapping kinematic patterns (Fig. 1B, Patients P1 vs. P2). By establishing a bidirectional neural mapping (***z*** *↔* ***μ***), our framework unmasks these hidden dynamics, projecting degenerate kinematics back onto interpretable control parameters in ***μ***-space (Fig. 1B).

**Fig. 1.**
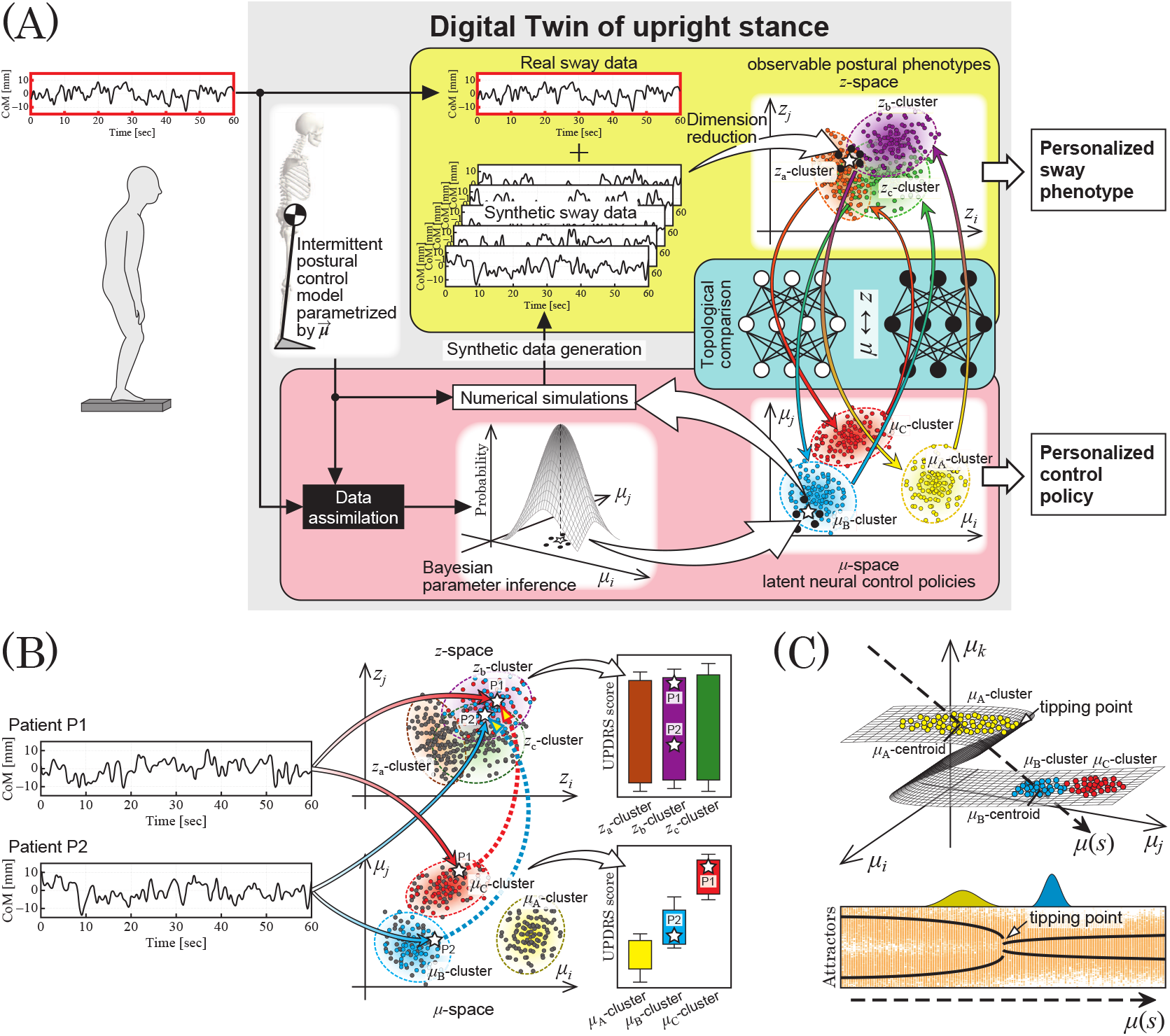
Dynamical digital twin framework for unmasking latent neural control policies and catastrophic phase transitions in Parkinsonian postural collapse. (A) Schematics of the data-driven digital twin framework. Empirical center of mass (CoM) sway data (top left) are assimilated into a mechanistic intermittent control model using Bayesian inference (ABC-SMC) to estimate the posterior distribution of control parameters (***μ***). Synthetic sway time series generated from the posterior distributions are combined with empirical data to form a large-scale digital twin dataset (*N* = 1, 038). This dataset spans two coupled domains: an observable phenotypic space (***z***-space, yellow domain) constructed via dimension reduction of empirical sway metrics, and a latent control policy space (***μ***-space, pink domain). A bidirectional neural network mapping (***z*** *↔* ***μ***) resolves the topological relationships between observable phenotypes and underlying control mechanisms. (B) Resolution of phenotypic degeneracy via bidirectional mapping. Patients P1 and P2 exhibit nearly identical observable sway kinematics in ***z***-space (collapsing into overlapping clusters, e.g., ***z***_*b*_-cluster), rendering conventional clinical metrics insensitive to disease severity (top right). Bidirectional inverse mapping (***z*** *↔* ***μ***) projects these degenerate phenotypes back onto distinct clusters in latent ***μ***-space (***μ***_*C*_ vs. ***μ***_*B*_), unmasking hidden neurological deterioration that closely correlates with clinical motor scores (UPDRS-III; bottom right). (C) Folded control manifold and catastrophic phase transition. In ***μ***-space, neural control policies reside on a folded low-dimensional manifold separated by a distinct parameter void (top). A critical tipping point (attractor bifurcation) is embedded within this structural gap along the disease progression trajectory (***μ***(*s*)). As neurodegeneration advances, the central nervous system loses the dynamic stability of the intermittent control regime, precipitating a catastrophic jump across the parameter void to a pathologically altered attractor (bottom).

Leveraging this digital twin framework, we uncover the fundamental dynamical principles governing human standing and its neurological decay. First, across a massive cohort, we empirically prove that healthy human stance is universally governed by flexible intermittent control operating near an optimal intermittency ratio (*ρ* ≈ 0.5), whereas PD progression driven by neurodegeneration induces a structural regression toward rigid continuous control (*ρ →* 1). Second, parameter topology in ***μ***-space reveals that latent neural control policies reside on a folded low-dimensional manifold interrupted by a pronounced parameter gap devoid of empirical data (Fig. 1C, top). Through attractor bifurcation analysis along disease progression trajectories, we prove that a critical tipping point—marking the boundary between deterministic and stochastic stability of intermittent control— is embedded precisely within this structural gap (Fig. 1C). This demonstrates that parkinsonian postural breakdown is fundamentally a discontinuous phase transition: as neurodegeneration progresses, the central nervous system loses its ability to sustain intermittent dynamics, forcing a catastrophic jump in control space toward pathologically altered attractors (Fig. 1C, bottom).

In doing so, our data-driven digital twin provides a modern computational realization of the classical “dynamical disease” paradigm (26). By demonstrating that parkinsonian motor breakdown stems from an attractor bifurcation on a neural control manifold, this study shifts the paradigm from viewing neurodegenerative symptoms as a passive loss of neural resources to a qualitative, non-smooth phase transition in complex neural control dynamics.

## Results

### Data assimilation based on Bayesian parameter inference

To visualize the latent neural control mechanisms underlying observable postural sway phenotypes, anterior-posterior (AP) center-of-mass (CoM) time-series data collected from 140 patients with Parkinson’s disease (PD) and 59 healthy elderly control subjects were assimilated into a mechanistic intermittent control model. This data assimilation identified the Bayesian posterior probability distribution of model parameters ***μ*** for each participant. In this study, we adopted an intermittent control model that describes human quiet standing posture in the sagittal plane (20, 22). Upright stance is modeled as an inverted single pendulum with a pin joint at the ankle, where the tilt angle and angular velocity relative to the upright vertical are defined as *θ* and *ω*, respectively. The intermittent control framework operates as a state-dependent switching hybrid control system, in which the dynamics of the pendulum are selectively driven by either a delayed or non-delayed differential equation depending on the delayed state vector (*θ*(*t −* Δ), *ω*(*t −* Δ))*T* (switched hybrid control system). Specifically, when the delayed state is located within the active feedback region (ON region) (*θ*(*t* − Δ), *ω*(*t* − Δ))^T^ ∈ *S*_ON_, the system follows the following equation of motion:

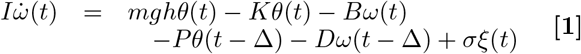

On the other hand, when the delayed state is located within the non-assisted region (OFF region) (*θ*(*t −* Δ), *ω*(*t −* Δ))*T*∈ *S*_OFF_, the dynamics of the system are simplified as follows:

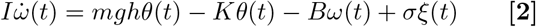

Here, *t* denotes time, *I* is the body moment of inertia around the ankle joint, *m* is the body mass, *g* is the gravitational acceleration, *h* is the height from the ankle joint to the CoM, *K* and *B* represent the passive muscular stiffness and damping coefficients of the ankle joint, *σξ*(*t*) is zero-mean Gaussian white noise, and Δ denotes the neural feedback delay time. Furthermore, *P* and *D* are the proportional and derivative feedback gains of active delayed control, respectively, with the proportional gain normalized as *p* = *P/mgh*.

Figure 2 shows representative examples of stochastic dynamical simulations (synthetic data) generated via data assimilation alongside empirically measured postural sway trajectories. The data assimilation pipeline inferred Bayesian posterior distributions of model parameters by minimizing the distance (statistical divergence) between empirical time series and model-generated synthetic time series (SI Appendix Fig. S11) (25, 27–29). The empirical and synthetic trajectories demonstrated remarkably high agreement in the time-domain profiles (Fig. 2, left), and the model successfully captured the switching dynamics between *S*_ON_ and *S*_OFF_ from its phase portrait (Fig. 2, right). The geometric boundary separating *S*_ON_ and *S*_OFF_ is defined by the intermittent control ratio *ρ* = *S*_ON_/(*S*_ON_ + *S*_OFF_) and the dead-zone radius *r* around the origin. The identification of these six neural control parameters ***μ*** = (*P, D, ρ*, Δ, *σ, r*) enabled person-level mechanistic interpretation and quantification of diverse postural sway characteristics across healthy individuals and the PD cohort (25).

**Fig. 2.**
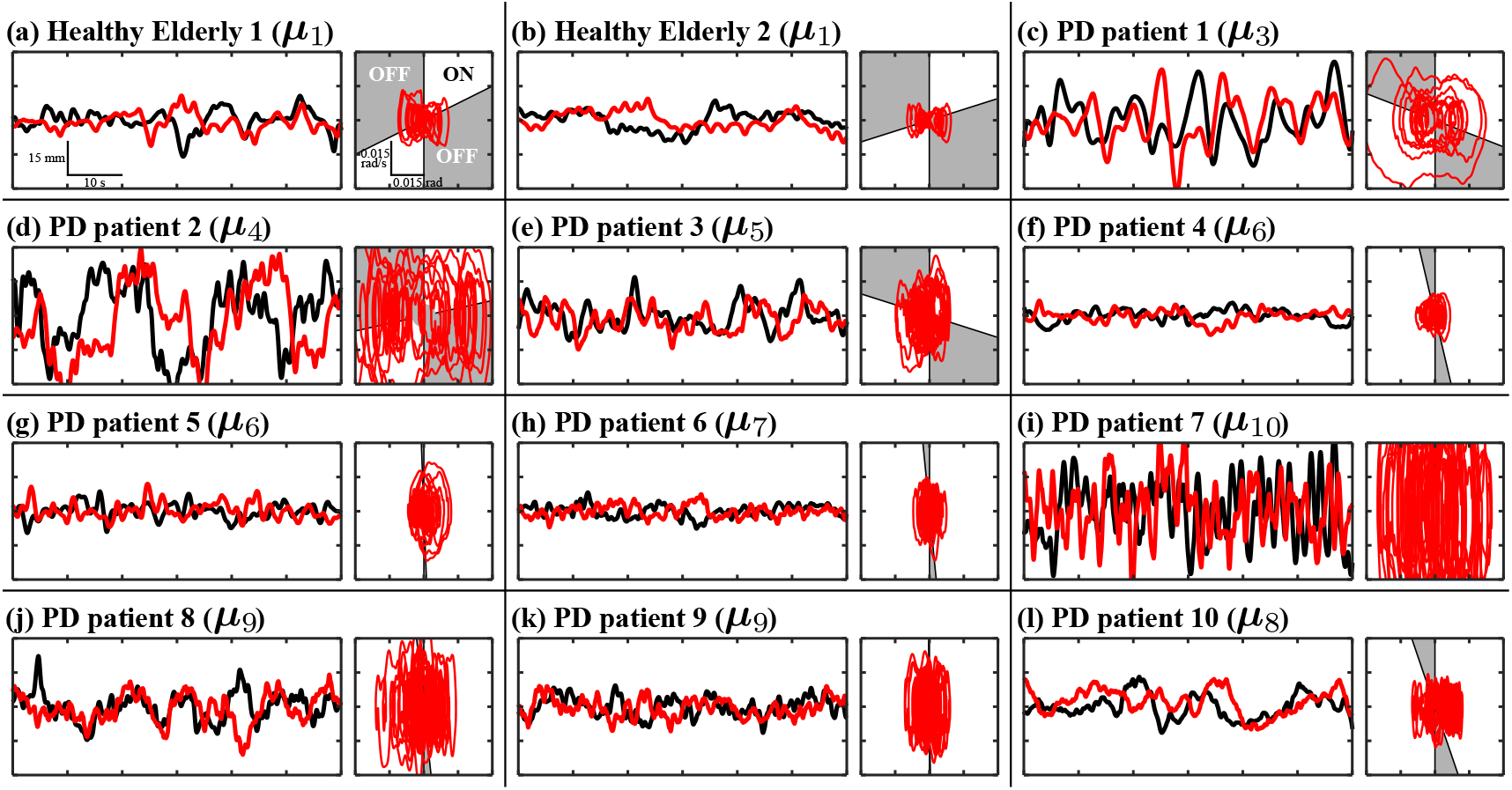
Representative postural sway waveforms of participants. The left panel shows the AP CoM sway time series over 60 s (centered at zero mean sway). The black line represents empirical data, and the red line represents synthetic data. The vertical axis (CoM position) is displayed within a range of *±*30 mm. The right panel shows the synthetic data trajectory in the state plane composed of the angle from vertical and the angular velocity of the inverted pendulum in the intermittent control model. Active control is ON in the white region and OFF in the gray region. The display range is *±*0.03 rad for the horizontal axis and *±*0.03 rad/s for the vertical axis.

### Relationship between data point distribution in model parameter space and disease severity

Data assimilation yielded stable posterior parameter distributions for a total of 173 participants (120 out of 140 patients with PD and 53 out of 59 healthy older adults). All subsequent downstream analyses were grounded in these 173 validated individuals. To enhance spatial resolution within the continuous parameter manifold, we constructed a hybrid digital-twin dataset comprising *N* = 1, 038 data points (173 empirical plus 865 synthetic trials) by drawing five independent posterior samples for each participant. Crucially, this augmentation does not constitute conventional statistical data imputation; rather, it represents a physical digital-twin ensemble analysis driven by model trajectories generated directly from the Bayesian posterior distributions.

Hierarchical clustering analysis in the five-dimensional control parameter space ***μ*** = (*p* = *P/mgh, D, ρ*, Δ, *σ*), excluding the dead-zone radius *r* which exhibits large intra-individual variance, identified 10 qualitatively distinct control strategy clusters (***μ***_1_, ***μ***_2_, …, ***μ***_10_) (Fig. 3(A)(B); see the SUM rows of SI Appendix, Tables S1 and S2 for the constituent breakdown of ***μ***-clusters). From a dynamical systems perspective, these 10 clusters aggregated primarily into three major control regimes (Fig. 3(B)):

**Fig. 3.**
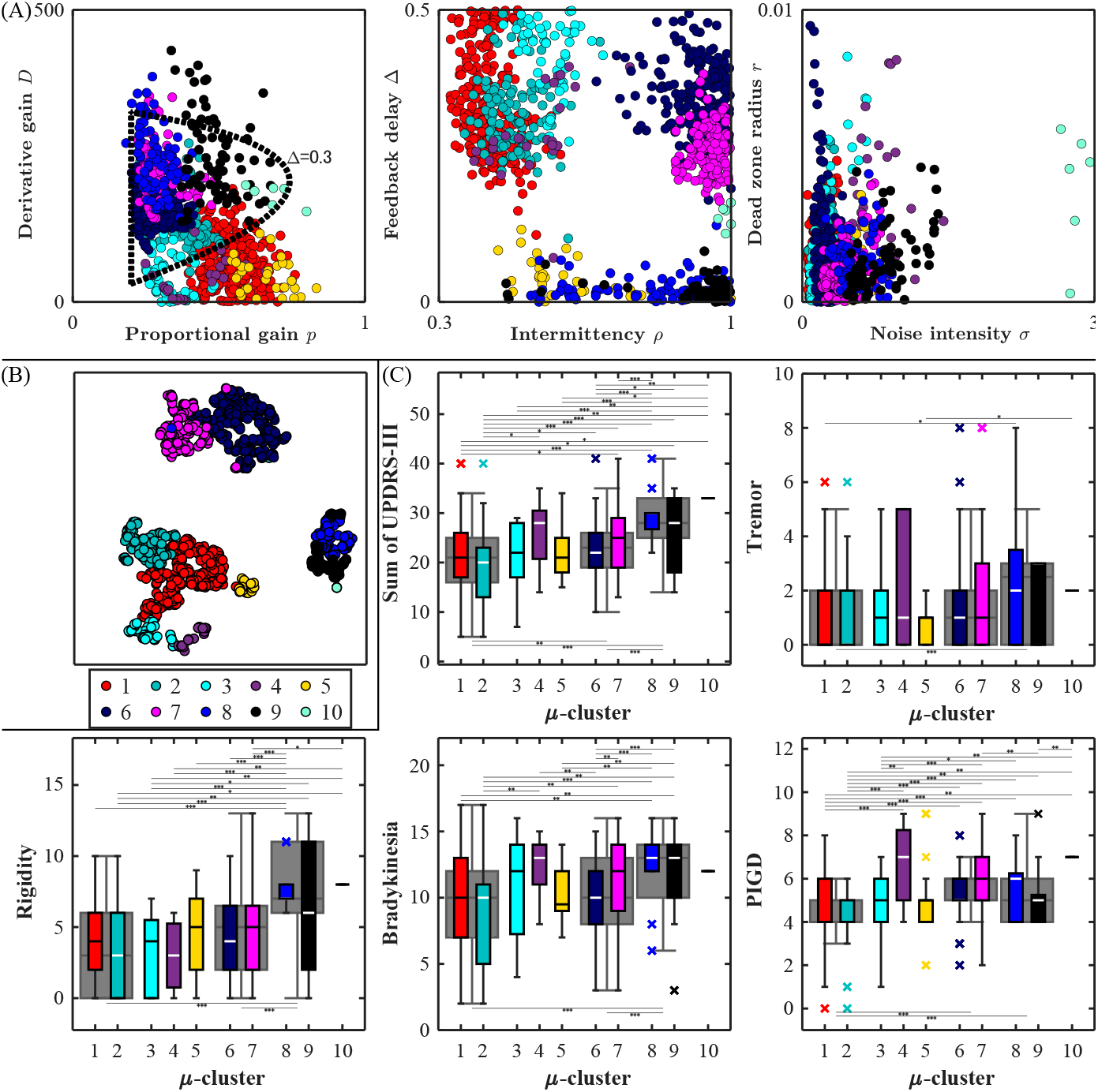
(A) Distribution of ***μ*** values on two-dimensional parameter planes in the six-dimensional space including *r*. The dashed region in the *p*-*D* plane represents the stable region of the pendulum when active control is continuously applied at Δ = 0.3. (B) Visualization of 10 clusters in the five-dimensional ***μ*** = (*p* = *P/mgh, D, ρ*, Δ, *σ*) space using t-SNE. (C) Clinical scores among PD patients in each ***μ***-cluster (Total, Tremor, Rigidity, Bradykinesia, and PIGD scores). The lines above the boxplots represent comparisons among the 10 clusters, whereas the lines below represent comparisons among the three broad cluster groups (three gray boxes). *: *p*_*bf*_ *<* 0.05, **: *p*_*bf*_ *<* 0.01, ***: *p*_*bf*_ *<* 0.001, where *p*_*bf*_ denotes the *p*-value of the Mann–Whitney U test with Bonferroni correction.

1. ***M***_INT_ (***μ***_1_ ∪ ***μ***_2_): “Intermittent active control” (*ρ* ∼ 0.5), characterized by functionally exploiting the OFF region (passive region).
2. ***M***_CONT_ (***μ***_6_ ∪ ***μ***_7_): “Continuous active control” (*ρ →* 1), in which intermittency collapses, resulting in persistent feedback intervention.
3. ***M***_STIFF_ (***μ***_8_ ∪ ***μ***_9_): “Stiffness/passive control,” exhibiting near-zero neural delay (Δ ∼ 0) and generating high passive mechanical impedance through antagonist muscle co-contraction.

Comparison of clinical motor severity across these clusters (Total, Tremor, Rigidity, Bradykinesia, and Postural Instability and Gait Difficulty (PIGD) scores based on UPDRS Part III; Kruskal–Wallis test and Bonferroni-corrected Mann–Whitney U test, Fig. 3(C)) revealed statistically significant differences across all motor domains (SI Appendix, Tables S3–S13). In particular, the PIGD score—a primary clinical marker of postural breakdown— demonstrated significant discriminative power across the largest number of cluster pairs. These results demonstrate that neurological motor decline in PD is directly linked to structural transformations of latent neural control strategies (namely, the breakdown of intermittency and regression to passive stiffness strategies).

### Limitations of conventional sway indices and chaotic distribution in phenotypic latent space

Next, we evaluated the diagnostic efficacy of conventional observational approaches that rely strictly on observable kinematic sway metrics. For each time series in the dataset, 18 standard postural sway features (RMS, Power, DFA/SDA scaling exponents, etc.) were computed, and six latent factors (*z*_1_ to *z*_6_) were extracted via factor analysis (Fig. 4). Each factor reflects low-frequency power ratio (*z*_1_), total trajectory length and sway amplitude (*z*_2_), DFA/SDA critical transition points (*z*_3_, *z*_4_), high-frequency scaling exponent (*z*_5_), and velocity zero-crossing interval distribution (*z*_6_).

**Fig. 4.**
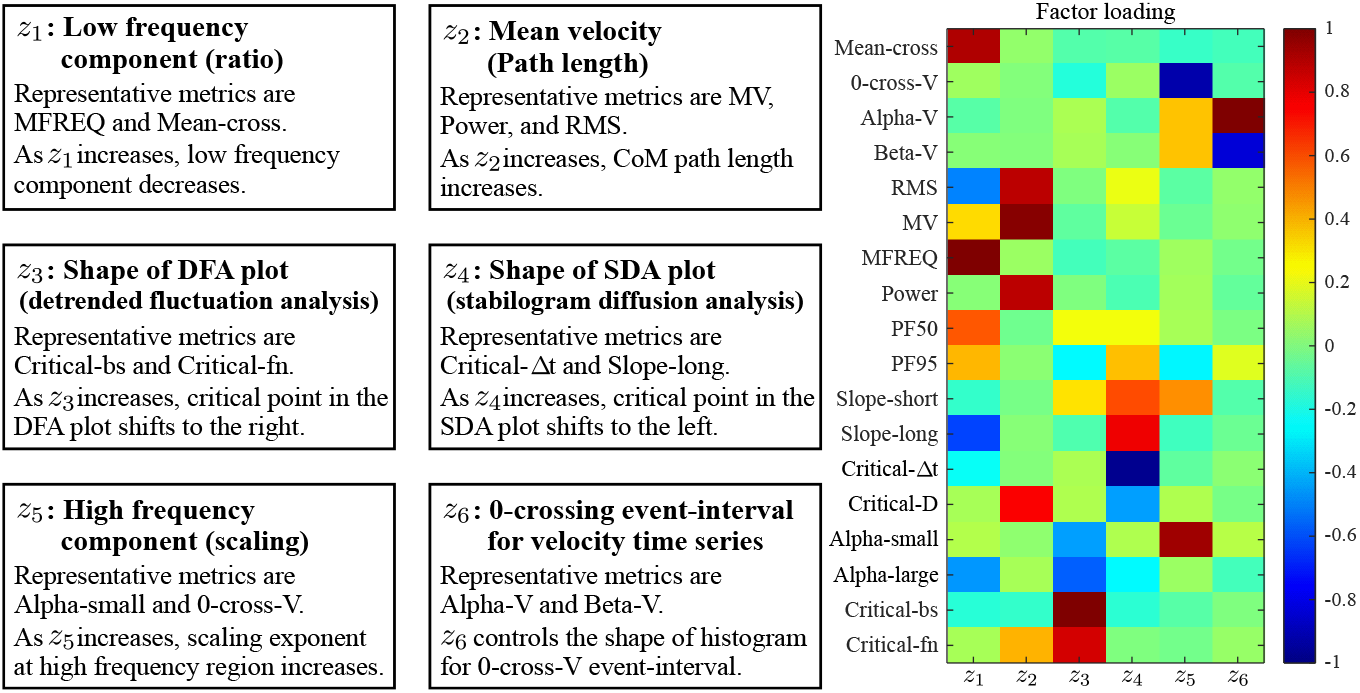
Overview of extracted latent factors (left) and color map of factor loadings (right). In the color map, colors closer to red and blue indicate that the latent factor has a larger positive or negative influence on the original index, respectively.

Hierarchical clustering within this six-dimensional factor space (***z***-space) identified 10 phenotypic sway clusters (***ζ***_1_ to ***ζ***_10_) (Fig. 5(A)(B); see the SUM columns of SI Appendix, Tables S1 and S2 for the constituent breakdown of ***ζ***-clusters). However, unlike the ***μ***-space, which exhibited clear geometric separation (Fig. 3(B)), clusters in ***z***-space demonstrated severe topological overlap and ambiguous boundaries, reflecting profound “phenotypic degeneracy” (a phenomenon wherein radically different internal states generate identical outward phenotypes) (Fig. 5(B)). Furthermore, comparisons of clinical scores among ***z***-clusters (Fig. 5(C)) revealed a dramatic reduction in the number of significant cluster pairs across major motor domains compared to ***μ***-space (SI Appendix, Tables S14–S19). Notably, cluster pairs exhibiting significant differences in PIGD score dropped substantially from 17 pairs in ***μ***-space to 11 pairs in ***z***-space. These findings indicate that data-driven analyses relying solely on superficial sway metrics (***z***-space) are confounded by phenotypic heterogeneity and fail to accurately stratify the clinical severity of postural impairment.

**Fig. 5.**
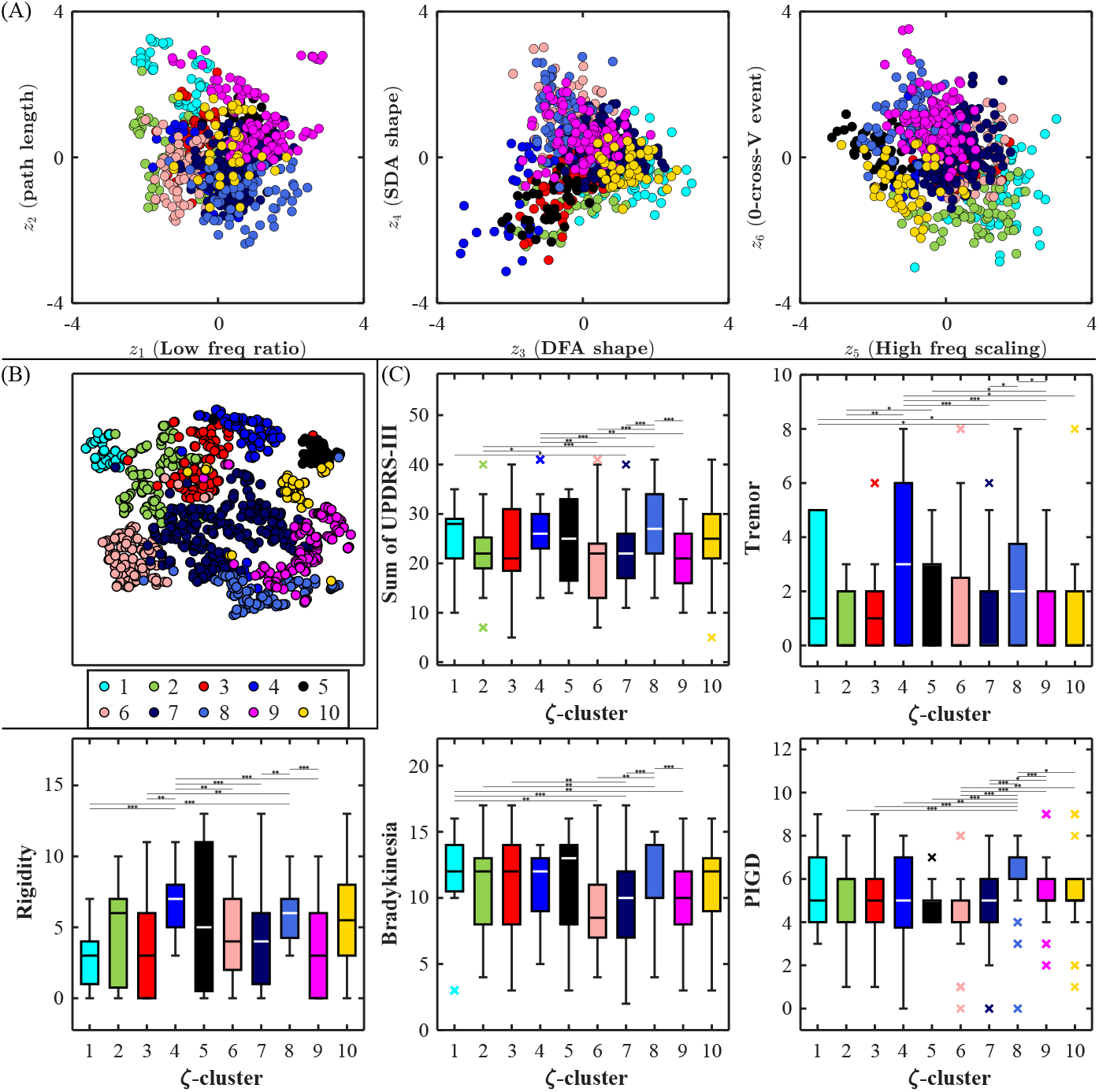
(A) Distribution of ***z*** values for each cluster. Circles: PD patients, Triangles: healthy elderly. (B) Visualization of clusters in the latent variable (factor score) ***z*** space using t-SNE. (C) Clinical scores among PD patients in each ***ζ***-cluster. *: *p*_*bf*_ *<* 0.05, **: *p*_*bf*_ *<* 0.01, ***: *p*_*bf*_ *<* 0.001; *p*_*bf*_ denotes the *p*-value of the Mann– Whitney U test with Bonferroni correction.

### Bidirectional mapping between parameter and latent spaces (*μ* ↔ *z*) and unmasking of hidden severity

To resolve phenotypic degeneracy and enable real-time mapping from observable sway metrics to latent control strategies, we constructed fully connected neural networks (NNs) trained to learn forward (***μ*** *→* ***z***) and inverse (***z*** *→* ***μ***) mappings. Leveraging the augmented dataset, both NNs achieved remarkably high predictive accuracy on test data, approaching a bijective (one-to-one) mapping (MSE ≈ 0.359, SI Appendix, Figs. S12–S13 and Tables S20–S23). Using cross-cluster correspondence probabilities generated from surrogate data (transition probability ≥ 33.3%), we established a detailed correspondence matrix between ***μ***-space and ***z***-space (Table 1). While many clusters demonstrated clear one-to-one canonical correspondences (e.g., ***μ***_1_ ↔ ***ζ***_3_, ***μ***_8_ ↔ ***ζ***_4_), other clusters unveiled complex “many-to-one convergent” or “one-to-many divergent” mappings. Detailed descriptions for each ***μ***–***ζ*** pair are provided in SI Appendix C.

**Table 1.** Correspondence table between *μ*- and *ζ*-clusters.

| | $\mu_1$ | $\mu_2$ | $\mu_3$ | $\mu_4$ | $\mu_5$ | $\mu_6$ | $\mu_7$ | $\mu_8$ | $\mu_9$ | $\mu_{10}$ |
| --- | --- | --- | --- | --- | --- | --- | --- | --- | --- | --- |
| $\zeta_3$ | **48.7\|67.3** | 13.3\|22.1 | 1.9\|4.1 | 22.7\|1.9 | 0.0\|0.8 | 2.4\|1.9 | 0.7\|0.8 | 11.9\|1.0 | 1.4\|0.1 | 0.0\|0.0 |
| $\zeta_7$ | 21.9\|11.0 | **60.4\|34.5** | 0.1\|0.6 | 0.0\|0.0 | 2.0\|2.0 | **55.8\|41.1** | 0.4\|8.8 | 0.3\|1.4 | 2.9\|0.5 | 0.0\|0.0 |
| $\zeta_1$ | 1.1\|2.9 | 2.4\|0.1 | **72.9\|54.6** | **71.8\|41.6** | 0.0\|0.4 | 0.8\|0.4 | 0.0\|0.0 | 0.0\|0.0 | 0.0\|0.0 | 0.0\|0.0 |
| $\zeta_2$ | 11.3\|21.8 | 6.4\|19.9 | 24.7\|25.1 | 2.0\|1.7 | 1.5\|7.1 | 2.9\|9.9 | 0.0\|2.0 | 26.6\|12.3 | 0.2\|0.1 | 0.0\|0.0 |
| $\zeta_{10}$ | 0.9\|5.8 | 0.4\|14.0 | 0.0\|0.6 | 1.3\|3.5 | **96.3\|50.0** | 0.1\|0.8 | 0.0\|4.9 | 15.8\|8.2 | 18.8\|12.2 | 0.0\|0.0 |
| $\zeta_8$ | 0.2\|0.1 | 3.3\|3.4 | 0.0\|0.0 | 0.0\|0.0 | 0.1\|0.3 | 8.7\|13.2 | **40.5\|69.6** | 6.4\|8.0 | 9.9\|5.3 | 0.0\|0.0 |
| $\zeta_9$ | 0.1\|1.3 | 3.0\|8.5 | 0.0\|0.2 | 2.2\|3.8 | 0.1\|0.3 | 10.2\|15.8 | **46.6\|65.1** | 0.0\|0.1 | 0.5\|4.3 | **100.0\|0.5** |
| $\zeta_4$ | 6.0\|9.0 | 1.1\|12.7 | 0.0\|0.0 | 0.0\|0.1 | 0.0\|0.1 | 1.9\|11.3 | 5.6\|29.4 | **34.4\|36.9** | 1.8\|0.6 | 0.0\|0.0 |
| $\zeta_5$ | 0.0\|0.0 | 0.0\|0.0 | 0.0\|0.0 | 0.0\|0.0 | 0.1\|0.0 | 0.4\|0.0 | 5.1\|3.9 | 2.7\|4.2 | **64.5\|91.8** | 0.0\|0.0 |
| $\zeta_6$ | 9.8\|26.7 | 9.6\|**41.3** | 0.5\|17.5 | 0.0\|0.1 | 0.0\|0.0 | 16.7\|12.2 | 1.1\|1.4 | 1.8\|0.8 | 0.0\|0.0 | 0.0\|0.0 |

Rows represent $\zeta$ -clusters and columns represent $\mu$ -clusters. Left value: probability [%] that the prediction from the $\mu \rightarrow \zeta$ NN (neural network) for surrogate $\mu$ data belongs to the target $\zeta$ -cluster. Right value: probability [%] in the inverse mapping. Bold values indicate probabilities of 33.3% or higher. Gray cells indicate probabilities of 33.3% or higher in both directions.

### Primary mapping profiles

#### 1) One-to-one canonical mappings

##### Healthy / Mild control μ_1_ ↔ ζ_3_ pair (Fig. 6(a))

Characterized by low gains (*p, D*) and a low active duty cycle (small *ρ*), this pair exploits the stable manifold of the pendulum during control OFF periods to achieve low UPDRS scores and energy-efficient postural control.

**Fig. 6.**
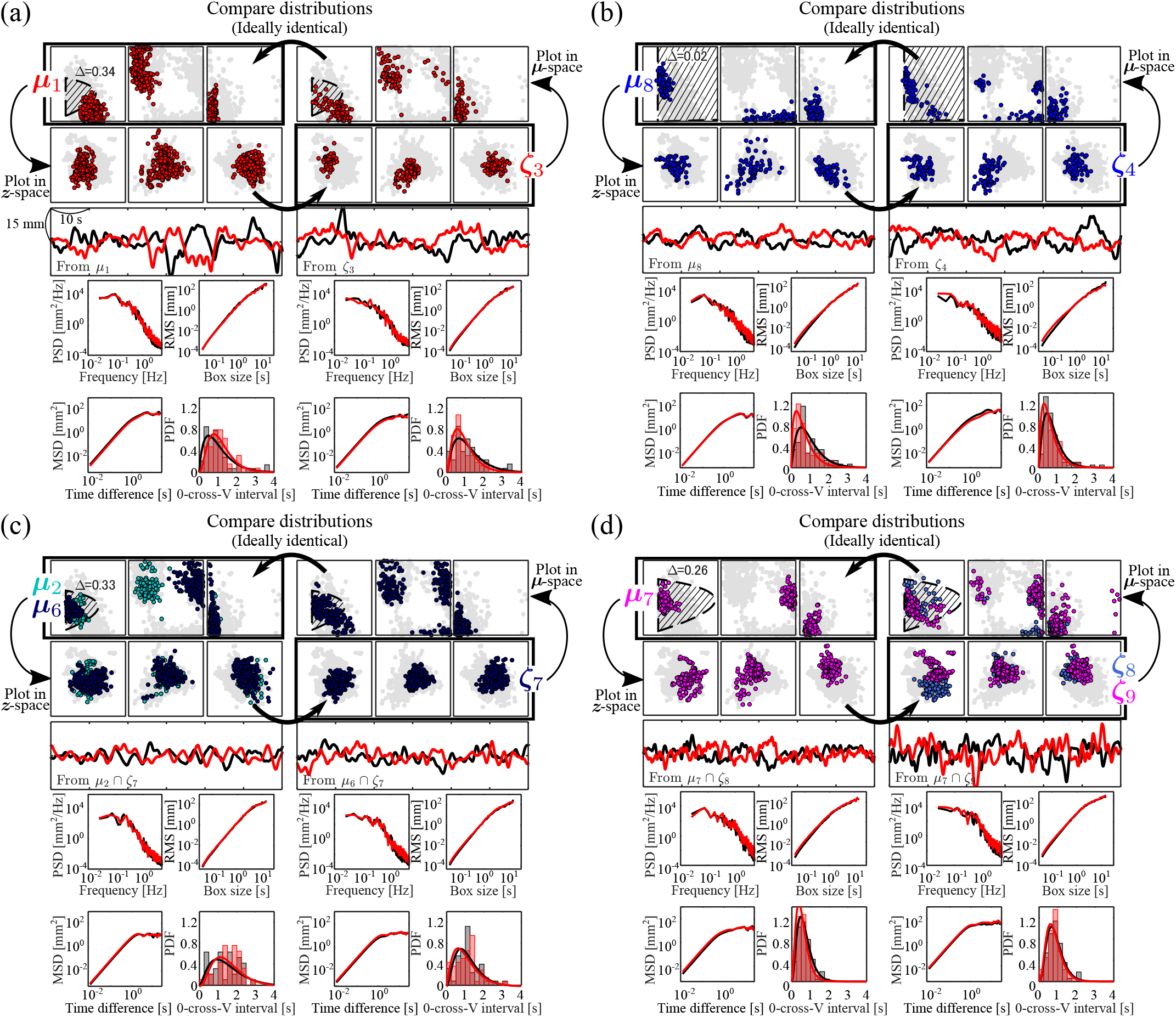
Four representative pairs of ***μ***- and ***ζ***-clusters: (a) ***μ***_1_ –***ζ***_3_ pair, (b) ***μ***_8_ –***ζ***_4_ pair, (c) (***μ***_2_, ***μ***_6_)–***ζ***_7_ pair, and (d) ***μ***_7_ –(***ζ***_8_, ***ζ***_9_) pair ((c) and (d) are shown in Fig. 7). In each of (a)–(d), the top-left and top-right panels show the distributions in ***μ***-space for the ***μ***-cluster and ***ζ***-cluster, respectively. The axes of ***μ***-space are identical to those in Fig. 3(A). The stable region shown in the *p*-*D* plane (hatched region) was calculated using the mean of the distribution of Δ values for each ***μ***-cluster. The left and right panels in the second row show these distributions in ***z***-space. The axes of ***z***-space are identical to those in Fig. 5(A). Panels below the third row show examples of CoM time-series waveforms (third row), power spectra (fourth row left), DFA plots (fourth row right), SDA plots (fifth row left), and velocity sign-change interval histograms (fifth row right) of representative experimental participants extracted from the corresponding clusters (black: empirical data, red: synthetic data).

##### Severe rigidity μ_8_ ↔ ζ_4_ pair (Fig. 6(b))

Characterized by near-zero delay (Δ ∼ 0) and continuous activation (*ρ* ∼ 1), reflecting passive joint stiffening driven by antagonist muscle co-contraction. Patients in this cluster exhibited paradoxically small sway amplitudes (*z*_2_) yet presented the highest Rigidity and Total UPDRS scores, highlighting the diagnostic hazard of judging disease severity purely from kinematic sway amplitude.

#### 2) Many-to-one convergent mapping (Unmasking “hidden severity”)

##### (μ_2_, μ_6_) → ζ_7_ pair (Fig. 7(c))

Two mechanistically distinct control strategies?flexible intermittent control (***μ***_2_) and rigid continuous control (***μ***_6_)?converged onto (and were masked by) identical small-amplitude sway phenotypes (***ζ***_7_). While ***μ***_2_ comprised clinically mild patients, ***μ***_6_ contained severe PD patients with markedly elevated PIGD scores who were forced into continuous feedback due to the breakdown of intermittent dynamics. Conventional sway measurements (***ζ***_7_) risk overlooking these “hidden severe” patients; however, inverse parameter mapping (*z → μ*) instantly unmasks the underlying neurological breakdown.

**Fig. 7.**
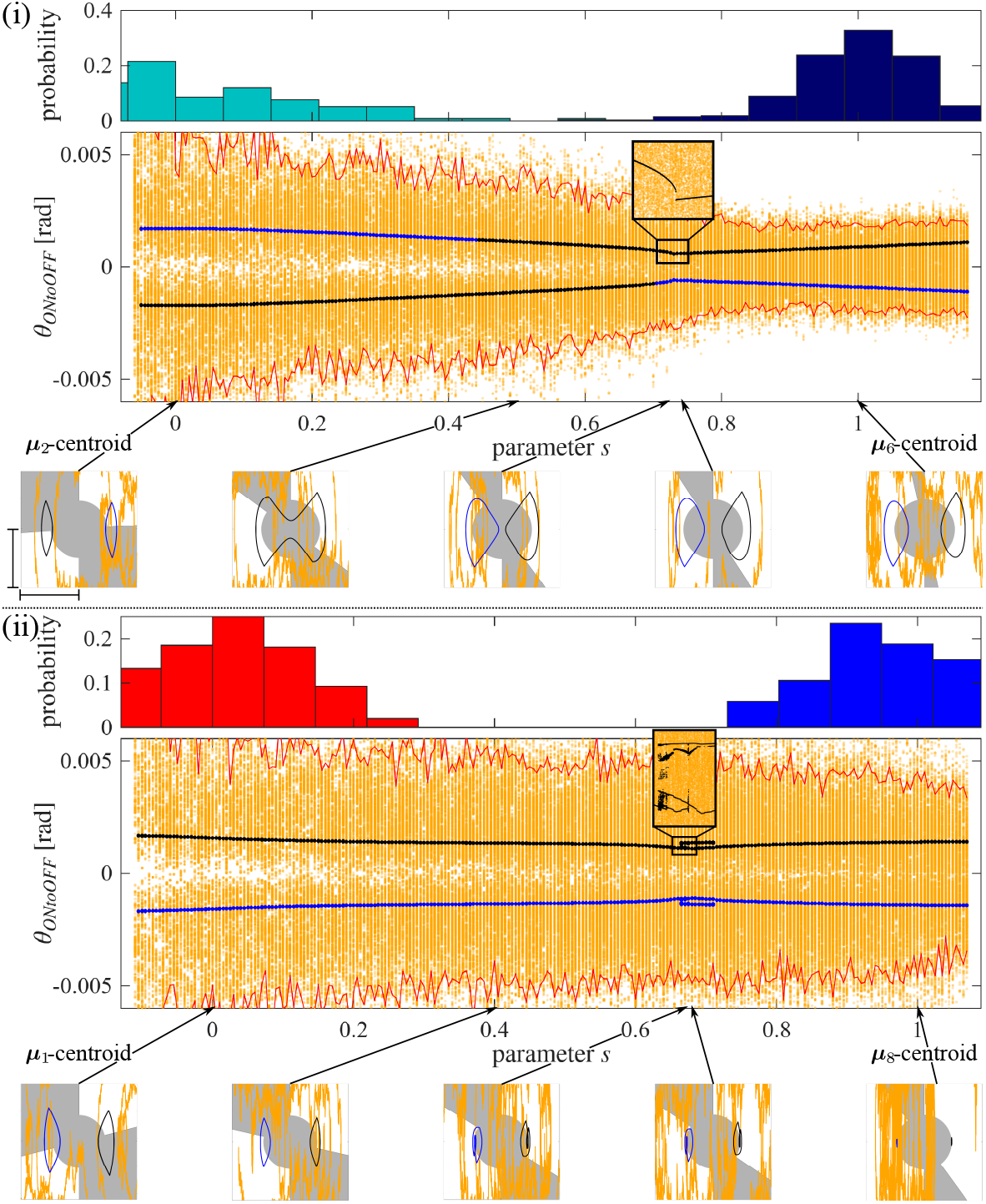
Distribution of data points along a hypothetical linear trajectory in ***μ***-space (representing hypothetical disease severity progression) (top row), bifurcation diagrams of the postural control model (middle row), and sample trajectories at various locations on the linear trajectory (bottom row). (i) ***μ***_2_ *→* ***μ***_6_, (ii) ***μ***_1_ *→* ***μ***_8_ . The top row displays data points belonging to the start and end clusters projected onto the linear trajectory ***μ***(*s*) constructed in ***μ***-space, showing histograms along the parametric variable *s* connecting the centroids of the two clusters. ***μ***(*s* = 0) and ***μ***(*s* = 1) correspond to the centroids of the start and end clusters, respectively. Histograms are normalized per cluster and represented as probabilities. The horizontal axis of the bifurcation diagrams in the middle row is the linear parameter *s*, identical to the top row, whereas the vertical axis represents stroboscopic measurement points (periodic sampling points) of *θ* using the switching boundary (where active control switches from ON to OFF) as a Poincaré map. Black and blue dots represent deterministic simulations calculated from different initial conditions, whereas orange dots represent results from stochastic simulations. Upper and lower red lines indicate the 2.5th and 97.5th percentiles of stochastic measurement points, respectively. In both (i) and (ii), the 2.5th percentile exhibited a significant negative correlation with *s* ((i) *r* = *−* 0.84, *p <* 0.001; (ii) *r* = *−* 0.70, *p <* 0.001), while the 97.5th percentile exhibited a significant positive correlation ((i) *r* = 0.90, *p <* 0.001; (ii) *r* = 0.41, *p <* 0.001). The bottom row shows the state space of the pendulum at each *s* point, displaying steady-state trajectories for 20 s. The horizontal axis of the state space represents pendulum angle, and the vertical axis represents angular velocity; scale bars correspond to 0.003 rad and 0.003 rad/s, respectively.

#### 3) One-to-many divergent mapping and disease subtyping

##### μ_7_ → (ζ_8_, ζ_9_) pair (Fig. 7(d))

A single “high-noise continuous control strategy” (***μ***_7_) diverged into two distinct phenotypes: a suppressed phenotype (***ζ***_8_) and a pathologically amplified phenotype (***ζ***_9_). Patients in ***ζ***_8_ exhibited significantly worse PIGD scores than those in ***ζ***_9_, suggesting a compensatory co-contraction strategy that forcibly suppresses sway under conditions of severe instability.

### Multiple System Atrophy (MSA) specific signature μ_4_ ↔ ζ_1_ pair

Follow-up investigation of cluster ***μ***_4_, characterized by exceptionally high internal noise (*σ*) and elevated PIGD scores, confirmed that all constituent patients were diagnosed with Multiple System Atrophy (MSA). This demonstrates the efficacy of our model-driven approach for the automated differentiation of specific atypical parkinsonisms, such as progressive supranuclear palsy and MSA.

### Disease progression prediction based on attractor bifurcation diagrams

To elucidate how continuous parameter degradation in ***μ***-space reshapes phase-plane attractors and dynamical stability, we constructed bifurcation diagrams along hypothetical disease progression pathways (Fig. 7). Based on the severity gradient observed across ***μ***-clusters (Fig. 3(C)), two primary progression scenarios were established using a linear trajectory ***μ***(*s*) parameterized by *s* ∈ [*−*1, 2]:

**Scenario (i) *μ***_2_ *→* ***μ***_6_ (Transition from intermittent to continuous control)

**Scenario (ii) *μ***_1_ *→* ***μ***_8_ (Transition from flexible control to passive muscle rigidity)

Poincaré map analysis in deterministic simulations (*σ* = 0) revealed non-smooth orbit bifurcations (border-collision bifurcations) and discontinuous jumps at a critical threshold *s ≈* 0.7 in both scenarios (Fig. 8, insets). This critical tipping point marks the boundary where intermittent control loses dynamical stability. Examination of phaseplane trajectories reveals that beyond *s ≈* 0.7, state points no longer enter the control OFF region near the stable manifold, forcing an irreversible transition into continuous active feedback or passive rigidity. Investigating the parameter distribution along ***μ***(*s*) unveiled a prominent “void” around *s ≈* 0.7 (Fig. 8, top histograms). This reveals that neural control strategies reside along low-dimensional folded directions, separated by an unstable parameter gap where intermittent stabilization fails.

In stochastic simulations with physiological noise (*σ* /= 0), the dispersion of Poincaré sampling points (2.5th and 97.5th percentiles, *p <* 0.001, Fig. 8) monotonically contracted as disease severity progressed (*s →* 1). While clinical intuition often equates motor collapse with amplified sway, bifurcation analysis of our digital twin proves a dynamical paradox: as parkinsonian postural instability (PIGD) worsens, the loss of intermittency and compensatory rigidity mechanisms contract the phase-plane attractor, resulting in “pathologically distorted, small-amplitude sway.”

## Discussion

In this study, we applied data assimilation to anterior-posterior postural sway time-series data from patients with Parkinson’s disease (PD) and healthy elderly controls, identifying their latent neural control parameter space (***μ***-space). This mechanistic digital-twin approach resolved the phenomenon of “phenotypic degeneracy,” which remained uncapturable through conventional observational sway metrics (***z***-space), and demonstrated the ability to precisely extract “hidden disease severity” and specific clinical subtypes (such as MSA) embedded within the pathology. Below, we discuss the neurocontrollability, dynamic phase transitions, and clinical implications of our primary findings.

### Phenotypic degeneracy and dynamic unmasking of “hidden severity”

One of the central discoveries of this study is the elucidation of a “many-to-one convergent mapping (***μ***_2_, ***μ***_6_ *→* ***ζ***_7_),” wherein vastly different internal control strategies (***μ***-space) underlie virtually identical observable postural sway profiles (***z***-space). In clinical practice, a small postural sway amplitude is frequently misinterpreted as reflecting stable and superior postural control. Dynamically, however, a small-amplitude sway profile (***ζ***_7_) can emerge from two fundamentally distinct physical mechanisms:

1. *Flexible intermittent control under noise-suppressed environments* (***μ***_2_) A state (mild PD/healthy) in which the system functionally exploits the stable manifold of the OFF region (passive region), maintaining stance with minimal active intervention. In our model, intermittent control inherently permits drift within the passive region in the presence of typical motor noise levels, thereby spontaneously generating relatively large-amplitude, flexible sway. Under specific conditions where internal noise (*σ*) is exceptionally small, however, this intermittent control dynamic can naturally produce a small-amplitude sway profile (***ζ***_7_).
2. *Pathological over-suppression driven by excessive continuous feedback* (***μ***_6_): A state (severe PD) in which the switching mechanism of intermittent control has collapsed, forcing persistent, high-gain feedback intervention. In this state, unbroken and powerful mechanical restoring forces act continuously, physically constraining the trajectory’s degrees of freedom and forcibly suppressing the outward phenotype into a pathologically small sway profile (***ζ***_7_).

Severe patients belonging to ***μ***_6_ forcibly stiffen and continuously control their joints to compensate for the loss of intermittent switching associated with basal ganglia network dysfunction. Observational diagnostics relying exclusively on conventional sway measurements (***z***-space) face the fatal risk of misclassifying patients with severe postural instability and gait difficulty (PIGD) as having “good balance” due to small sway amplitudes, thereby overlooking them (failure to detect “hidden severity”). The digital-twin-based inverse mapping (***z*** *→* ***μ***) breaks through this phenotypic degeneracy, serving as an innovative, non-invasive tool to accurately visualize hidden neurodynamical collapse beneath superficial sway.

### One-to-many divergent mapping and diversity of compensatory strategies

Furthermore, we identified a “one-to-many divergent mapping” where a single control parameter regime (***μ***_7_) yields two diametrically opposed sway phenotypes (***ζ***_8_: suppressed type vs. ***ζ***_9_: amplified type). Although ***μ***_7_ is characterized by continuous control under high noise, patients in ***ζ***_8_ exhibited significantly worse PIGD scores than those in ***ζ***_9_. This reflects a compensatory adaptive behavior wherein patients facing severe postural instability attempt to physically stifle sway amplitude by augmenting muscle co-contraction. In other words, even under identical internal deficit states, the observed phenotype diverges depending on the individual patient’s choice of compensatory strategy.

Additionally, follow-up investigation of control cluster ***μ***_4_ (corresponding to ***ζ***_1_), which exhibited exceptionally high internal noise (*σ*), revealed that all constituent patients were diagnosed with Multiple System Atrophy (MSA). Capturing distinct signatures within the control parameter space strongly suggests that model-driven automated differential diagnosis among parkinsonian syndromes is clinically feasible.

### Bifurcation structure and “dynamical paradox” during disease progression

Bifurcation analysis utilizing our digital twin (Fig. 7) vividly depicts the phase transitions (tipping points) of the neural control system accompanying disease progression, offering a clear dynamical interpretation of a longstanding clinical mystery. Clinical management of Parkinson’s disease has long relied on observational scales (UPDRS and Hoehn & Yahr). Pathologically, neurode-generative processes such as nigrostriatal dopaminergic loss progress continuously over years to decades at the cellular level (4, 5). In clinical reality, however, patient postural instability and fall risk deteriorate abruptly and catastrophically once reaching H&Y Stage III (6–9).

### Folded low-dimensional manifold and parameter gaps

The physical reality underpinning this clinical paradox becomes clear from the topological structure of the identified ***μ***-space. Identified data points across participants are not uniformly distributed; instead, they are distinctly clustered, separated by a discontinuous structural void (parameter gap) between healthy/mild groups (*M*_INT_) and advanced groups (*M*_CONT_, *M*_STIFF_) (Fig. 3(B)). This demonstrates that the neural control policy space available for humans to maintain stable stance is strictly constrained onto a “folded low-dimensional nonlinear manifold” embedded within high-dimensional ***μ***-space.

### Fold structure and catastrophic tipping at s ≈ 0.7

Consider a latent disease severity parameter *s* driving the control system along this low-dimensional compensatory manifold ***μ***(*s*). As pathology progresses, the central nervous system continuously adapts its control policy to preserve postural stability. However, when this trajectory reaches the fold point of the manifold, active compensation reaches its physical limits. This limit is precisely the critical threshold *s ≈* 0.7 identified by our Poincaré map analysis:

#### Catastrophic tipping

The moment *s ≈* 0.7 is crossed, the control system permits no intermediate states and catastrophically jumps to a second stable branch (continuous control *M*_CONT_ or passive stiffness strategy *M*_STIFF_). Because no steady-state control policy can exist within the gap region (Fig. 7, top row), this manifests clinically as a “sudden, discontinuous postural failure (H&Y Stage III).”

#### Border-collision bifurcation and irreversible loss of intermittency

At this phase transition, the state trajectory is physically and permanently expelled from the stable manifold of intermittent control (border-collision/non-smooth bifurcation), irreversibly losing its capacity for flexible dynamical stabilization within the OFF region. Consequently, the control system is forced to undergo structural reorganization into excessive continuous feedback (***μ***_6_) or global muscle rigidity (***μ***_8_).

### Unveiling the truth of instability: The “Dynamical Paradox”

Crucially, our framework resolves a long-standing clinical paradox: as the disease progresses and PIGD severity increases, postural sway often exhibits a counterintuitive shrinkage and rigidity of the dynamic attractor, rather than simple hyper-sway (10, 11, 13). In our model, this attractor contraction reflects a pathological compensatory strategy. As the basal ganglia’s intermittent control capacity declines (captured by parameter shifts in ***μ***_6_ and ***μ***_8_), the central nervous system compensates for the imminent threat of falling by enforcing continuous, high-rigidity feedback via muscle co-contraction. While this compensatory constraint temporarily stabilizes the center of mass near the vertical equilibrium, it severely reduces the system’s dynamic range and phase-space volume. Consequently, this state masks the underlying loss of intermittent adaptability until the system hits a critical tipping point (*s ≈* 0.7)—where even a minor external perturbation forces a sudden, catastrophic transition to postural collapse.

### Clinical necessity and bounded design of mechanistic data augmentation

Another major contribution of our clinical digital-twin framework lies in addressing the ubiquitous “small-data” challenge in clinical cohorts (30) through a mechanics-based strategy. By sampling from the Bayesian posterior distributions of identified control parameters (***μ***-space), we generated highly realistic synthetic trajectories that faithfully mirror empirical nonlinear postural sway dynamics, thereby dramatically enhancing spatial resolution within the control manifold.

As demonstrated in our supplementary analyses (SI Appendix, Figs. S14 and S15), the core topological structures and parameter clustering trends were fully preserved when restricting the analysis solely to the empirical dataset (*N* = 173). However, relying exclusively on unaugmented empirical data severely diminished the statistical power required to detect fine-grained correlations with clinical motor scores, while significantly deteriorating the predictive accuracy of neural networks bridging parameter space and phenotypic space (SI Appendix, Tables S24 and S25). These findings demonstrate that mechanistic data augmentation is a crucial mathematical prerequisite for overcoming high patient heterogeneity and advancing robust, evidence-based precision medicine (31).

On the other hand, explicitly delineating the appropriate boundaries of synthetic data generation is equally critical. Even when leveraging physical models, generating infinite synthetic data or excessively expanding minute, isolated clusters introduces stochastic noise and overfitting risks, potentially compromising inference validity (32). To mitigate this risk, we constrained our augmentation scale based on a pre-calculated theoretical sample size (aggregating 5 trials per participant) and adopted a rigorous evaluation framework measuring practical effect sizes (Cliff’s *δ*). This mechanics-backed, principled data augmentation approach serves as a benchmark standard for data generation in future clinical digital-twin research.

### Clinical implications and therapeutic applications

Beyond pathological characterization, our findings establish a pragmatic foundation for next-generation precision medicine. *Objective, non-invasive biomarkers via digital twins*: By deploying the inverse mapping neural network (*z → μ*), clinicians can estimate an individual’s latent control parameters (***μ***-space) directly from non-invasive, brief (e.g., 60-second) force-plate measures obtainable via low-cost devices. This capability allows for the unmasking of “hidden disease severity”—identifying patients who appear clinically stable in body sway (***z***-space) but are precariously close to an attractor bifurcation tipping point in parameter space (***μ***-space). *Personalized rehabilitation and optimization of interventions*: Distinguishing whether small-amplitude phenotypes (***ζ***_7_) stem from normal intermittent control with low noise (***μ***_2_) or from forced high-gain feedback compensating for broken intermittency (***μ***_6_) fundamentally transforms therapeutic strategies. For instance, prescribing conventional balance or strength training to patients exhibiting excessive continuous control (***μ***_6_ or ***μ***_8_) may prove counterproductive. Instead, interventions aimed at alleviating excessive co-contraction to re-acquire intermittent control flexibility within the OFF region (such as sensory feedback therapy or targeted pharmacological adjustment) are required. Furthermore, trajectory tracking within this latent parameter space offers a quantitative dynamic biomarker for evaluating therapeutic interventions, such as optimizing levodopa dosage or tuning Deep Brain Stimulation (DBS) parameters to actively steer the neuromuscular system away from unstable bifurcation boundaries.

### Model limitations and future directions

Several limitations of this study warrant consideration, yet they simultaneously underscore the structural robustness of our computational framework.

#### Simplification of the inverted pendulum model

Our digital twin model adopts a single-joint inverted pendulum in the sagittal plane (AP direction). Rather than a limitation, the fact that a minimal 1-joint formulation successfully captures the catastrophic bifurcation dynamics across over 1,000 clinical profiles demonstrates the structural robustness of the underlying attractor physics—suggesting that the primary mechanism of parkinsonian postural collapse is governed by a fundamental loss of low-dimensional manifold stability. Extension to multi-joint (ankle-hip (33, 34)) and multi-directional (AP-ML (35–37)) models remains a valuable future direction.

#### Cross-sectional study design

While the disease progression trajectory (***μ***(*s*)) presented here is derived from cross-sectional severity gradients across clusters, the continuous folded manifold mapped in ***μ***-space strongly implies a deterministic disease path. Future longitudinal studies tracking individual patients over several years across this latent space will directly validate the predictive timeline of these tipping points.

#### Direct correspondence with neurocircuitry

While identified parameters (*ρ*, Δ, *σ*, etc.) reflect functional control characteristics, establishing direct correspondences with specific neural activities—such as vestibular feedback control (38, 39), basal ganglia, cerebellar (39, 40), and spinal circuitry—will require integration with physiological modalities such as EMG, EEG, and fMRI.

### Conclusion

In conclusion, by unifying data assimilation with dynamical systems theory, this study elucidated the transformation of neural control strategies and phenotypic degeneracy in Parkinson’s disease. Reconstructing latent control mechanisms from observable sway data substantially advances our mechanistic understanding of movement disorders and paves a new path toward clinical digital-twin applications in precision medicine.

## Materials and Methods

### Participants

In this study, postural sway data were collected and analyzed from patients with Parkinson’s disease (PD) and healthy elderly individuals. The PD group comprised 140 participants (61 males, 79 females; mean age 70.5 *±* 7.7 years; total UPDRS-III score 23.8 *±* 7.8). The control group consisting of healthy elderly individuals comprised 59 participants (29 males, 30 females; mean age 70.0 *±* 3.7 years). Patient participants were recruited using a nonprobability convenience sampling approach. To capture a wide range of disease severity, investigators approached patients without filtering by clinical severity; however, the ability to maintain a standing posture without assistance was required as an inclusion criterion. This study was conducted with the approval of the Ethics Committees of the National Hospital Organization Osaka Toneyama Medical Center and the Graduate School of Engineering Science, Osaka University, and written informed consent was obtained from all participants.

Postural sway was measured using a Wii Balance Board (Nintendo, Kyoto) (29, 41). Participants were instructed to maintain a quiet standing posture in a relaxed state with eyes open, arms hanging naturally at their sides, feet positioned shoulder-width apart, and gaze fixed on a target placed 2 meters ahead. For each participant, a 60-second time series of the center of pressure (CoP) was recorded once at a sampling frequency of 100 Hz. Because the postural control model adopted in this study describes the movement dynamics of the body center of mass (CoM), the measured CoP time-series data were converted into CoM time-series data using the filter proposed by Morasso et al. (19). The resulting CoM data were smoothed using a 4th-order zero-phase Butterworth filter (cutoff frequency 10 Hz). To eliminate the effects of slow postural drift (low-frequency trends), linear detrending was applied to all CoM data prior to analysis.

The severity of motor impairment in PD patients was evaluated using UPDRS part III (3). The UPDRS-III consists of multiple examination items, each scored on a 5-point scale from 0 (normal) to 4 (severe), with higher scores indicating greater severity. These scores were determined based on clinical examinations and motor tests by experienced movement disorder specialists. In this study, the following five clinical subscores derived from items 18 to 31 of UPDRS-III were used as primary outcome measures:

- Total (total score for items 18–31, maximum 108 points)
- Tremor (sum of items 20 and 21, maximum 28 points)
- Rigidity (sum of item 22, maximum 20 points)
- Bradykinesia (sum of items 23–26 and 31, maximum 36 points)
- PIGD (postural instability and gait difficulty: sum of items 27–30, maximum 16 points)

### Intermittent control model of upright posture

To describe anterior-posterior (AP) postural sway during quiet standing, an intermittent control model was adopted (20). This model is a state-dependent switched hybrid system selected and driven by Eqs. [1] and [2].

In the equation of motion for the pendulum when neural feedback is inactive (OFF region, Eq. [2]), the equilibrium point (origin) is an unstable saddle point. On the state plane, this is described by a stable manifold (along which state trajectories approach the origin) and an unstable manifold (along which trajectories diverge from the origin). Conversely, when neural feedback is active (ON region, Eq. [1]), the dynamical character of the origin depends on the proportional (*P*) and derivative (*D*) gains. In typical intermittent control reproducing healthy postural sway, *P* and *D* are set to relatively small values, rendering the origin an unstable focus. This forms an unstable vector field that spirals outward due to transmission delays. By dynamically switching between these two unstable subsystems based on the state of the pendulum, intermittent control transiently exploits the stable manifold of the OFF state, achieving bounded dynamical stability. In the deterministic limit, this gives rise to limit-cycle oscillations. The OFF region is determined by the switching boundary lines of *θ* = 0 and the line with a slope *l* (Fig. 2, right). The intermittency ratio *ρ*, one of the most important parameter of the model is then expreseed as *ρ* = 0.5 *−* arctan(*l*)*/π*. In contrast, conventional “continuous” control models apply large *P* and *D* gains to directly stabilize the origin of the “ON” state (Eq. [1]) itself, causing the state to asymptotically converge to the equilibrium point without switching control OFF.

Numerical simulations were performed using the Euler–Maruyama scheme for stochastic differential equations (time step *dt* = 0.001 s) (20). The initial state (*θ*_0_, *ω*_0_) was randomly chosen within the range [−0.01, 0.01]. Simulations were run for 70 seconds for data assimilation; after discarding the initial 10 seconds as a transient period, the remaining 60 seconds were subjected to analysis. If the pendulum angle *θ* exceeded *π*/4 ≃ 0.7854 rad, the trial was deemed a fall, and the computation was forcibly terminated. The simulated time series were downsampled to 100 Hz and smoothed using the same 4th-order Butterworth filter applied to the empirical data.

The model parameters ***μ*** were identified such that the model-generated data accurately and statistically reproduced the experimental CoM data. The parameter space targeted for identification comprised the normalized proportional gain *P* ∈ [0, 500], derivative gain *D* ∈ [0, 500], active control frequency *ρ* ∈ [0.3, 1], transmission delay Δ ∈ [0, 0.5], noise intensity *σ* ∈ [0, 3], and deadzone radius *r*∈ [0, 0.01]. Subject-specific fixed parameters included mass *m* [kg] (subject’s body weight), height *h* = 1 [m], gravitational acceleration *g* = 9.81 [m/s^2^], and moment of inertia *I* = *mh*^2^. The passive stiffness and damping coefficients of the ankle joint were fixed at *K* = 0.8*mgh* [N m/rad] and *B* = 4.0 [m s/rad], respectively, based on prior literature (42, 43).

### Data assimilation

For model parameter estimation, Approximate Bayesian Computation based on Sequential Monte Carlo sampling (ABC-SMC) was constructed and applied (25, 27–29). This method identifies the posterior distribution of ***μ*** such that summary statistics Φ_sim_ computed from model-synthesized data statistically match the summary statistics Φ_obs_ obtained from observed data:

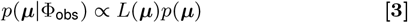

where *p*(***μ***) denotes the prior distribution (a uniform distribution in this study), and *p*(***μ*** Φ_obs_) represents the target posterior distribution. Instead of evaluating the explicit likelihood function *L*(***μ***), the ABC approach directly estimates *p*(***μ*** Φ_obs_) based on goodness-of-fit with summary statistics Φ_obs_. Simulations were executed using candidate parameters ***μ***_*i*_ sampled at the *i*-th iteration, and a parameter set was accepted only if the “statistical distance” between Φ_sim_ and Φ_obs_ fell below a tolerance threshold *ϵ*. This procedure was repeated until *N* = 500 valid samples were converged and retained. The threshold *ϵ* was updated and reduced in a stepwise manner by the SMC algorithm. Detailed algorithmic specifics of the sampling procedure are provided in SI Appendix A.

The summary statistic vector Φ (75 dimensions) consisted of normalized histograms (15 *×* 3 dimensions) of kinematic CoM variables (position, velocity, and acceleration) and their power spectral densities (10 *×* 3 dimensions). Each histogram was normalized so that the total probability equaled 1. Power spectral densities were calculated via FFT and normalized across 10 frequency bands on log-log axes up to 1.5 Hz.

The information-theoretic distance between Φ_obs_ and Φ_sim_ was evaluated using the Jensen–Shannon divergence *D*_JS_ (44):

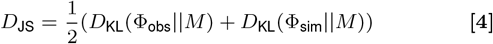

where 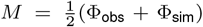, and *D*_KL_ denotes the Kullback–Leibler divergence. The threshold was initiated at *ϵ* = 1 with a final target set to *ϵ* = 0.01; however, considering the trade-off with computational cost, *ϵ* = 0.03 was applied as a practical convergence criterion.

### Synthetic data generation

For each of the 173 participants, parameters ***μ*** were sampled from their posterior distributions, generating synthetic data that satisfied the condition *D*_JS_ ≤ *ϵ*. Five independent sample paths (simulation trials) were generated for each participant and integrated with their single empirical CoM time series, constructing a hybrid digital-twin dataset comprising a total of 1,038 data points (173 empirical data + 865 synthetic data).

### Cluster analysis in model parameter *μ*-space

In the 5-dimensional control parameter space ***μ*** = (*p* = *P/mgh, D, ρ*, Δ, *σ*), hierarchical cluster analysis was performed on the constructed hybrid dataset. Note that the deadzone radius *r* was excluded from analysis as an explanatory variable due to its exceptionally high intra-individual variance in the posterior distribution. Ward’s method (45) was used for inter-cluster distance calculations. The optimal number of clusters was determined to be *k* = 3, comprehensively taking into account the Upper Tail method criterion (46) and the qualitative dynamics of generated sway trajectories. Clusters identified in this space are defined as “***μ***-clusters.” To evaluate differences in clinical motor scores across clusters, Kruskal–Wallis tests and post-hoc Mann–Whitney U tests with Bonferroni correction for multiple comparisons were applied. Effect sizes between groups were computed using Cliff’s *δ* to quantify practical magnitude.

### Postural sway indices

From each CoM time series, a total of 18 standard postural sway indices were calculated, including mean-crossing rate (Mean-cross), root mean square (RMS), mean velocity (MV), and scaling exponents derived from stabilized detrended fluctuation analysis (SDA) and detrended fluctuation analysis (DFA) (47–49). Formulations and details for each index are documented in SI Appendix B. Each calculated index was smoothed using Box–Cox transformation to ensure normality. To preserve biological variability and unique dynamics intrinsic to individual postural control mechanisms, no outlier removal was performed.

### Cluster analysis in latent variable *z*-space

Hierarchical cluster analysis with Ward’s method was performed on data points in the ***z***-space extracted via factor analysis. To enable direct topological comparisons with ***μ***-clusters, the number of clusters was fixed at 10. These are defined as “***ζ***-clusters.” The identical statistical analysis pipeline used for ***μ***-clusters was applied to compare and test clinical scores across ***ζ***-clusters.

### Learning the mapping functions between *μ* and *z*

To quantitatively identify bidirectional mapping relationships between the two parametric spaces and linguistic spaces, a fully connected 5-layer deep neural network (NN) was constructed to learn forward mapping ***μ*** ↦ ***z*** and inverse mapping ***z*** ↦ ***μ***. The dataset was randomly shuffled and partitioned into a training set (80 %), a validation set (10 %), and a test set (10 %). The NN architecture comprised an input layer (6 neurons), three hidden layers (128, 128, and 64 neurons, respectively, with Sigmoid activation functions), and an output layer (6 neurons with linear activation functions). The Adam optimizer (3000 epochs, early stopping applied) was employed for optimization. To prevent model overfitting, an *L*_2_ regularization term (*λ* = 1.0 *×* 10^*−*5^) was incorporated into the loss function. Model generalization performance and inference accuracy were validated using 9-fold cross-validation and geometric comparison of inferred cluster centroids.

### Correspondence table via surrogate data

To construct transition probability and mapping correspondence tables between ***μ***-space and ***z***-space, 10,000 surrogate data points per cluster were generated from multivariate normal distributions based on each cluster’s centroid and covariance matrix. These were input into the trained NNs to identify which target cluster centroid in the destination space the predicted output vector belonged to with minimum distance in the sense of the *L*_2_ norm.

### Bifurcation Diagrams along *μ* Transitions

To dynamically analyze transformations in attractor structure accompanying disease progression, a linear trajectory connecting two contrasting cluster centroids 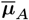 and 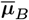 was defined in ***μ***-space, standardized across each dimension:

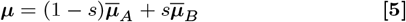

The parameter *s* was continuously varied over the range from *−* 1 to 2. For each parameter point ***μ***(*s*), both deterministic (*σ* = 0) and stochastic simulations were executed. Bifurcation diagrams were constructed using stroboscopic mapping, continuously recording pendulum angle *θ* values at the “ON/OFF” control switching boundary as Poincaré sections. Furthermore, to evaluate distribution characteristics of stochastic trajectories at each *s*, the 2.5th and 97.5th percentile values of sampled points were computed. To rigorously quantify contraction and expansion trends of attractor distributions relative to variations in control intermittency *ρ* and noise intensity *σ*, Pearson correlation coefficients were calculated between *s* and the 2.5th and 97.5th percentile values of stochastic plots. Additionally, individual data points constituting clusters A and B were orthogonally projected onto line [5] using the following equation:

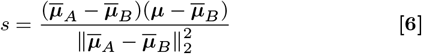

where ∥·∥_2_ denotes the *L*_2_ norm. To visualize the relative distribution density of data points projected onto the *s*-axis, histograms were plotted along the *s*-axis with the vertical axis representing probability density.

## Data, Materials, and Software Availability

All data supporting the findings of this study, as well as the custom computational code and data assimilation programs developed for the mechanistic digital-twin modeling and neural network inverse mapping, are available from the corresponding author upon reasonable request. The empirical postural sway time-series data from human participants are not publicly deposited due to ethical restrictions and the terms of informed consent regarding third-party data sharing.

## ACKNOWLEDGMENTS

This study was supported by the JSPS KAKENHI, No. 22H03662 (T.N.) and Suzuken Memorial Foundation (T.N.).

## References

1. FB Horak, Postural orientation and equilibrium: what do we need to know about neural control of balance to prevent falls? Age Ageing 35, ii7–ii11 (2006).

2. Y Ivanenko, VS Gurfinkel, Human postural control. Front. Neurosci. 12, 171 (2018).

3. S Fahn, RL Elton, Unified Parkinson’s disease rating scale in Recent Developments in Parkinson’s Disease. (Macmillan Healthcare Information) Vol. 2, pp. 153–163 (1987).

4. H Braak, et al., Staging of brain pathology related to sporadic Parkinson’s disease. Neurobiol. Aging 24, 197–211 (2003).

5. JM Fearnley, AJ Lees, Ageing and Parkinson’s disease: substantia nigra regional selectivity. Brain 114, 2283–2301 (1991).

6. MM Hoehn, MD Yahr, Parkinsonism: onset, progression, and mortality. Neurology 17, 427–427 (1967).

7. BR Bloem, JM Hausdorff, JE Visser, N Giladi, Falls and freezing of gait in Parkinson’s disease: a review of two interconnected, episodic phenomena. Mov. Disord. 19, 871–884 (2004).

8. CG Goetz, et al., Movement Disorder Society Task Force report on the Hoehn and Yahr staging scale: status and recommendations. Mov. Disord. 19, 1020–1028 (2004).

9. FM Skidmore, et al., The emerging postural instability phenotype in idiopathic Parkinson disease. npj Park. Dis. 8, 28 (2022).

10. FB Horak, JG Nutt, LM Nashner, Postural inflexibility in parkinsonian subjects. J. Neurol. Sci. 111, 46–58 (1992).

11. T Yamamoto, et al., A classification of postural sway patterns during upright stance in healthy adults and patients with parkinson’s disease. J. Adv. Comput. Intell. Intell. Informatics 15, 997–1010 (2011).

12. P Panyakaew, C Anan, R Bhidayasiri, Visual deprivation elicits subclinical postural inflexibilities in early parkinson’s disease. J. Neurol. Sci. 349, 214–219 (2015).

13. T Perera, et al., Balance control systems in parkinson’s disease and the impact of pedunculopontine area stimulation. Brain 141, 3009–3022 (2018).

14. C Maurer, RJ Peterka, A new interpretation of spontaneous sway measures based on a simple model of human postural control. J. Neurophysiol. 93, 189–200 (2005).

15. S Fujii, Y Takamura, K Ikuno, S Morioka, N Kawashima, A comprehensive multivariate analysis of the center of pressure during quiet standing in patients with parkinson’s disease. J. NeuroEngineering Rehabil. 21, 59 (2024).

16. A Julienne, E Verbecque, S Besnard, Normative data for instrumented posturography: a systematic review and meta-analysis. Front. Hum. Neurosci. 18, 1498107 (2024).

17. W Ge, CJ Lueck, D Apthorp, H Suominen, Which features of postural sway are effective in distinguishing parkinson’s disease from controls? a systematic review. Brain Behav. 11, e01929 (2021).

18. A Kamieniarz, et al., A posturographic procedure assessing balance disorders in parkinson’s disease: a systematic review. Clin. Interv. Aging 13, 2301–2316 (2018).

19. PG Morasso, G Spada, R Capra, Computing the COM from the COP in postural sway movements. Hum. Mov. Sci. 18, 759–767 (1999).

20. Y Asai, et al., A model of postural control in quiet standing: robust compensation of delay-induced instability using intermittent activation of feedback control. PLoS ONE 4, e6169 (2009).

21. ID Loram, H Gollee, M Lakie, PJ Gawthrop, Human control of an inverted pendulum: is continuous control necessary? is intermittent control effective? is intermittent control physiological? The J. Physiol. 589, 307–324 (2011).

22. T Nomura, Y Suzuki, PG Morasso, Intermittent control strategy for stabilizing human quiet stance, a model of the in Encyclopedia of Computational Neuroscience. (Springer), pp. 1694–1704 (2022).

23. JG Milton, T Insperger, W Cook, DM Harris, G Stepan, Acting together, destabilizing influences can stabilize human balance. Philos. Transactions Royal Soc. A: Math. Phys. Eng. Sci. 377, 20190044 (2019).

24. T Takazawa, et al., How the brain can be trained to achieve an intermittent control strategy for stabilizing quiet stance by means of reinforcement learning. Biol. Cybern. 118, 229–248 (2024).

25. Y Suzuki, et al., Postural instability via a loss of intermittent control in elderly and patients with parkinson’s disease: A model-based and data-driven approach. Chaos 30, 113140 (2020).

26. J Bélair, L Glass, U an der Heiden, J Milton, Dynamical disease: identification, temporal aspects and treatment strategies of human illness. Chaos: An Interdiscip. J. Nonlinear Sci. 5, 1–7 (1995).

27. T Toni, D Welch, N Strelkowa, A Ipsen, MPH Stumpf, Approximate Bayesian computation scheme for parameter inference and model selection in dynamical systems. J. Royal Soc. Interface 6, 187–202 (2009).

28. J Lintusaari, MU Gutmann, R Dutta, S Kaski, J Corander, Fundamentals and recent developments in approximate Bayesian computation. Syst. Biol. 66, e66–e82 (2017).

29. A Tietä vä inen MU Gutmann, E Keski-Vakkuri, J Corander, E Hæggström Bayesian inference of physiologically meaningful parameters from body sway measurements. Sci. Reports 7, 3771 (2017).

30. A Julienne, E Verbecque, S Besnard, Normative data for instrumented posturography: a systematic review and meta-analysis. Front. Hum. Neurosci. 18, 1498107 (2024).

31. B van Breugel, T Liu, D Oglic, M van der Schaar, Synthetic data in biomedicine via generative artificial intelligence. Nat. Rev. Bioeng. 2, 991–1004 (2024).

32. A Koul, D Duran, T Hernandez-Boussard, Synthetic data, synthetic trust: navigating data challenges in the digital revolution. The Lancet Digit. Heal. 7, 11 (2025).

33. T Kato, et al., Anti-phase action between the angular accelerations of trunk and leg is reduced in the elderly. Gait & Posture 40, 107–112 (2014).

34. Y Suzuki, T Nomura, M Casadio, PG Morasso, Intermittent control with ankle, hip, and mixed strategies during quiet standing: A theoretical proposal based on a double inverted pendulum model. J. Theor. Biol. 310, 55–79 (2012).

35. T Exley, S Moudy, RM Patterson, J Kim, MV Albert, Predicting UPDRS motor symptoms in individuals with Parkinson’s disease from force plates using machine learning. IEEE J. Biomed. Heal. Informatics 26, 3486–3494 (2022).

36. M Mangalam, et al., Older adults and individuals with Parkinson’s disease control posture along suborthogonal directions that deviate from the traditional anteroposterior and mediolateral directions. Sci. Reports 14, 4117 (2024).

37. D Engel, et al., Sway frequencies may predict postural instability in Parkinson’s disease: a novel convolutional neural network approach. J. NeuroEngineering Rehabil. 22, 29 (2025).

38. RJ Peterka, Sensorimotor integration in human postural control. J. Neurophysiol. 88, 1097–1118 (2002).

39. RJ Peterka, KD Statler, DM Wrisley, FB Horak, Postural compensation for unilateral vestibular loss. Front. Neurol. 2, 57 (2011).

40. JM Goldberg, et al., The Vestibular System: A Sixth Sense. (Oxford University Press), (2012).

41. RA Clark, et al., Validity and reliability of the Nintendo Wii balance board for assessment of standing balance. Gait & Posture 31, 307–310 (2010).

42. ID Loram, M Lakie, Direct measurement of human ankle stiffness during quiet standing: the intrinsic mechanical stiffness is insufficient for stability. The J. Physiol. 545, 1041–1053 (2002).

43. M Casadio, PG Morasso, V Sanguineti, Direct measurement of ankle stiffness during quiet standing: implications for control modelling and clinical application. Gait & Posture 21, 410–424 (2005).

44. J Vanlier, CA Tiemann, PA Hilbers, NA van Riel, Optimal experiment design for model selection in biochemical networks. BMC Syst. Biol. 8, 1–16 (2014).

45. JHJ Ward, Hierarchical grouping to optimize an objective function. J. Am. Stat. Assoc. 58, 236–244 (1963).

46. R Mojena, Hierarchical grouping methods and stopping rules: an evaluation. The Comput. J. 20, 359–363 (1977).

47. T Yamamoto, et al., Universal and individual characteristics of postural sway during quiet standing in healthy young adults. Physiol. Reports 3, e12329 (2015).

48. M Duarte, VM Zatsiorsky, Long-range correlations in human standing. Phys. Lett. A 283, 124–128 (2001).

49. TE Prieto, JB Myklebust, RG Hoffmann, EG Lovett, AM Myklebust, Measures of postural steadiness: differences between healthy young and elderly adults. IEEE Transactions on Biomed. Eng. 43, 956–966 (1996).

