## Supporting Information for "A Dynamical Digital Twin Unmasks Hidden Neuromotor Control Policies and Catastrophic Tipping Points in Parkinson’s Disease"

**This PDF file includes:**

Figs. S1 to S15
Tables S1 to S25

### A Generation of parameter candidates from posterior distribution

In this study, we employed Approximate Bayesian Computation based on Sequential Monte Carlo sampling (ABC-SMC) for data assimilation. ABC-SMC is a method for inferring multiple model parameters that parameterize stochastic dynamic nonlinear systems based on Bayesian inference. It estimates the Bayesian posterior distribution of model parameters $\boldsymbol{\mu}$ such that the summary statistics $\Phi_{\mathrm{sim}}$ of the synthetic time-series data generated by the model are sufficiently similar to the summary statistics $\Phi_{\mathrm{obs}}$ of the observed time-series data (real data), based on the likelihood function $L(\boldsymbol{\mu})$ using the following equation:

$$p(\boldsymbol{\mu}|\Phi_{\mathrm{obs}}) \propto L(\boldsymbol{\mu})p(\boldsymbol{\mu})$$

where $p(\boldsymbol{\mu})$ is the prior probability distribution of the parameters $\boldsymbol{\mu}$ (a uniform distribution in this study), and $p(\boldsymbol{\mu}|\Phi_{\mathrm{obs}})$ is the posterior probability distribution given the observed data $\Phi_{\mathrm{obs}}$. The posterior distribution approaches the true parameter distribution by conditioning on the observed data. In ABC, $p(\boldsymbol{\mu}|\Phi_{\mathrm{obs}})$ is estimated based on summary statistics $\Phi_{\mathrm{obs}}$ calculated from the observed data instead of using $L(\boldsymbol{\mu})$. First, parameter candidates $\boldsymbol{\mu}_i$ are randomly generated according to $p(\boldsymbol{\mu})$, and simulations are performed using these generated $\boldsymbol{\mu}_i$.

Next, the summary statistics $\Phi_{\mathrm{sim}}$ of the resulting simulation data are calculated. If the value representing the "distance" between the summary statistics of the observed data $\Phi_{\mathrm{obs}}$ and those of the simulation data $\Phi_{\mathrm{sim}}$ is smaller than a certain threshold $\epsilon$, the parameter candidate $\boldsymbol{\mu}_i$ is accepted and used as an element to construct the posterior probability distribution. This trial is repeated until the number of accepted samples reaches $N \in \mathbb{N}$, and the posterior distribution $p(\boldsymbol{\mu}|\Phi_{\mathrm{obs}})$ is estimated from the set of accepted parameters. When performing ABC based on SMC, the bias of importance sampling for the obtained posterior distribution $p(\boldsymbol{\mu}|\Phi_{\mathrm{obs}})$ is corrected to serve as the prior distribution $p(\boldsymbol{\mu})$ for the next step. The ABC process is then repeated while stepwise decreasing the distance threshold $\epsilon$. The algorithm for generating parameter candidates $\boldsymbol{\mu}_i$ from the prior distribution $p(\boldsymbol{\mu})$ is described as follows.

In this study, the first round of the algorithm begins with the initial distance threshold $\epsilon = 1$. The range of uniform distribution for each of the six components of $\boldsymbol{\mu}$ is identical to the exploration range of estimated parameters (proportional gain $P \in [0, 500]$; derivative gain $D \in [0, 500]$; intermittency of active control $\rho \in [0.3, 1]$; feedback time delay $\Delta \in [0, 0.5]$; noise intensity $\sigma \in [0, 3]$; radius of sensory dead zone $r \in [0, 0.01]$).

A parameter candidate $\boldsymbol{\mu}_i$ is generated from the uniform distribution, and $\boldsymbol{\mu}_i$ will be accepted when the Jenssen-Shannon divergence $D_{\mathrm{JS}}$ between the real CoM data and the simulated CoM data with $\boldsymbol{\mu}_i$ is less than 1 (Eq. 3 in the main text).

Generation of parameter candidates is repeated until the number of accepted parameters $N$ reaches 500. For each accepted parameter $\boldsymbol{\mu}_i$, we assigned a weight parameter as $w_i = 1/N$.

From the second round onwards, we use a gradually reduced value of $\epsilon < 1$. We used the set of accepted parameter values $M^{\mathrm{pre}}$ collected in the previous loop for preparing the prior distribution for the second round onwards.

$$M^{\mathrm{pre}} = \begin{pmatrix} P_1^{\mathrm{pre}} & D_1^{\mathrm{pre}} & \rho_1^{\mathrm{pre}} & \Delta_1^{\mathrm{pre}} & \sigma_1^{\mathrm{pre}} & r_1^{\mathrm{pre}} \\ P_2^{\mathrm{pre}} & D_2^{\mathrm{pre}} & \rho_2^{\mathrm{pre}} & \Delta_2^{\mathrm{pre}} & \sigma_2^{\mathrm{pre}} & r_2^{\mathrm{pre}} \\ \vdots & \vdots & \vdots & \vdots & \vdots & \vdots \\ P_N^{\mathrm{pre}} & D_N^{\mathrm{pre}} & \rho_N^{\mathrm{pre}} & \Delta_N^{\mathrm{pre}} & \sigma_N^{\mathrm{pre}} & r_N^{\mathrm{pre}} \end{pmatrix}$$

Overall, to determine the index number $n_{\mathrm{core}} \in [1, 2, \ldots, N]$, which is used to determine which parameter candidate of $M^{\mathrm{pre}}$ accepted in the previous round should be selected as the proposed parameter this time, $n_{\mathrm{core}}$ is sampled from the discrete distribution with probabilities proportional to the normalized weights $\boldsymbol{w}$ according to the uniform distribution prior. After determining $n_{\mathrm{core}}$ as described later, taking $\boldsymbol{\mu}^{\mathrm{pre}}_{n_{\mathrm{core}}}$ as the core parameter, a parameter candidate $\boldsymbol{\mu}_i$ for $i$-th sampling is generated by adding fluctuations while preserving the correlation structure of $M^{\mathrm{pre}}$.

Specifically, let the standard deviations of each column of $M^{\mathrm{pre}}$ be denoted by $s_P^{\mathrm{pre}}$, $s_D^{\mathrm{pre}}$, $s_\rho^{\mathrm{pre}}$, $s_\Delta^{\mathrm{pre}}$, $s_\sigma^{\mathrm{pre}}$, and $s_r^{\mathrm{pre}}$, respectively.

Let $\Sigma$ be the variance-covariance matrix of the column-wise normalized $M^{\mathrm{pre}}$. Since $\Sigma$ is a symmetric positive-definite matrix, the following Cholesky decomposition of $2\Sigma$ is possible:

$$2\Sigma = HH^{\mathrm{T}}$$

where $H$ is a lower triangular matrix. The factor of 2 applied to $\Sigma$ is introduced to widen the range of fluctuations added to $\boldsymbol{\mu}^{\mathrm{pre}}_{n_{\mathrm{core}}}$. Furthermore, from the cumulative sum of weight parameter $\boldsymbol{w}^{\mathrm{pre}}$ for the previous round, $w_i^{\mathrm{cum}}$ is computed as

$$w_i^{\mathrm{cum}} = \sum_{j=1}^{i} w_j^{\mathrm{pre}}.$$

Then, the iterative procedure for generating parameter candidates is as follows: First, based on $\boldsymbol{w}^{\mathrm{cum}}$ and a random number $z_r$ drawn from a uniform distribution over $[0, 1]$, $n_{\mathrm{core}}$ is determined as

$$n_{\mathrm{core}} = \min\left\{i | w_i^{\mathrm{cum}} \geq z_r\right\}.$$

This implies that $\boldsymbol{\mu}_i^{\mathrm{pre}}$ corresponding to larger weight parameters $w_i^{\mathrm{pre}}$ are more likely to be selected as the core. Next, six random numbers (corresponding to the number of model parameters to be estimated) are generated from the standard normal distribution to construct the standard normal vector $\boldsymbol{a} = (a_1, a_2, a_3, a_4, a_5, a_6)^{\mathrm{T}}$. Each element of $\boldsymbol{a}$ is mutually independent and follows $N(0,1)$; therefore, $\boldsymbol{a}$ is a single sample from the six-dimensional standard normal distribution $N(\mathbf{0},\boldsymbol{I})$. From $\boldsymbol{a}$ and $H$, we define

$$\boldsymbol{z} = H\boldsymbol{a}.$$

Then, we have the following equations:

$$\mathrm{Cov}(\boldsymbol{z}) = E\left[\left(\boldsymbol{z} - E\left[\boldsymbol{z}\right]\right)\left(\boldsymbol{z} - E\left[\boldsymbol{z}\right]\right)^{\mathrm{T}}\right] = 2\Sigma.$$

That is, $\boldsymbol{z}$ is a random vector that has the (normalized) correlation structure of $M^{\mathrm{pre}}$. Using the matrix product of $\boldsymbol{z}$ and the sample standard deviation vector $\boldsymbol{s}^{\mathrm{T}} = \left(s_P^{\mathrm{pre}}, s_D^{\mathrm{pre}}, s_\rho^{\mathrm{pre}}, s_\Delta^{\mathrm{pre}}, s_\sigma^{\mathrm{pre}}, s_r^{\mathrm{pre}}\right)$ as the fluctuation term, the parameter candidate $\boldsymbol{\mu}_i$ is generated as

$$\boldsymbol{\mu}_i = \boldsymbol{\mu}_{n_{\mathrm{core}}}^{\mathrm{pre}} + \boldsymbol{z}\boldsymbol{s}^{\mathrm{T}}.$$

Multiplication by the sample standard deviations restores the scaling of the variance-covariance matrix $2\Sigma$, followed by $\boldsymbol{z}$, to that of the original $M^{\mathrm{pre}}$. The generated parameter candidate $\boldsymbol{\mu}_i$ is then evaluated using $D_{\mathrm{JS}}$ (Eq. 3 in the main text), and the candidate generation process is repeated until the number of accepted samples reaches $N = 500$.

For each accepted $\boldsymbol{\mu}_i$, the weight parameter $w_i$ is assigned as

$$w_i = \frac{1}{\sum_{j=1}^{N} w_j^{\mathrm{pre}} f_j\left(\boldsymbol{\mu}_i \middle| \boldsymbol{\mu}_j^{\mathrm{pre}}, \Sigma\right)},$$

where, $f_j\left(\boldsymbol{x}|\boldsymbol{\mu}_j^{\mathrm{pre}}, \Sigma\right)$ denotes the probability density function of the six-dimensional normal distribution with mean vector $\boldsymbol{\mu}_j^{\mathrm{pre}}$ and covariance matrix $\Sigma$, given by

$$f_j\left(\boldsymbol{x}|\boldsymbol{\mu}_j^{\mathrm{pre}}, \Sigma\right) = \frac{1}{\sqrt{(2\pi)^6|\Sigma|}} \exp\left(-\frac{(\boldsymbol{x}-\boldsymbol{\mu}_j^{\mathrm{pre}})^{\mathrm{T}}\Sigma^{-1}(\boldsymbol{x}-\boldsymbol{\mu}_j^{\mathrm{pre}})}{2}\right).$$

A larger value of $f_j\left(\boldsymbol{\mu}_i|\boldsymbol{\mu}_j^{\mathrm{pre}}, \Sigma\right)$ indicates a higher likelihood that the accepted $\boldsymbol{\mu}_i$ was sampled using $\boldsymbol{\mu}_j^{\mathrm{pre}}$ as the core (important sampling). However, since $w_i$ is defined as the reciprocal of the weighted sum of $f_j\left(\boldsymbol{\mu}_i|\boldsymbol{\mu}_j^{\mathrm{pre}}, \Sigma\right)$, larger weights are assigned to those accepted $\boldsymbol{\mu}_i$ that are less likely to have been sampled from the parameter set $M^{\mathrm{pre}}$ (the bias of importance sampling for the obtained posterior distribution $p(\boldsymbol{\mu}|\Phi_{\mathrm{obs}})$ is corrected, according the uniform distribution prior).

After the number of accepted samples reaches $N = 500$, each $w_i$ is divided by $\sum_{i=1}^{N} w_i$ so that the total sum of the weights is normalized to 1.

### B Postural sway indices calculation

In this study, we analyzed time series data of the Center of Mass (CoM) in the anteroposterior direction. The data length was 60 s, and the sampling frequency was 100 Hz. As a preprocessing, low-pass filtering was applied to the CoM data using a fourth order zero lag Butterworth filter with a cutoff frequency of 10 Hz. Then, for real data, linear trend was removed from the CoM data. Here, we denote the preprocessed CoM data at time $n$ as $\mathrm{CoM}[n]$, and the corresponding data length is expressed as $N = 60 \times 100 = 6000$. In the following sections, we present the definitions of 18 sway indices used in this study. Most indices are calculated using MATLAB R2024b.

**(1) Mean-cross:** Mean-cross is the number of times the CoM time series crosses its mean position. In this study, we calculated this as the number of sign changes in the CoM time series because the CoM data was linearly detrended.

**(2) 0-cross-V:** 0-cross-V is the number of zero-crossing events in the CoM velocity time series. This velocity time series was calculated using the central difference method, and then low-pass filtered at a cutoff frequency of 2.5 Hz using a fourth order zero lag Butterworth filter. As in (1), we defined 0-cross-V as the number of sign changes. The filtered velocity time series was also used in (3) and (4).

**(3), (4) Alpha-V and Beta-V:** Alpha-V and Beta-V are the shape and scale parameters of a gamma distribution, respectively. The probability density function of the gamma distribution is written as

$$p\left(x\right) = \frac{1}{\Gamma\left(\alpha\right)\beta^{\alpha}} x^{\alpha-1} e^{-\frac{x}{\beta}}$$

where $x$ is the time interval of 0-cross-V events, $\Gamma\left(k\right) = \int_0^{\infty} t^{k-1} e^{-t} dt$ is a gamma function.

$\alpha$ and $\beta$ in the above equation correspond to Alpha-V and Beta-V, respectively. To calculate these parameters, we constructed a histogram of the zero-crossing time intervals for the low-pass filtered CoM velocity time series. This histogram was normalized to have an area of 1, and then a gamma distribution was fitted to the histogram. This fitting was performed using the *fitdistr* function in R.

**(5) RMS:** RMS is a root mean square of the CoM time series, defined as

$$\mathrm{RMS} = \sqrt{\frac{1}{N}\sum_{n=1}^{N} \mathrm{CoM}[n]^2}.$$

**(6) MV:** MV is the mean velocity of the CoM time series, and is not an average of the velocity time series. MV is calculated as

$$\mathrm{MV} = \frac{1}{T}\sum_{n=1}^{N-1} \left|\mathrm{CoM}[n+1] - \mathrm{CoM}[n]\right|.$$

**(7) MFREQ:** MFREQ is an angular frequency of total path length converted to circular motion with radius of effective value for mean amplitude of the CoM time series. Using (6), MFREQ is defined as

$$\mathrm{MFREQ} = \frac{\mathrm{MV}}{4\sqrt{2}\sum_{n=1}^{N}\left|\mathrm{CoM}[n]\right|}.$$

**(8) Power:** Power is the total power of the CoM time series. In this study, we defined Power as the sum of Power Spectral Density (PSD) from 0.75 Hz to 5 Hz. We estimated the PSD of the CoM time series using the short term Fourier transformation with a Hamming window and Welch's method. The window size was 40 s and the overlap was 35 s. Thus, we divided the CoM time series into five segments (0–40, 5–45, 10–50, 15–55, and 20–60 s). The Hamming window is defined as

$$w[n] = 0.54 - 0.46\cos\frac{2\pi\left(n-1\right)}{40 \times f_s} \qquad \left(n = 1, 2, \ldots, 40 \times f_s\right)$$

where $f_s$ is the sampling frequency.

Using Welch's method, the PSD is calculated by the following equation:

$$\mathrm{PSD}[m] = \mathrm{mean}\left(\frac{\left|\mathcal{F}[\mathrm{CoM}_{40} \cdot w][m]\right|^2}{\sum w[n]^2}\right)$$

Where $\mathrm{CoM}_{40}$ is each CoM time series with data length of 40 s, $\mathcal{F}[\cdot]$ is the Fourier transformation. $m = 1, 2, \ldots, 4000$ are discrete frequencies corresponding to $0, 0.025, \ldots$,99.975 Hz, respectively.

Let $i$ and $j$ be the discrete frequencies corresponding to 0.75 Hz and 5 Hz, respectively, Power is defined as below.

$$\mathrm{Power} = \sum_{m=i}^{j} \mathrm{PSD}[m]$$

**(9), (10) PF50 and PF95:** PF50 and PF95 are frequencies where power of the CoM reaches 50 % and 95 % of the total power, respectively. PF50 is calculated as

$$\mathrm{PF50} = \min\left\{ u \,\middle|\, \sum_{m=i}^{u} \mathrm{PSD}[m] \geq 0.5 \times \mathrm{Power} \right\} \times \Delta f$$

where $\Delta f$ the an increment of frequency. In this study, $\Delta f = 0.025$ Hz because the window size was 40 s. Similarly, PF95 is calculated as

$$\mathrm{PF95} = \min\left\{ u \,\middle|\, \sum_{m=i}^{u} \mathrm{PSD}[m] \geq 0.95 \times \mathrm{Power} \right\} \times \Delta f.$$

**(11), (12) Slope-short and Slope-long:** We performed Stabilogram Diffusion Analysis (SDA). In SDA, we calculate the distance between CoM at time $t$ and at time $t + \Delta t$, then plot a time lag $\Delta t$ on x-axis and the squared mean difference $\Delta\mathrm{CoM}^2$ on y-axis. Let $\Delta n = \Delta t \times f_s$, $\Delta\mathrm{CoM}^2$ is calculated as

$$\Delta\mathrm{CoM}^2[\Delta n] = \frac{1}{N - \Delta n} \sum_{n=1}^{N-\Delta n} \left(\mathrm{CoM}[n] - \mathrm{CoM}[n + \Delta n]\right)^2 .$$

Slope-long and Slope-short are the scaling exponents at short-term and long-term regions, respectively, in log-log scale SDA plot. We describe how to calculate the approximate straight line in each region later.

**(13), (14) Critical-Δt and Critical-D:** Critical-Δt and Critical-D are the coordinates of an intersection point of two approximate straight lines in log-log scale SDA plot, i.e., the critical time lag (Critical-Δt) and the critical squared mean difference (Critical-D), respectively. These two indices are determined in the calculation process of the two approximate straight lines.

**(15), (16) Alpha-small and Alpha-large:** We also performed Detrended Fluctuation Analysis (DFA). In DFA, an integrated time series $Y[n]$ ($n = 1, 2, \ldots, N$) is constructed from the original CoM time series. $Y[n]$ is then divided into non-overlapping segments (boxes) of length $T_{\mathrm{bs}}$ (box size: bs), and the fluctuation relative to the local trend, $F[\mathrm{bs}]$, is calculated within each box ($\mathrm{bs} = T_{\mathrm{bs}} \times f_s$). $Y[n]$ and $F[\mathrm{bs}]$ are calculated as followings.

$$Y[n] = \sum_{i=1}^{n} \mathrm{CoM}[i]$$

$$F[\mathrm{bs}] = \sqrt{\frac{1}{\lfloor N/\mathrm{bs} \rfloor} \sum_{j=1}^{\lfloor N/\mathrm{bs} \rfloor} \left( \frac{1}{\mathrm{bs}} \sum_{k=(j-1)\mathrm{bs}+1}^{j\mathrm{bs}} \left( Y[k] - p_j^{(2)}[k] \right)^2 \right)}$$

where $\lfloor \cdot \rfloor$ is the floor function, and $p_j^{(m)}$ is the polynomial representing $m$-th order trend in $j$-th box. In this study, we performed second-order detrending. In addition, the values of bs were determined by rounding to integers a geometric sequence with an initial term of 4 and a common ratio of $10^{0.05}$. By adopting this scheme, the plotted points become evenly spaced (on a logarithmic scale), preventing an uneven concentration in the number of points across the range.

Alpha-small and Alpha-large are the scaling exponents at short-term and long-term regions, respectively, in log-log scale DFA plot. We also describe how to calculate the approximate straight line in each region later.

**(17), (18) Critical-bs and Critical-fn:** Critical-bs and Critical-fn are the coordinates of an intersection point of two approximate straight lines in log-log scale DFA plot, i.e., the critical box size (Critical-bs) and the critical fluctuation value (Critical-fn), respectively. These two indices are determined during the calculation process of the two approximate straight lines.

**Estimation of scaling exponents in SDA and DFA plots.** Scaling exponents in SDA and DFA analysis were estimated based on the approximate straight lines in the short-term and long-term regions of the SDA and DFA plots. Due to the large sample size in this study, it would be difficult to apply fixed ranges for the two target regions commonly across participants. Therefore, by assuming that the two target regions were contiguous, we explored possible boundaries between the two regions to minimize the fitting error of the two regression lines for each CoM data.

To perform the two-line approximation, we defined a boundary point A such that the first region extended up to point A and the second region began at point A. We then devised a two-line approximation in which the two lines intersect at point A, using the least-squares method.

For a simple linear regression over a given range, where $n$ data points exist, the least-squared error between the data points $(x_i, y_i)$ and the line is defined as

$$S = \sum_{i=1}^{n} \left(y_i - (a x_i + b)\right)^2$$

and the parameters $a$ and $b$ of the approximating line are obtained by using the condition

$$\frac{\partial S}{\partial a} = \frac{\partial S}{\partial b} = 0.$$

Specifically,

$$a = \frac{\mathrm{Cov}(x, y)}{\mathrm{Var}(x)},$$

$$b = \bar{y} - a\bar{x},$$

where $\mathrm{Cov}(x, y)$ denotes the covariance between $x$ and $y$, $\mathrm{Var}(x)$ denotes the variance of $x$, and $\bar{x}$ and $\bar{y}$ denote the mean values of $x$ and $y$, respectively.

Now consider a computational range $[x_{\mathrm{start}}, x_{\mathrm{end}}]$ divided into two subranges at a boundary point $x_A$. Suppose that the approximating line in the first range $[x_{\mathrm{start}}, x_A]$, given by $y = a_1 x + b_1$, is obtained using the least-squares method as described above. We then determine the approximating line for the second range $[x_A, x_{\mathrm{end}}]$, constrained to pass through the point $(x_A, a_1 x_A + b_1)$. The least-squares error in the second range is defined as

$$S_2 = \sum_{i=i_A}^{i_{\mathrm{end}}} \left(y_i - (a_2 x_i + b_2)\right)^2,$$

where $i_A$ and $i_{\mathrm{end}}$ denote the indices satisfying $x_{i_A} = x_A$ and $x_{i_{\mathrm{end}}} = x_{\mathrm{end}}$, respectively. Because the line $y = a_2 x + b_2$ passes through $(x_A, a_1 x_A + b_1)$, we substitute $b_2 = y - a_2 x = (a_1 x_A + b_1) - a_2 x_A$ into $S_2$, yielding

$$S_2 = \sum_{i=i_A}^{i_{\mathrm{end}}} \left((x_A - x_i)\, a_2 + y_i - a_1 x_A - b_1\right)^2.$$

The value of $a_2$ is obtained from the condition $\frac{dS_2}{da} = 0$, resulting in

$$a_2 = \frac{\sum_{i=i_A}^{i_{\mathrm{end}}} (y_i - a_1 x_A - b_1)(x_i - x_A)}{\sum_{i=i_A}^{i_{\mathrm{end}}} (x_i - x_A)^2},$$

$$b_2 = (a_1 - a_2)\, x_A + b_1.$$

In this study, for both SDA and DFA, the computational range was set from 0.2 s to 15 s. Since the coordinate axes are logarithmic in both directions, common logarithms were used for $x$ and $y$. Specifically, for SDA we defined $x = \log_{10} \Delta n$ and $y = \log_{10} \Delta\mathrm{CoM}^2[\Delta n]$; for DFA we defined $x = \log_{10} \mathrm{bs}$ and $y = \log_{10} F[\mathrm{bs}]$.

In DFA, the fitting error between the two approximating lines (with boundary point $x_A$) and the data points was calculated as

$$e[x_A] = \sum_{i=i_{\mathrm{start}}}^{i_A} (y[i] - a_1 x[i] - b_1) + \sum_{i=i_A}^{i_{\mathrm{end}}} (y[i] - a_2 x[i] - b_2)$$

In contrast, in SDA, the density of data points increases as $x[i]$ increases. Consequently, the above error definition would overfit the large-$x$ region. Therefore, for SDA the error was defined as

$$e[x_A] = \sum_{i=i_{\mathrm{start}}}^{i_A} \frac{y[i] - a_1 x[i] - b_1}{x[i]} + \sum_{i=i_A}^{i_{\mathrm{end}}} \frac{y[i] - a_2 x[i] - b_2}{x[i]}$$

In this study, the optimal boundary point $x_A$ minimizing the error $e$ was searched within the range 0.25 to 10 s. Finally, the slope of the approximating line (scaling exponent) in the short-term region was defined as $a_1$ at the optimal $x_A$, the scaling exponent in the long-term region as $a_2$ at the optimal $x_A$, and the intersection point of two lines was calculated as $(x_A, a_1 x_A + b_1)$ using the optimal $x_A$.

### C Correspondence between $\mu$ and $z$

This section summarizes the characteristics of the corresponding $\boldsymbol{\mu}$-cluster and $\boldsymbol{\zeta}$-cluster pairs. In addition, for clusters that did not exhibit correspondence with any counterpart, we discuss their properties based on the destinations of the surrogate data.

First, we present below the four pairs in which a one-to-one correspondence was observed.

$\boldsymbol{\mu}_1 \Leftrightarrow \boldsymbol{\zeta}_3$

**Destination of surrogate data** For $\boldsymbol{\mu}_1$, 48.7 % of the surrogate data were mapped to the vicinity of $\boldsymbol{\zeta}_3$. Conversely, for $\boldsymbol{\zeta}_3$, 67.3 % of the surrogate data were mapped to the vicinity of $\boldsymbol{\mu}_1$. The distributions in $\boldsymbol{\mu}$- and $\boldsymbol{z}$-spaces are very simmilar to each other (Fig. S1), i.e., $\boldsymbol{\mu}_1$–$\boldsymbol{\zeta}_3$ pair is the corresponding pair.

**Postural control strategy characteristics** Intermittent control was performed with a large proportional gain $p$ and a small derivative gain $D$. With respect to the region on the $p$-$D$ plane in which an inverted pendulum model assuming continuous control can maintain posture (the stability region), the control gain values of this pair were distributed so as to straddle this region. The intervention ratio was approximately $\rho \sim 0.4$, indicating a relatively low intervention frequency even among the intermittent controll groups. The noise intensity $\sigma$ and the dead-zone radius $r$ were also relatively small. The feedback time delay $\Delta$ took a moderate value.

**Postural sway characteristics** Because $z_1$ was small, the proportion of low-frequency components relative to the total power was large. Because $z_5$ was large, the scaling exponent in the high-frequency band was large. Given that $z_3$ was small, indicating that the critical point of DFA was shifted to the left, the scaling exponent in the low-frequency band is considered to be large. In contrast, because $z_4$ was small, the critical point of SDA tended to shift to the right. The travel distance ($z_2$) and the parameter representing the tendency of sign changes in velocity ($z_6$) took values close to zero.

**UPDRS characteristics** This pair exhibited low clinical scores, especially in Total and PIGD scores. $\boldsymbol{\zeta}_3$ shows less number of significant pairs compared to $\boldsymbol{\mu}_1$, and, especially in the $\boldsymbol{z} - space$,
$bm\mu_1$ has wider distribution ranges. Given these, some data point in $\boldsymbol{\mu}_1$ may be classified into other $\boldsymbol{\zeta}$-cluster (e.g., $\boldsymbol{\zeta}_6$), leading increases in clinical scores among the $\boldsymbol{\zeta}$-cluster.

**Remarks** $\boldsymbol{\mu}_1$ was the second largest cluster among the $\boldsymbol{\mu}$-clusters (Table S2). In addition, $\boldsymbol{\mu}_1$ included the largest number of healthy older adults (Table S1). This suggests that this pair is the healthy/mild-sympton cluster.

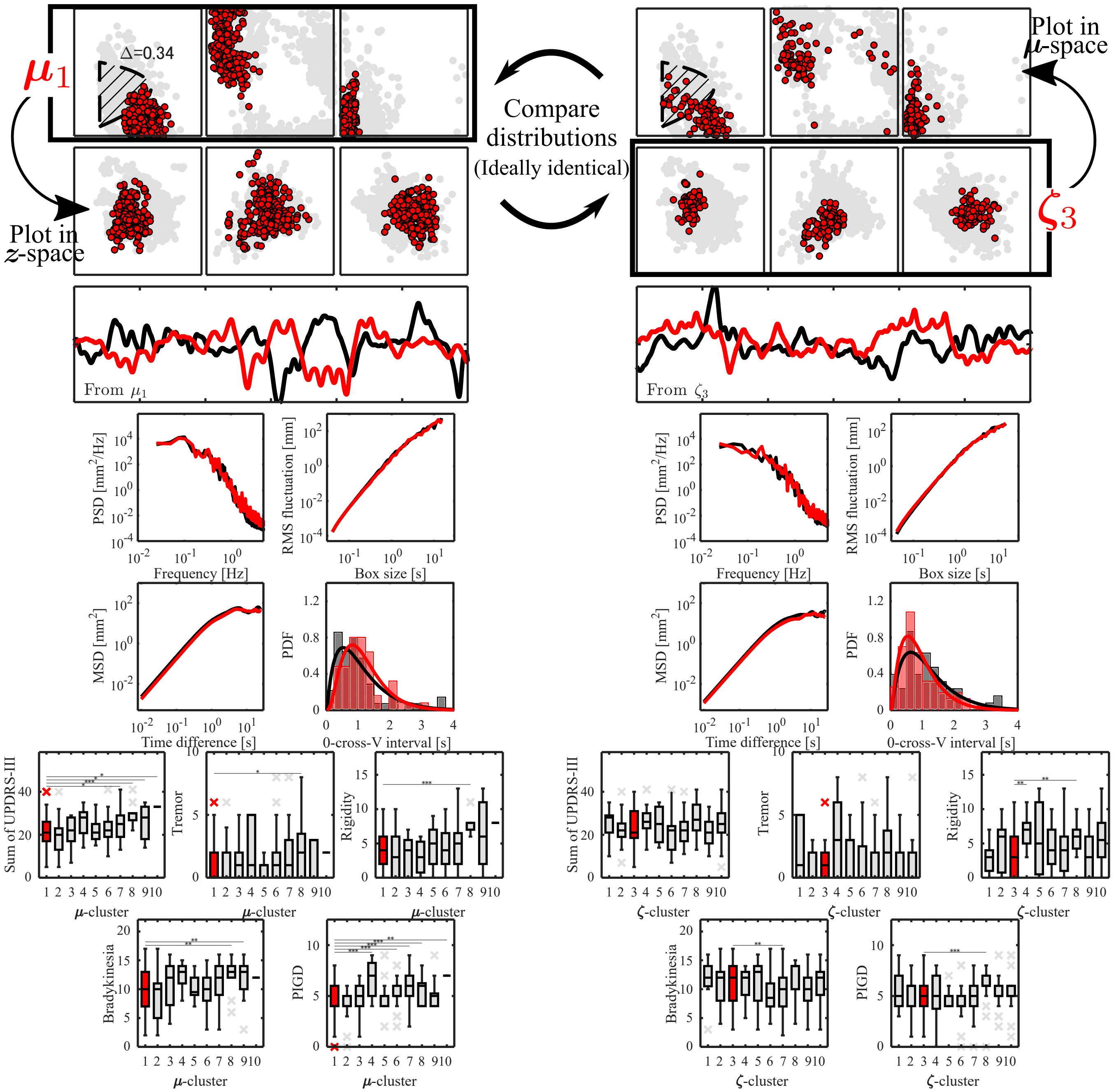


**Fig. S1.** The characteristics of $\mu_1$–$\zeta_3$ pair. The top-left and top-right panels show the distributions in $\mu$-space for the $\mu$-cluster and $\zeta$-cluster, respectively. The axes of $\mu$-space are identical to those in Fig. 3(A). The stable region shown in the $p$-$D$ plane (hatched region) was calculated using the mean of the distribution of $\Delta$ values for each $\mu$-cluster. The left and right panels in the second row show these distributions in $z$-space. The axes of $z$-space are identical to those in Fig. 5(A). Panels from the third row to the fifth row show examples of CoM time-series waveforms (third row), power spectra (fourth row left), DFA plots (fourth row right), SDA plots (fifth row left), and velocity sign-change interval histograms (fifth row right) of representative experimental participants extracted from the corresponding clusters (black: empirical data, red: synthetic data). The bottom-left and bottom-right panels show the boxplots of five clinical scores (Total, Tremor, Rigidity, Bradykinesia, and PIGD scores) among the $\mu$- and $\zeta$-clusters, respectively. *: $p_{bf} < 0.05$, **: $p_{bf} < 0.01$, ***: $p_{bf} < 0.001$, where $p_{bf}$ denotes the $p$-value of the Mann-Whitney U test with Bonferroni correction.

$\boxed{\boldsymbol{\mu}_5 \Leftrightarrow \boldsymbol{\zeta}_{10}}$

**Destination of surrogate data** For $\boldsymbol{\mu}_5$, 96.3 % of the surrogate data were mapped to the vicinity of $\boldsymbol{\zeta}_5$. Conversely, for $\boldsymbol{\zeta}_5$, 50.0 % of the surrogate data were mapped to the vicinity of $\boldsymbol{\mu}_5$. The distributions in $\boldsymbol{\mu}$- and $\boldsymbol{z}$-spaces are very simmilar to each other (Fig. S2).

**Postural control strategy characteristics** $p$ and $\sigma$ are large, while $D$, $\rho$, and $\Delta$ are small. Since $\Delta$ is very small, most data point in $\boldsymbol{\mu}_5$ is in the stable region (hatched region). Thus, this pair may stabilize the posture by switching between stable ON- and unstable OFF-subsystems based on slightly past information. Regarding the physically implausible situation where $\Delta$ was estimated close to 0, it is conceivable that the data in this pair compensate for time delays through prediction, and that this resulted in parameterization as $\Delta \sim 0$ via data assimilation.

**Postural sway characteristics** The postural sway waveforms were characterized by large values in $z_3$ and small values in $z_4$, $z_5$, and $z_6$. In addition, $z_1$ and $z_2$ were distributed around zero. In other words, this group exhibited postural sway waveforms characterized by large high-frequency components and small scaling exponents in the high- and low-frequency bands. Qualitatively, even though the system uses intermittent control switching active control ON and OFF, low-frequency components do not dominate, and the scaling exponents are small. This finding indicates that the existence of a feedback time delay could be a potential cause for the generation of $1/f$-like fluctuation waveforms in intermittent control.

**UPDRS characteristics** The motor severity of PD patients exhibited moderate clinical scores, which significantly differed from those of other clusters that were substantially higher or lower.

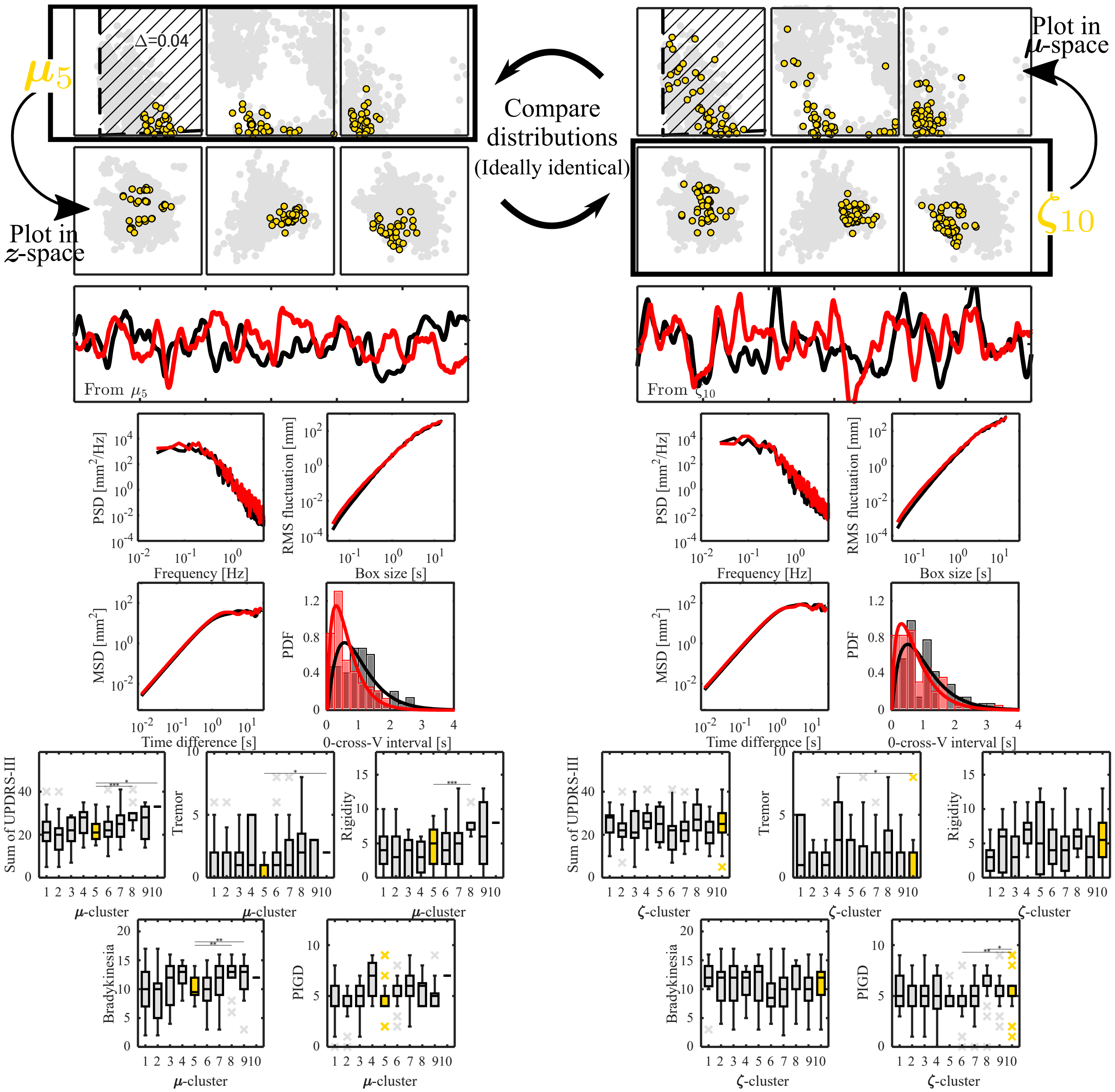


**Fig. S2.** The characteristics of $\mu_5$–$\zeta_{10}$ pair. The top-left and top-right panels show the distributions in $\boldsymbol{\mu}$-space for the $\boldsymbol{\mu}$-cluster and $\zeta$-cluster, respectively. The axes of $\boldsymbol{\mu}$-space are identical to those in Fig. 3(A). The stable region shown in the $p$-$D$ plane (hatched region) was calculated using the mean of the distribution of $\Delta$ values for each $\boldsymbol{\mu}$-cluster. The left and right panels in the second row show these distributions in $\boldsymbol{z}$-space. The axes of $\boldsymbol{z}$-space are identical to those in Fig. 5(A). Panels from the third row to the fifth row show examples of CoM time-series waveforms (third row), power spectra (fourth row left), DFA plots (fourth row right), SDA plots (fifth row left), and velocity sign-change interval histograms (fifth row right) of representative experimental participants extracted from the corresponding clusters (black: empirical data, red: synthetic data). The bottom-left and bottom-right panels show the boxplots of five clinical scores (Total, Tremor, Rigidity, Bradykinesia, and PIGD scores) among the $\boldsymbol{\mu}$- and $\boldsymbol{\zeta}$-clusters, respectively. *: $p_{bf} < 0.05$, **: $p_{bf} < 0.01$, ***: $p_{bf} < 0.001$, where $p_{bf}$ denotes the $p$-value of the Mann-Whitney U test with Bonferroni correction.

 K. Matsui, Y. Suzuki, C. E. Smith, T. Nakamura, T. Endo, S. Sakoda, P. G. Morasso, and T. Nomura

$\boxed{\boldsymbol{\mu}_8 \Leftrightarrow \boldsymbol{\zeta}_4}$

**Destination of surrogate data** For $\boldsymbol{\mu}_8$, 34.4 % of the surrogate data were mapped to the vicinity of $\boldsymbol{\zeta}_4$. Conversely, for $\boldsymbol{\zeta}_4$, 36.9 % of the surrogate data were mapped to the vicinity of $\boldsymbol{\mu}_8$. The distributions in $\boldsymbol{\mu}$- and $\boldsymbol{z}$-spaces are very simmilar to each other (Fig. S3). Compared with other $\mu?\zeta$ cluster pairs, this number is slightly lower. This implies either low accuracy of the neural network that learned the mapping function, a high frequency of data belonging to $\boldsymbol{\mu}_8$ ($\boldsymbol{\zeta}_4$) being classified into $\zeta$-clusters ($\mu$-clusters) other than $\boldsymbol{\zeta}_4$ ($\boldsymbol{\mu}_8$), or a combination of both. Looking at Table S2, it can indeed be observed that PD patients in $\boldsymbol{\mu}_8$ are distributed across various $\zeta$-clusters. Similarly, for $\boldsymbol{\zeta}_4$, ten or more data points are assigned to $\boldsymbol{\mu}_1$ or $\boldsymbol{\mu}_7$.

**Postural control strategy characteristics** Rigid continuous control was performed with large $p$ and $D$. $\Delta$ was close to 0, and this is physically implausible as described in above. As a control policy that can be parameterized in this way, passive postural control due to the high biomechanical viscoelasticity of ankle muscles is a representative example. Model parameter values of $\Delta \approx 0$ and $\rho \approx 1$ result in adding the proportional gain and derivative gain directly to the elasticity and viscosity coefficients of the ankle muscles. Hence, it can be interpreted that conscious or unconscious co-contraction of the ankle muscles increases passive viscoelasticity. In fact, PD patients categorized into $\boldsymbol{\mu}_8$ and $\boldsymbol{\zeta}_4$ exhibited significantly higher muscle rigidity scores compared to other clusters.

**Postural sway characteristics** This group exhibits postural sway waveforms characterized by a high scaling exponent and a dominance of low-frequency components in the low-frequency band. Conversely, high-frequency components are prominent, whereas the scaling exponent for this band is slightly smaller. Furthermore, the overall power is lower than that of other postural sway waveforms, yielding a reduced sway area.

**UPDRS characteristics** This pair exhibited low clinical scores, especially in Total and PIGD scores. $\boldsymbol{\zeta}_3$ shows less number of significant pairs compared to $\boldsymbol{\mu}_1$, and, especially in the $\boldsymbol{z} - space$,
$bm\mu_1$ has wider distribution ranges. Given these, some data point in $\boldsymbol{\mu}_1$ may be classified into other $\boldsymbol{\zeta}$-cluster (e.g., $\boldsymbol{\zeta}_6$), leading increases in clinical scores among the $\boldsymbol{\zeta}$-cluster.

**Remarks** This pair showed the highest Total score among $\boldsymbol{\mu}$- and $\boldsymbol{\zeta}$-clusters.

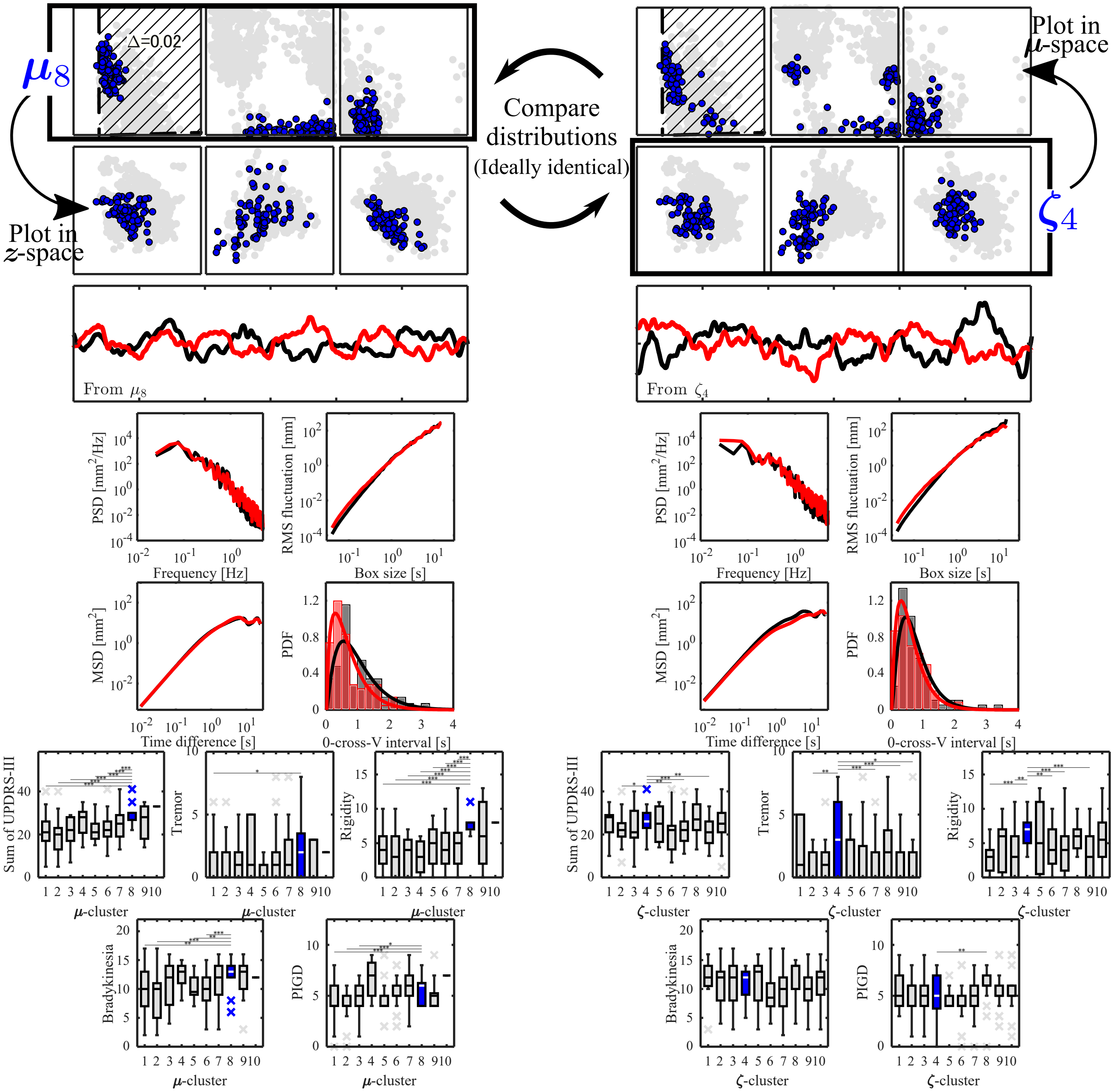


**Fig. S3.** The characteristics of $\mu_8$–$\zeta_4$ pair. The top-left and top-right panels show the distributions in $\mu$-space for the $\mu$-cluster and $\zeta$-cluster, respectively. The axes of $\mu$-space are identical to those in Fig. 3(A). The stable region shown in the $p$-$D$ plane (hatched region) was calculated using the mean of the distribution of $\Delta$ values for each $\mu$-cluster. The left and right panels in the second row show these distributions in $z$-space. The axes of $z$-space are identical to those in Fig. 5(A). Panels from the third row to the fifth row show examples of CoM time-series waveforms (third row), power spectra (fourth row left), DFA plots (fourth row right), SDA plots (fifth row left), and velocity sign-change interval histograms (fifth row right) of representative experimental participants extracted from the corresponding clusters (black: empirical data, red: synthetic data). The bottom-left and bottom-right panels show the boxplots of five clinical scores (Total, Tremor, Rigidity, Bradykinesia, and PIGD scores) among the $\mu$- and $\zeta$-clusters, respectively. *: $p_{bf} < 0.05$, **: $p_{bf} < 0.01$, ***: $p_{bf} < 0.001$, where $p_{bf}$ denotes the $p$-value of the Mann-Whitney U test with Bonferroni correction.

$\boxed{\boldsymbol{\mu}_9 \Leftrightarrow \boldsymbol{\zeta}_5}$

**Destination of surrogate data** For $\boldsymbol{\mu}_9$, 64.5 % of the surrogate data were mapped to the vicinity of $\boldsymbol{\zeta}_5$. Conversely, for $\boldsymbol{\zeta}_5$, 91.8 % of the surrogate data were mapped to the vicinity of $\boldsymbol{\mu}_9$. The distributions in $\boldsymbol{\mu}$- and $\boldsymbol{z}$-spaces are very simmilar to each other (Fig. S4).

**Postural control strategy characteristics** The postural control policy is similar to that of the $\boldsymbol{\mu}_8$?-$\boldsymbol{\zeta}_4$ cluster pair. Specifically, $\Delta$ is small, whereas $D$, $\rho$, and $\sigma$ are large. However, this set exhibits a larger $p$, with $\rho$ concentrated in the range of 0.9-?1 and a larger $\sigma$. Therefore, it is inferred that posture is maintained through high biomechanical viscoelasticity resulting from greater muscle co-contraction under noisier environments. Regarding the Rigidity score of PD patients, although the variance was large, this group also exhibited high values, similar to the $\boldsymbol{\mu}_8$?-$\boldsymbol{\zeta}_4$ cluster pair.

**Postural sway characteristics** $z_3$, $z_4$, and $z_5$ were small, $z_1$ and $z_2$ were large, and $z_6$ was distributed around zero. In other words, the high-frequency components are prominent, whereas the scaling exponent in this frequency band is small. Unlike the $\boldsymbol{\mu}_8$?$\boldsymbol{\zeta}_4$ cluster pair, low-frequency components do not dominate; nevertheless, both the total signal power and the sway area are larger than those for the $\boldsymbol{\mu}_8$?$\boldsymbol{\zeta}_4$ pair. The power spectrum has a peculiar shape that resembles triple scaling raw, possessing different scaling exponents across three frequency bands: below 0.1 Hz, 0.1?1 Hz, and above 1 Hz. The slope for the frequency range up to 0.1 Hz is nearly zero, which is considered to appear as the prominent feature of $z_1$.

**UPDRS characteristics** Regarding the box plots of clinical scores, $\boldsymbol{\mu}_9$ exhibits a high Rigidity score despite having a wide interquartile range (long box). Although $\boldsymbol{\zeta}_5$ showed a similar trend, no significant differences were observed compared to other $\zeta$-clusters. Furthermore, this group exhibited the highest Bradykinesia score among all groups. On the other hand, unlike the $\boldsymbol{\mu}_8$?$\boldsymbol{\zeta}_4$ cluster pair, the Total score, Tremor score, and PIGD score showed small-to-moderate values.

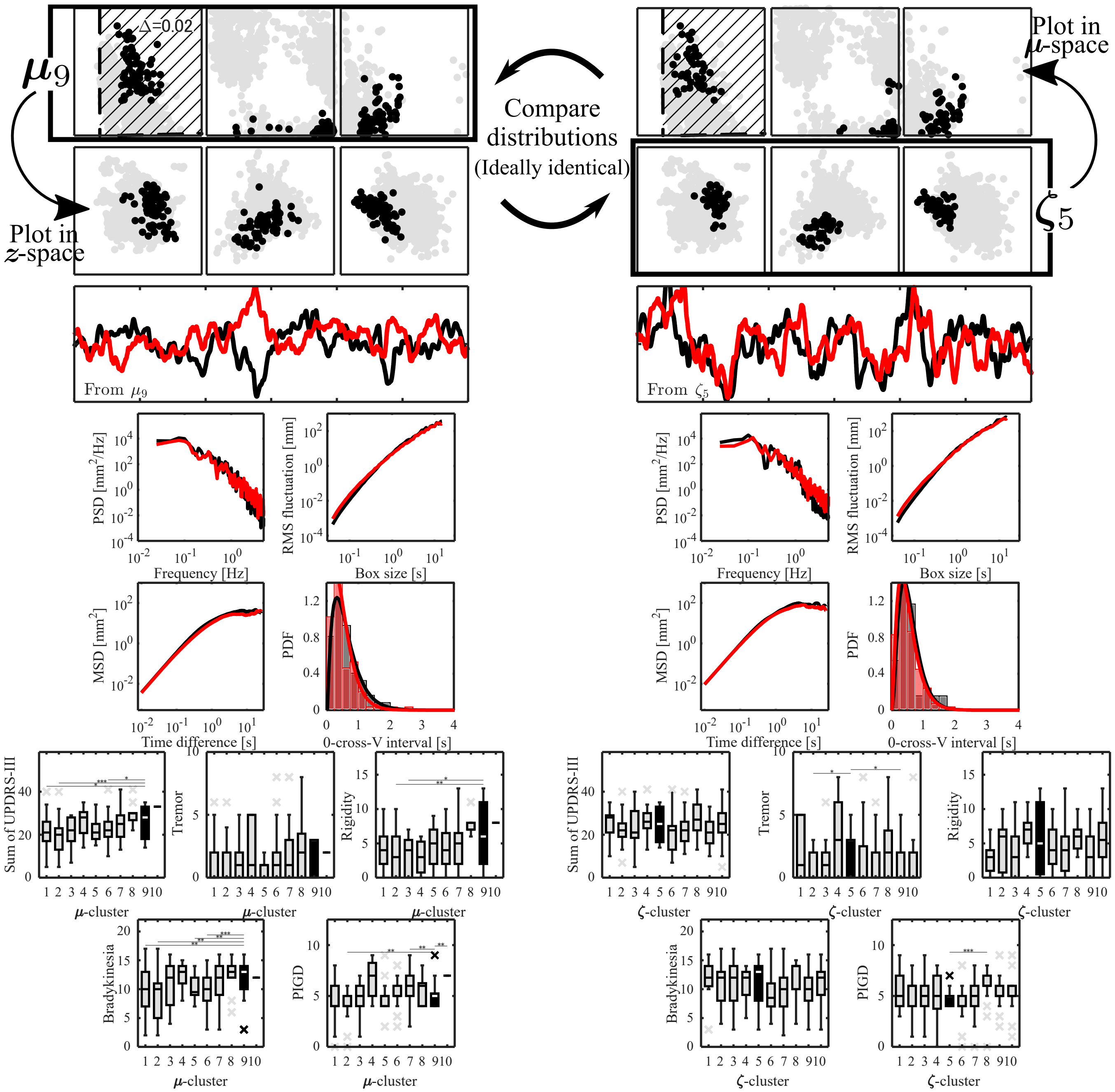


**Fig. S4.** The characteristics of $\mu_9$–$\zeta_5$ pair. The top-left and top-right panels show the distributions in $\mu$-space for the $\mu$-cluster and $\zeta$-cluster, respectively. The axes of $\mu$-space are identical to those in Fig. 3(A). The stable region shown in the $p$-$D$ plane (hatched region) was calculated using the mean of the distribution of $\Delta$ values for each $\mu$-cluster. The left and right panels in the second row show these distributions in $z$-space. The axes of $z$-space are identical to those in Fig. 5(A). Panels from the third row to the fifth row show examples of CoM time-series waveforms (third row), power spectra (fourth row left), DFA plots (fourth row right), SDA plots (fifth row left), and velocity sign-change interval histograms (fifth row right) of representative experimental participants extracted from the corresponding clusters (black: empirical data, red: synthetic data). The bottom-left and bottom-right panels show the boxplots of five clinical scores (Total, Tremor, Rigidity, Bradykinesia, and PIGD scores) among the $\mu$- and $\zeta$-clusters, respectively. *: $p_{bf} < 0.05$, **: $p_{bf} < 0.01$, ***: $p_{bf} < 0.001$, where $p_{bf}$ denotes the $p$-value of the Mann-Whitney U test with Bonferroni correction.

Next, we present the three pairs in which a many-to-one correspondence was observed.

$(\boldsymbol{\mu}_2, \boldsymbol{\mu}_6) \Leftrightarrow \boldsymbol{\zeta}_7$

**Destination of surrogate data** 60.4 % of the surrogate data generated from $\boldsymbol{\mu}_2$ and 55.8 % of those generated from $\boldsymbol{\mu}_6$ were mapped to the vicinity of $\boldsymbol{\zeta}_7$. Conversely, 34.5 % of the surrogate data generated from $\boldsymbol{\zeta}_7$ were mapped to the vicinity of $\boldsymbol{\mu}_2$, and 41.1 % were mapped to the vicinity of $\boldsymbol{\mu}_6$. The distributions in $\boldsymbol{\mu}$- and $\boldsymbol{z}$-spaces are very simmilar to each other (Fig. S5).

**Postural control strategy characteristics** $\boldsymbol{\mu}_2$ utilizes intermittent active feedback control, similar to $\boldsymbol{\mu}_1$ mentioned earlier. The key difference lies in the magnitude of the control gains; $\boldsymbol{\mu}_2$ features a larger derivative gain. As a result, the data points belonging to $\boldsymbol{\mu}_2$ are distributed inside the stable region of the inverted pendulum, and the vector field when active control is operative forms a stable focus that spirals and converges toward the equilibrium point. Therefore, $\boldsymbol{\mu}_2$ maintains an upright posture by switching between an unstable OFF-subsystem and a stable ON-subsystem in a posture-state-dependent manner.

In contrast, $\boldsymbol{\mu}_6$ is considered to stabilize posture through continuous active feedback control. In $\boldsymbol{\mu}_6$, the value of $\rho$ is close to 1, while the other model parameters show distributions similar to those of $\boldsymbol{\mu}_2$. The value of $\rho$ reflects the frequency of active control intervention, where a larger value corresponds to a longer duration in which active control is active (ON). When $\rho = 1$, active control is applied continuously. Thus, the postural control policy of $\boldsymbol{\mu}_6$ is highly likely to sustain upright posture via the stable focus formed by large control gains within the state space.

**Postural sway characteristics** Compared with the postural sway of the $\boldsymbol{\mu}_1$?$\boldsymbol{\zeta}_3$ cluster pair, the low-frequency components and sway area are smaller. As a result, the low-frequency band in the ($\boldsymbol{\mu}_2$, $\boldsymbol{\mu}_6$)-?$\boldsymbol{\zeta}_7$ cluster pairs was whitened. In addition, although the increase is small, the high-frequency components are larger than those of the $\boldsymbol{\mu}_1$?-$\boldsymbol{\zeta}_3$ cluster pair. A potential explanation for the reduction in sway area and low-frequency components is that the increased control gains?particularly the derivative gain—shift the overall direction of the vector field during the ON-subsystem toward the equilibrium point, thereby preventing large sway.

**UPDRS characteristics** $\boldsymbol{\mu}_2$, inferred to employ intermittent control, exhibited the lowest values among all $\mu$-clusters across the Total, Rigidity, and PIGD scores. In contrast, $\boldsymbol{\mu}_6$, inferred to utilize continuous control, showed moderate-to-high values for both the Total and PIGD scores. Additionally, $\boldsymbol{\zeta}_7$ displayed low-to-moderate clinical scores.

**Remarks** Despite the differences in postural control policy and motor severity of the included PD patients between $\boldsymbol{\mu}_2$ and $\boldsymbol{\mu}_6$, the observed postural sway waveforms were shown to be similar. Furthermore, PD patients belonging to $\boldsymbol{\zeta}_7$, a cluster based on postural sway metrics, were statistically classified as having low-to-moderate severity. These findings suggest that severity diagnosis using simple postural sway metrics (such as sway area) is highly likely to overlook severely affected PD patients, such as those belonging to $\boldsymbol{\mu}_6$. On the other hand, utilizing the standing posture digital twin constructed in this study enables the use of $\mu$-values, which demonstrated high discriminative ability in severity classification, thereby expected to provide more advanced and accurate support for severity diagnosis.

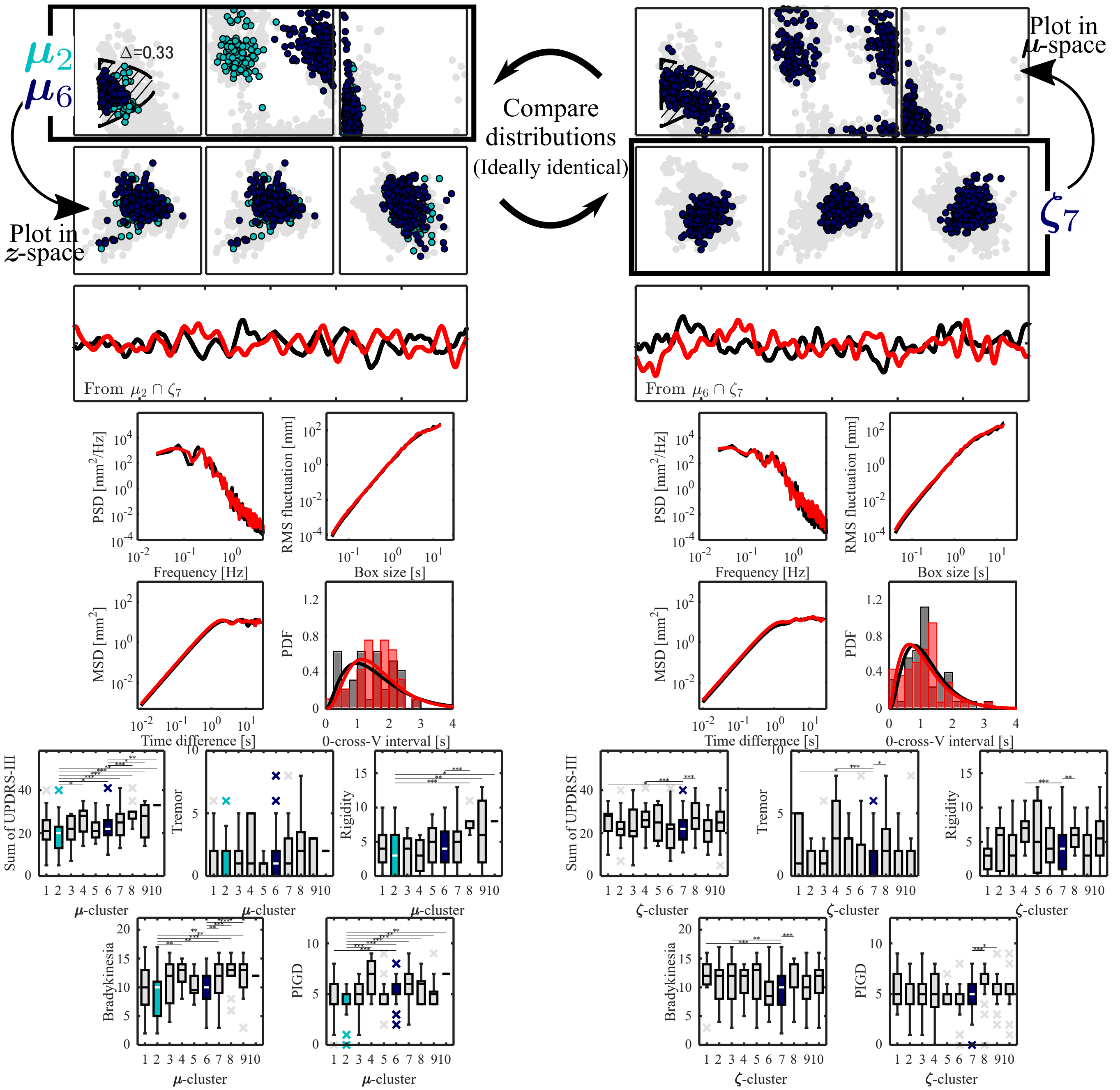


**Fig. S5.** The characteristics of $(\mu_2, \mu_6)$–$\zeta_7$ pair. The top-left and top-right panels show the distributions in $\mu$-space for the $\mu$-cluster and $\zeta$-cluster, respectively. The axes of $\mu$-space are identical to those in Fig. 3(A). The stable region shown in the $p$-$D$ plane (hatched region) was calculated using the mean of the distribution of $\Delta$ values for each $\mu$-cluster. The left and right panels in the second row show these distributions in $z$-space. The axes of $z$-space are identical to those in Fig. 5(A). Panels from the third row to the fifth row show examples of CoM time-series waveforms (third row), power spectra (fourth row left), DFA plots (fourth row right), SDA plots (fifth row left), and velocity sign-change interval histograms (fifth row right) of representative experimental participants extracted from the corresponding clusters (black: empirical data, red: synthetic data). The bottom-left and bottom-right panels show the boxplots of five clinical scores (Total, Tremor, Rigidity, Bradykinesia, and PIGD scores) among the $\mu$- and $\zeta$-clusters, respectively. *: $p_{bf} < 0.05$, **: $p_{bf} < 0.01$, ***: $p_{bf} < 0.001$, where $p_{bf}$ denotes the $p$-value of the Mann-Whitney U test with Bonferroni correction.

 

$(\boldsymbol{\mu}_3, \boldsymbol{\mu}_4) \Leftrightarrow \boldsymbol{\zeta}_1$

**Destination of surrogate data** 60.4 % of the surrogate data generated from $\boldsymbol{\mu}_3$ and 55.8 % of those generated from $\boldsymbol{\mu}_4$ were mapped to the vicinity of $\boldsymbol{\zeta}_1$. Conversely, 34.5 % of the surrogate data generated from $\boldsymbol{\zeta}_1$ were mapped to the vicinity of $\boldsymbol{\mu}_3$, and 41.1 % were mapped to the vicinity of $\boldsymbol{\mu}_4$. The distributions in $\boldsymbol{\mu}$- and $\boldsymbol{z}$-spaces are very simmilar to each other (Fig. S6).

**Postural control strategy characteristics** Both $\boldsymbol{\mu}_3$ and $\boldsymbol{\mu}_4$ are inferred to employ a postural control policy similar to that of $\boldsymbol{\mu}_1$; that is, they adopt postural stabilization via intermittent active feedback control using small control gains distributed outside the stable region. As for their differences, $\boldsymbol{\mu}_3$ is characterized by a large feedback time delay, whereas $\boldsymbol{\mu}_4$ exhibits high noise intensity.

**Postural sway characteristics** $z_1$ is low and $z_5$ is high, indicating prominent dominance of low-frequency components, low magnitude in high-frequency components, and high scaling exponents across both low- and high-frequency bands. Nevertheless, due to a large $z_3$ and small $z_4$, the critical points of the DFA and SDA plots are right-shifted, resulting in slightly reduced scaling exponents for each frequency band. Furthermore, the small value of $z_6$, together with the elevated $z_5$, yields a gamma distribution—fitted to the histogram of time intervals between sign changes in the velocity time series—characterized by a peak closer to zero and a heavy (long) tail. In addition, a characteristic feature is that $z_2$ is large. This is reflected in the large sway area; in fact, the data point with the highest RMS among all data belonged to $\boldsymbol{\mu}_4$ and $\boldsymbol{\zeta}_1$.

**UPDRS characteristics** This pair showed moderate-to-high Bradykinesia and PIGD scores.

**Remarks** Although the real dataset included only four PD patients belonging to $\boldsymbol{\mu}_4$, according to our clinical collaborators, all four presented symptoms resembling atypical parkinsonism and were subsequently diagnosed with multiple system atrophy (MSA) during follow-up evaluations after the measurement date. This suggests that $\boldsymbol{\mu}_4$ represents an MSA patient group. In addition, $\boldsymbol{\zeta}_1$ corresponds to both $\boldsymbol{\mu}_3$ and $\boldsymbol{\mu}_4$?meaning that $\boldsymbol{\mu}_3$ and $\boldsymbol{\mu}_4$ exhibit remarkably similar postural sway waveforms?and $\boldsymbol{\mu}_3$ showed a lower PIGD score than $\boldsymbol{\mu}_4$. These findings suggest the possibility that $\boldsymbol{\mu}_3$ includes patients in the early stages of MSA, and that as the disease progresses and worsens, noise in the neural control system for posture increases, causing patients in $\boldsymbol{\mu}_3$ to transition into the $\boldsymbol{\mu}_4$ group.

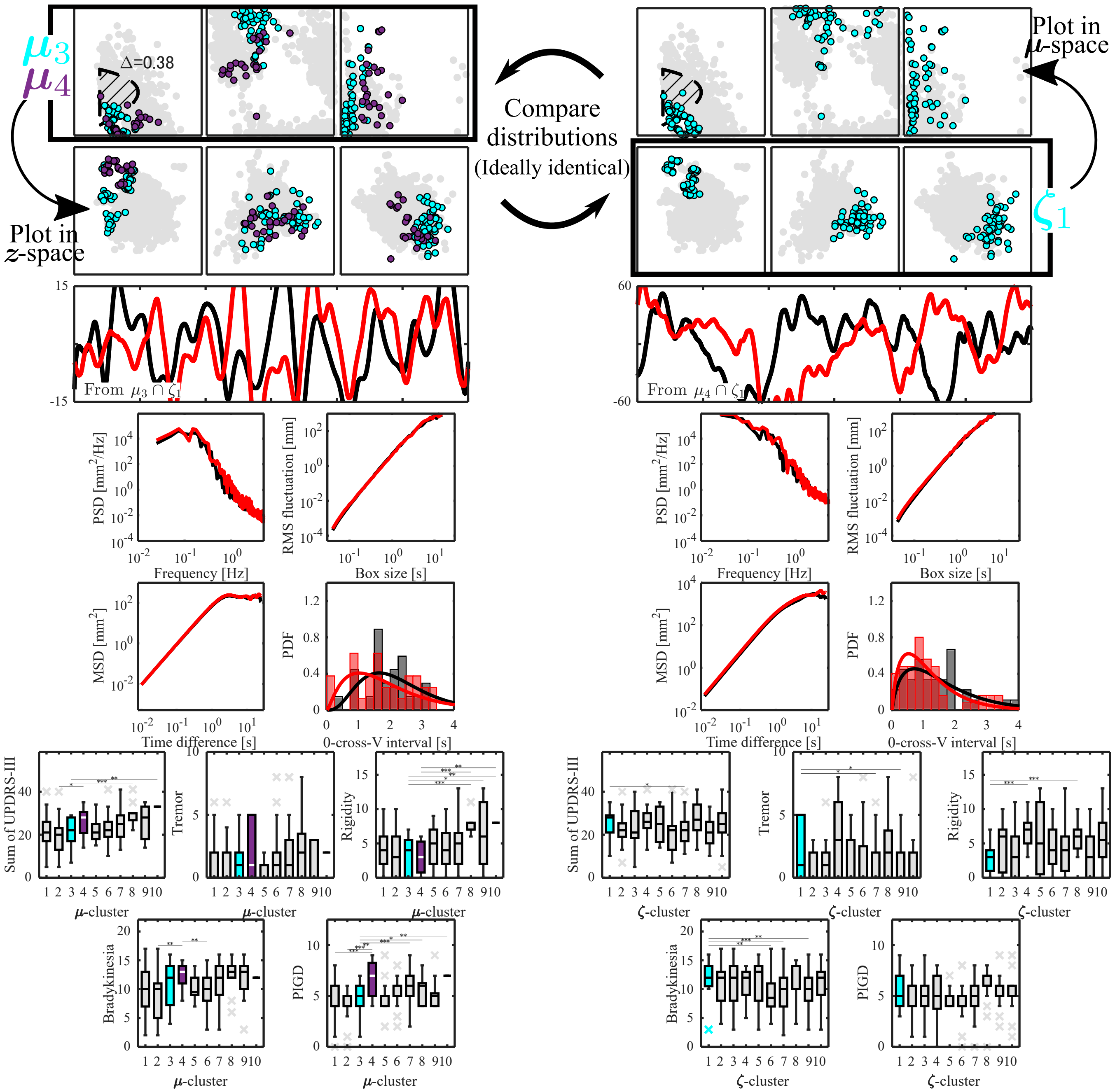


**Fig. S6.** The characteristics of $(\boldsymbol{\mu}_3, \boldsymbol{\mu}_4)$–$\boldsymbol{\zeta}_1$ pair. The top-left and top-right panels show the distributions in $\boldsymbol{\mu}$-space for the $\boldsymbol{\mu}$-cluster and $\boldsymbol{\zeta}$-cluster, respectively. The axes of $\boldsymbol{\mu}$-space are identical to those in Fig. 3(A). The stable region shown in the $p$-$D$ plane (hatched region) was calculated using the mean of the distribution of $\Delta$ values for each $\boldsymbol{\mu}$-cluster. The left and right panels in the second row show these distributions in $\boldsymbol{z}$-space. The axes of $\boldsymbol{z}$-space are identical to those in Fig. 5(A). Panels from the third row to the fifth row show examples of CoM time-series waveforms (third row), power spectra (fourth row left), DFA plots (fourth row right), SDA plots (fifth row left), and velocity sign-change interval histograms (fifth row right) of representative experimental participants extracted from the corresponding clusters (black: empirical data, red: synthetic data). The bottom-left and bottom-right panels show the boxplots of five clinical scores (Total, Tremor, Rigidity, Bradykinesia, and PIGD scores) among the $\boldsymbol{\mu}$- and $\boldsymbol{\zeta}$-clusters, respectively. *: $p_{bf} < 0.05$, **: $p_{bf} < 0.01$, ***: $p_{bf} < 0.001$, where $p_{bf}$ denotes the $p$-value of the Mann-Whitney U test with Bonferroni correction.

$\boxed{\boldsymbol{\mu}_7 \Leftrightarrow (\boldsymbol{\zeta}_8, \boldsymbol{\zeta}_9)}$

**Destination of surrogate data** 40.5 % of the surrogate data generated from $\boldsymbol{\mu}_7$ were mapped to the vicinity of $\boldsymbol{\zeta}_8$, and 46.6 % were mapped to the vicinity of $\boldsymbol{\zeta}_9$. Conversely, 69.6 % of the surrogate data generated from $\boldsymbol{\zeta}_8$ and 65.1 % of those generated from $\boldsymbol{\zeta}_9$ were mapped to the vicinity of $\boldsymbol{\mu}_7$. The distributions in $\boldsymbol{\mu}$- and $\boldsymbol{z}$-spaces are very simmilar to each other (Fig. S7).

**Postural control strategy characteristics** Regarding the postural control policy, $\boldsymbol{\mu}_7$ is inferred to employ continuous active feedback control, similar to $\boldsymbol{\mu}_6$. Three main differences are observed: a slightly larger derivative gain, a narrower distribution range of feedback time delay, and higher noise intensity. Therefore, compared with $\boldsymbol{\mu}_6$, it is considered that $\boldsymbol{\mu}_7$ stabilizes upright posture via continuous control with larger gains under a higher-noise environment.

**Postural sway characteristics** As for the postural sway characteristics, $\boldsymbol{\zeta}_8$ and $\boldsymbol{\zeta}_9$ exhibited highly similar distributions, with the exception of $z_2$, which reflects the total path length. In particular, they were characterized by elevated $z_1$, $z_3$, and $z_6$, reduced $z_5$, and $z_3$ values clustered around zero. Consequently, this set features postural sway waveforms characterized by a prominent dominance of high-frequency components alongside small scaling exponents across both low- and high-frequency regimes. In addition, the distribution of time intervals between velocity sign changes displays a left-skewed histogram (biased toward 0 s) accompanied by a heavy tail.

A distinct difference was noted for $z_2$, where $\boldsymbol{\zeta}_8$ showed low values and $\boldsymbol{\zeta}_9$ showed high values. This primarily emerges in the observed waveforms as the size of the sway area, with $\boldsymbol{\zeta}_9$ exhibiting greater wave amplitude at frequencies slightly lower than 1 Hz.

**UPDRS characteristics** Regarding the motor severity of PD patients, this set is characterized by a high severity in postural function (PIGD score). $\boldsymbol{\mu}_7$ showed the second highest PIGD score among all $\mu$-clusters, while $\boldsymbol{\zeta}_8$ exhibited the highest PIGD score among all $\zeta$-clusters. $\boldsymbol{\zeta}_9$ displayed a moderate PIGD score value. Regarding other clinical scores, $\boldsymbol{\zeta}_9$ showed relatively low values, whereas $\boldsymbol{\zeta}_8$ exhibited relatively high values.

**Remarks** The results for this group and the aforementioned $\boldsymbol{\mu}_6$ suggest that the loss of intermittency in active feedback control for postural regulation may be linked to the deterioration of postural function in PD patients. In addition, between the two $\zeta$-clusters ($\boldsymbol{\zeta}_8$ and $\boldsymbol{\zeta}_9$) corresponding to $\boldsymbol{\mu}_7$, $\boldsymbol{\zeta}_8$—which exhibits a smaller sway area—showed significantly higher values across clinical scores, including the PIGD score, compared with $\boldsymbol{\zeta}_9$. This suggests that even among PD patients who have lost active control intermittency, those with more severe motor symptoms may adopt a control strategy (a compensatory strategy) that increases the degree of muscle contraction to suppress postural sway.

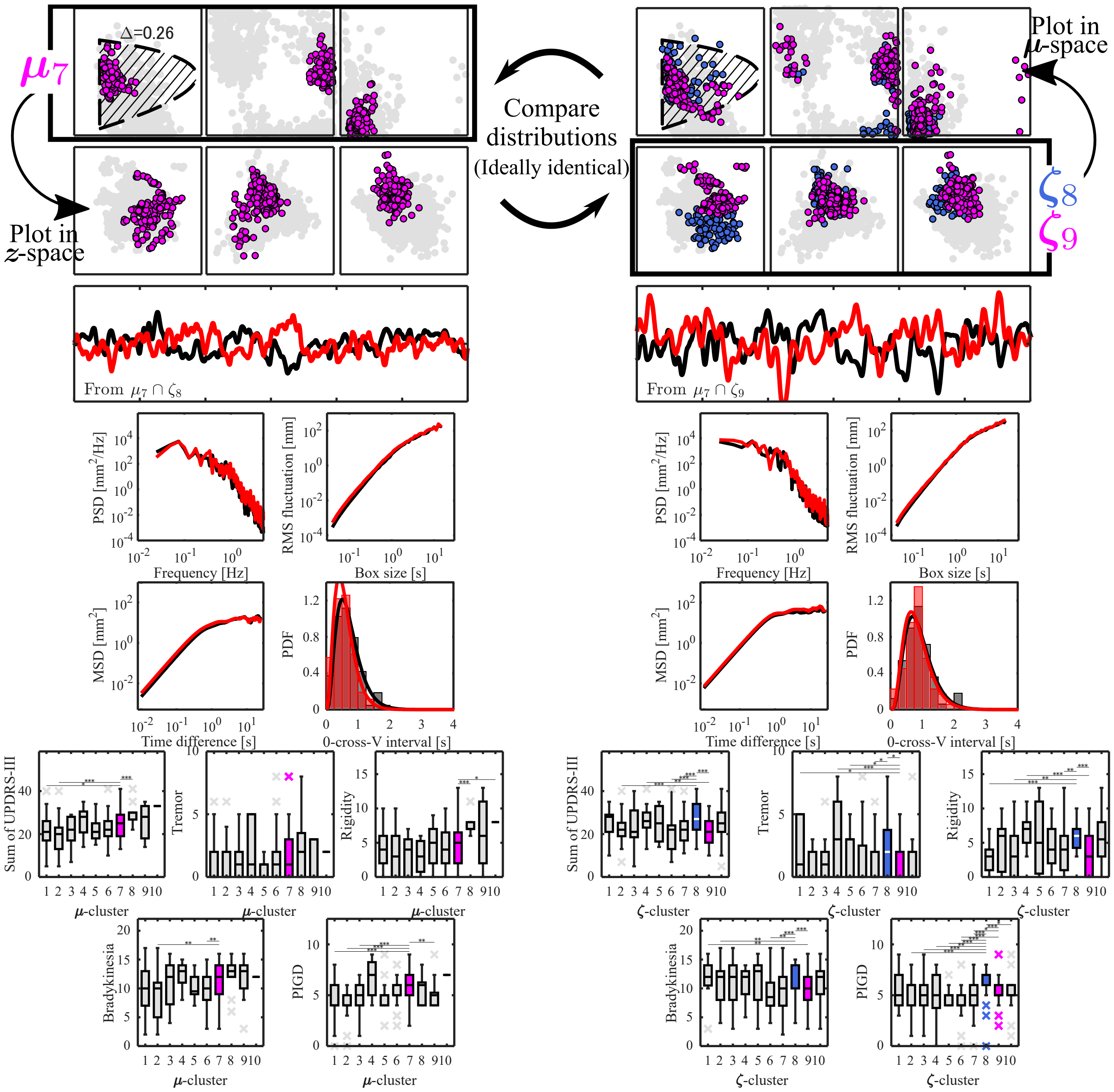


**Fig. S7.** The characteristics of $\mu_7$–$(\zeta_8, \zeta_9)$ pair. The top-left and top-right panels show the distributions in $\mu$-space for the $\mu$-cluster and $\zeta$-cluster, respectively. The axes of $\mu$-space are identical to those in Fig. 3(A). The stable region shown in the $p$-$D$ plane (hatched region) was calculated using the mean of the distribution of $\Delta$ values for each $\mu$-cluster. The left and right panels in the second row show these distributions in $z$-space. The axes of $z$-space are identical to those in Fig. 5(A). Panels from the third row to the fifth row show examples of CoM time-series waveforms (third row), power spectra (fourth row left), DFA plots (fourth row right), SDA plots (fifth row left), and velocity sign-change interval histograms (fifth row right) of representative experimental participants extracted from the corresponding clusters (black: empirical data, red: synthetic data). The bottom-left and bottom-right panels show the boxplots of five clinical scores (Total, Tremor, Rigidity, Bradykinesia, and PIGD scores) among the $\mu$- and $\zeta$-clusters, respectively. *: $p_{bf} < 0.05$, **: $p_{bf} < 0.01$, ***: $p_{bf} < 0.001$, where $p_{bf}$ denotes the $p$-value of the Mann-Whitney U test with Bonferroni correction.

In the following, we describe clusters for which no consistent correspondence was observed and provide considerations based on the destinations of the surrogate data.

$\boldsymbol{\mu}_{10}(\Leftrightarrow \boldsymbol{\zeta}_9)$

**Destination of surrogate data** Of the surrogate data of $\boldsymbol{\mu}_{10}$, 100.0 % were mapped near $\boldsymbol{\zeta}_9$. In contrast, only 0.5 % of the surrogate data of $\boldsymbol{\zeta}_9$ were mapped near $\boldsymbol{\mu}_{10}$. This asymmetry is likely attributable to the fact that $\boldsymbol{\mu}_{10}$ is a unique cluster consisting of data from only a single participant, and the data in this cluster took an outlier-like value within $\boldsymbol{\zeta}_9$. In the $z_1$-$z_2$ plane, $\boldsymbol{\mu}_{10}$ was located in the upper-right region, separated from the main population of $\boldsymbol{\zeta}_9$ (Fig. S8).

**Characteristics of postural control strategies** Regarding the postural control policy, $\boldsymbol{\mu}_{10}$ featured large values of $p$ and $D$, a $\rho$ close to 1, and a $\Delta$ of approximately 0.1. A distinct feature compared to other $\mu$-clusters was its exceptionally large $\sigma$. Based on these model parameter values, $\boldsymbol{\mu}_{10}$ is inferred to stabilize posture using continuous active feedback control in an environment with even higher noise than $\boldsymbol{\mu}_7$. The reason why $\boldsymbol{\mu}_{10}$ was embedded closer to $\boldsymbol{M}_{\mathrm{STIFF}}$ rather than $\boldsymbol{M}_{\mathrm{CONT}}$ in t-SNE can be attributed to its high noise intensity, which likely caused it to be pulled toward $\boldsymbol{\mu}_9$, the group with the largest $\sigma$ excluding $\boldsymbol{\mu}_{10}$.

**Characteristics of postural sway** Similar to $\boldsymbol{\zeta}_9$, this cluster possesses postural sway waveforms in which high-frequency components are strongly dominant, with small scaling exponents in both the low- and high-frequency bands. On top of that, the power across the entire frequency range is large, resulting in a large sway area.

**UPDRS characteristics** Although the statistical robustness is low due to this cluster comprising only one PD patient, $\boldsymbol{\mu}_{10}$ showed significantly elevated values in four clinical scores, with the exception of the Tremor score.

**Remarks** Postural sway waveforms such as those observed in this cluster, which are rare and specific, may be excluded as outliers depending on the analysis method. The present digital twin framework enables augmentation of such rare data and facilitates their inclusion in statistical analyses.

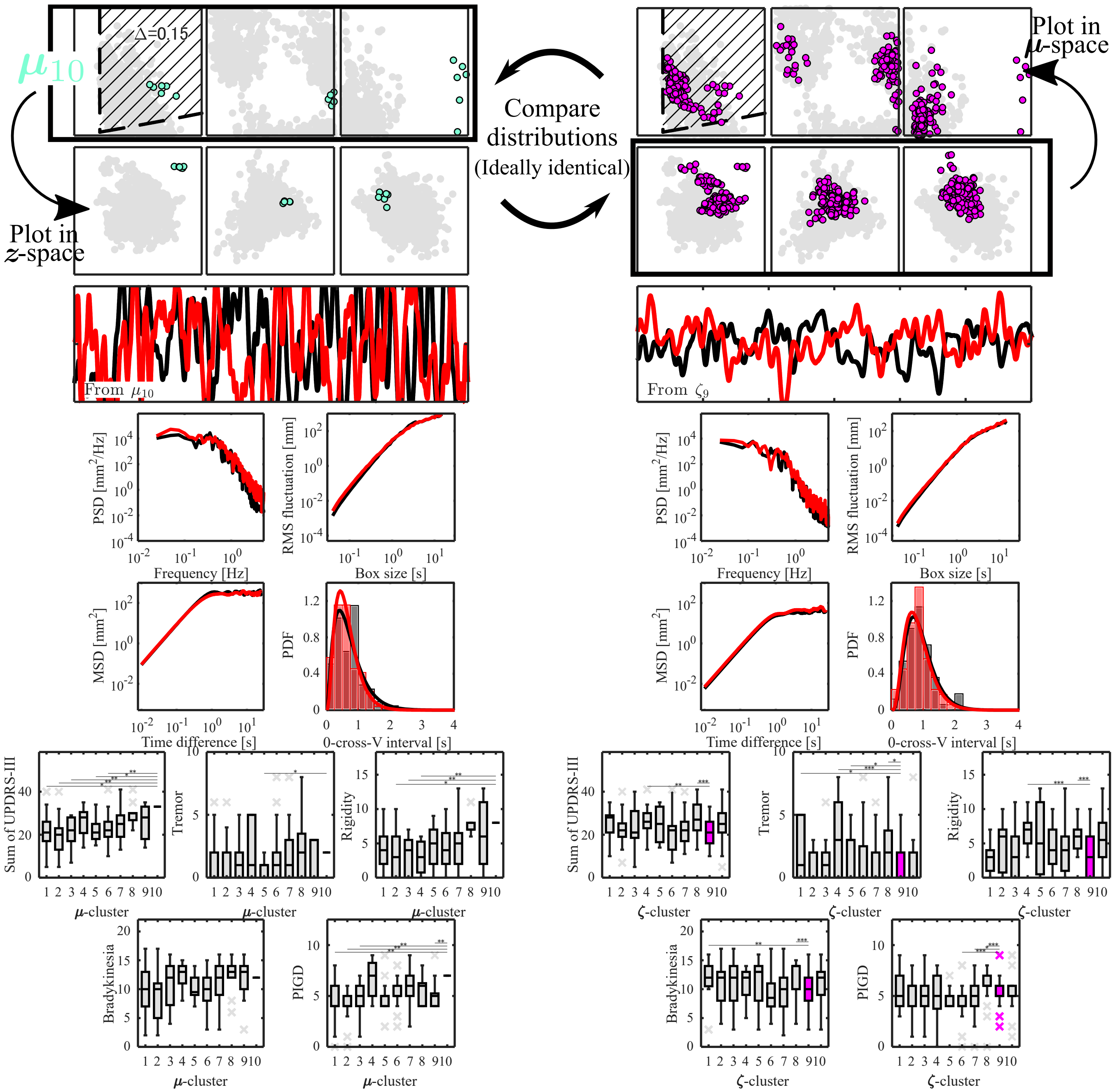

**Fig. S8.** The characteristics of $\mu_{10}$. The top-left and top-right panels show the distributions in $\mu$-space for the $\mu$-cluster and $\zeta$-cluster, respectively. The axes of $\mu$-space are identical to those in Fig. 3(A). The stable region shown in the $p$-$D$ plane (hatched region) was calculated using the mean of the distribution of $\Delta$ values for each $\mu$-cluster. The left and right panels in the second row show these distributions in $z$-space. The axes of $z$-space are identical to those in Fig. 5(A). Panels from the third row to the fifth row show examples of CoM time-series waveforms (third row), power spectra (fourth row left), DFA plots (fourth row right), SDA plots (fifth row left), and velocity sign-change interval histograms (fifth row right) of representative experimental participants extracted from the corresponding clusters (black: empirical data, red: synthetic data). The bottom-left and bottom-right panels show the boxplots of five clinical scores (Total, Tremor, Rigidity, Bradykinesia, and PIGD scores) among the $\mu$- and $\zeta$-clusters, respectively. *: $p_{bf} < 0.05$, **: $p_{bf} < 0.01$, ***: $p_{bf} < 0.001$, where $p_{bf}$ denotes the $p$-value of the Mann-Whitney U test with Bonferroni correction.

$(\boldsymbol{\mu}_3 \Leftrightarrow)\boldsymbol{\zeta}_2$

**Destination of surrogate data** Of the surrogate data of $\boldsymbol{\zeta}_2$, 25.1 % were mapped near $\boldsymbol{\mu}_3$. Conversely, 24.7 % of the surrogate data of $\boldsymbol{\mu}_3$ were mapped near $\boldsymbol{\zeta}_2$. $\boldsymbol{\zeta}_2$ distributed in wider range including that of $\boldsymbol{\mu}_3$ (Fig. S9).

**Characteristics of postural control strategies** Simillar to $\boldsymbol{\mu}_3$, $\boldsymbol{\zeta}_2$ utilized intermittent control which switches unstable ON and OFF subsystems. But some data shows different control policies (e.g., like $\boldsymbol{\mu}_5$).

**Characteristics of postural sway** A characteristic feature of the postural sway waveforms in $\boldsymbol{\zeta}_2$ is that CoM = 0 mm does not correspond to the upright position (equilibrium position), with sway being frequently observed around this off-center position. It is considered that data for which preprocessing failed to properly establish the equilibrium position at CoM = 0 mm due to stochastic factors—such as postural sway leaning toward one side relative to the equilibrium position or the equilibrium position shifting during measurement—were gathered in $\boldsymbol{\zeta}_2$. The sparse distribution of $\boldsymbol{\zeta}_2$ in the $\boldsymbol{\mu}$-space may be because such postural sway occurred stochastically from a variety of control policies and formed a cluster within the $\boldsymbol{z}$-space.

**UPDRS characteristics** $\boldsymbol{\zeta}_2$ showed low-to-moderate clinical scores.

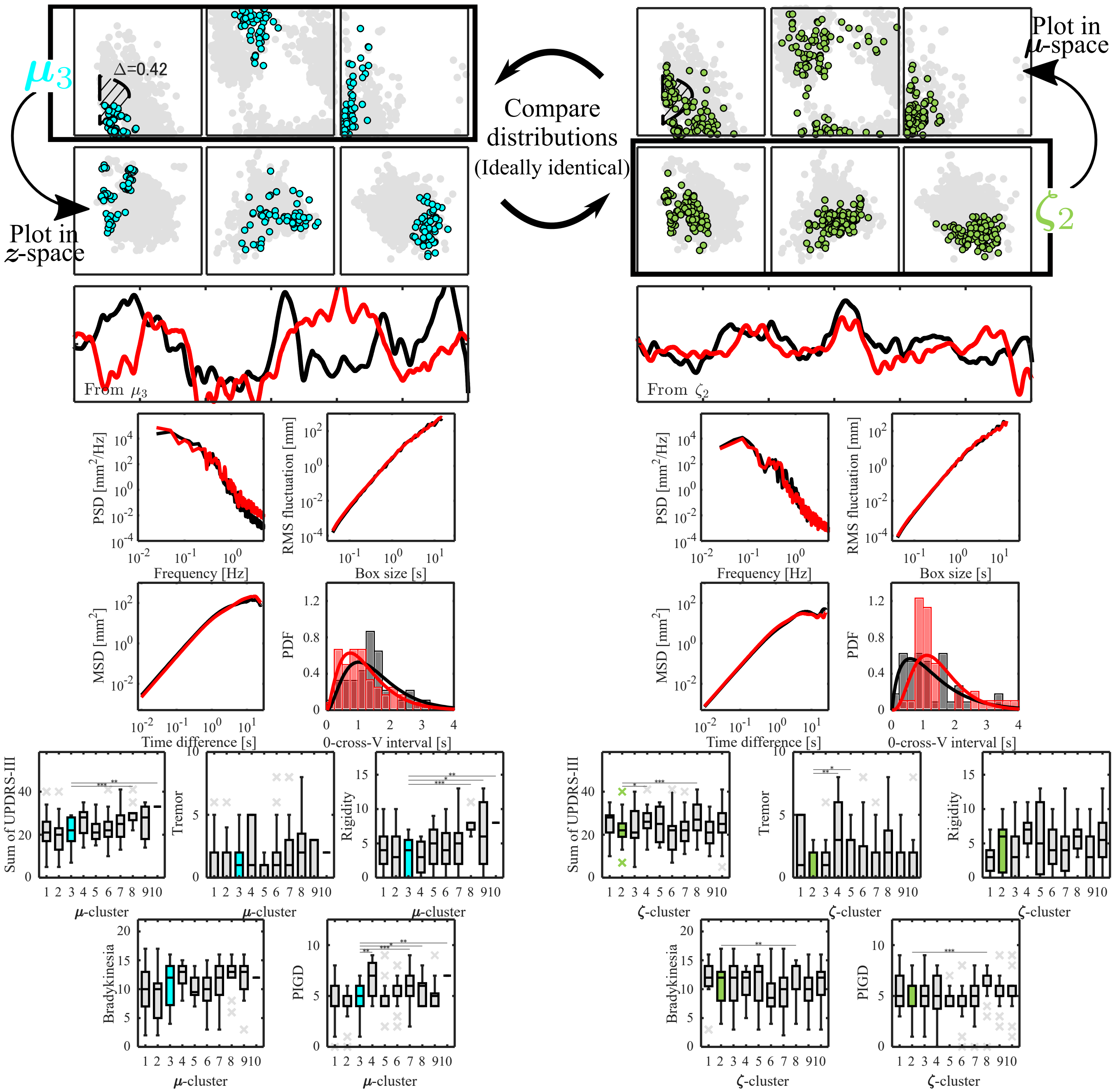


**Fig. S9.** The characteristics of $\zeta_2$. The top-left and top-right panels show the distributions in $\boldsymbol{\mu}$-space for the $\boldsymbol{\mu}$-cluster and $\boldsymbol{\zeta}$-cluster, respectively. The axes of $\boldsymbol{\mu}$-space are identical to those in Fig. 3(A). The stable region shown in the $p$-$D$ plane (hatched region) was calculated using the mean of the distribution of $\Delta$ values for each $\boldsymbol{\mu}$-cluster. The left and right panels in the second row show these distributions in $\boldsymbol{z}$-space. The axes of $\boldsymbol{z}$-space are identical to those in Fig. 5(A). Panels from the third row to the fifth row show examples of CoM time-series waveforms (third row), power spectra (fourth row left), DFA plots (fourth row right), SDA plots (fifth row left), and velocity sign-change interval histograms (fifth row right) of representative experimental participants extracted from the corresponding clusters (black: empirical data, red: synthetic data). The bottom-left and bottom-right panels show the boxplots of five clinical scores (Total, Tremor, Rigidity, Bradykinesia, and PIGD scores) among the $\boldsymbol{\mu}$- and $\boldsymbol{\zeta}$-clusters, respectively. *: $p_{bf} < 0.05$, **: $p_{bf} < 0.01$, ***: $p_{bf} < 0.001$, where $p_{bf}$ denotes the $p$-value of the Mann-Whitney U test with Bonferroni correction.

$((\boldsymbol{\mu}_2, \boldsymbol{\mu}_6) \Leftrightarrow) \boldsymbol{\zeta}_6$

**Destination of surrogate data** Of the surrogate data of $\boldsymbol{\zeta}_6$, 41.3 % were mapped near $\boldsymbol{\mu}_2$ and 12.2 % near $\boldsymbol{\mu}_6$. Conversely, 9.6 % of the surrogate data of $\boldsymbol{\mu}_2$ and 16.7 % of those of $\boldsymbol{\mu}_6$ were mapped near $\boldsymbol{\zeta}_6$.

**Characteristics of postural control strategies** Regarding the postural control policy, similar to the $(\boldsymbol{\mu}_2, \boldsymbol{\mu}_6)$–$\boldsymbol{\zeta}_7$ cluster pair, data points employing intermittent control and those utilizing continuous control are intermingled. Regarding the magnitude of control gains in each data point, approximately half possessed $p$ and $D$ values outside the stability region of the inverted pendulum assuming continuous control, whereas the remaining half were within the stability region.

**Characteristics of postural sway** Regarding the postural sway features, $z_1$, $z_2$, and $z_3$ were small, $z_4$ and $z_5$ were large, and $z_6$ had a value around zero. That is, similar to the $\boldsymbol{\mu}_1$–$\boldsymbol{\zeta}_3$ cluster pair, the low-frequency components are strongly dominant, the high-frequency components are small, and the scaling exponents in both the low- and high-frequency bands are large. On the other hand, unlike the $\boldsymbol{\mu}_1$–$\boldsymbol{\zeta}_3$ cluster pair, $z_2$ is small. Although the sway area across the overall time series is not particularly small due to the large low-frequency components, the short-term sway amplitude is small, resulting in a shorter total path length. Specifically, the body (modeled as an inverted single pendulum) shows a strong tendency to remain within either of the two regions in front of or behind the upright position. In other words, when considering the state space of the pendulum, the state point strongly tends to trace fan-shaped trajectories without crossing $\theta = 0$.

**UPDRS characteristics** Regarding the motor severity of PD patients, $\boldsymbol{\zeta}_6$ exhibits low to moderate clinical score values, influenced by the high proportion of data points belonging to $\boldsymbol{\mu}_1$ and $\boldsymbol{\mu}_2$, which correspond to milder cases with intermittent control.

**Remarks** Simmilar to the $(\boldsymbol{\mu}_2, \boldsymbol{\mu}_6) \Leftrightarrow \boldsymbol{\zeta}_7$ pair, the result from $\boldsymbol{\zeta}_6$ also suggest that severity diagnosis using simple postural sway metrics (such as sway area) is highly likely to overlook severely affected PD patients. The proposed DT can deal with this risk.

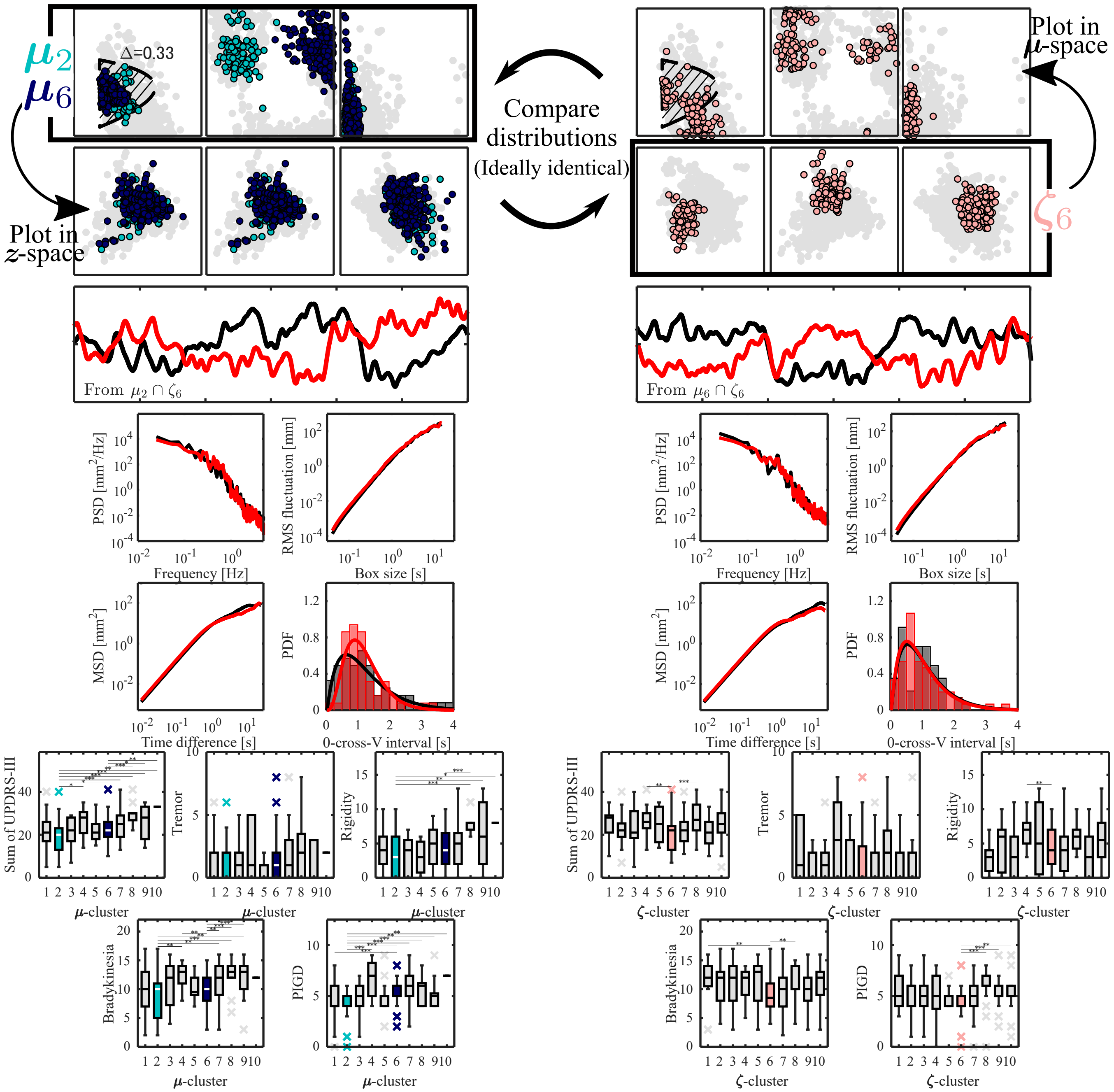


**Fig. S10.** The characteristics of $\zeta_6$. The top-left and top-right panels show the distributions in $\boldsymbol{\mu}$-space for the $\boldsymbol{\mu}$-cluster and $\boldsymbol{\zeta}$-cluster, respectively. The axes of $\boldsymbol{\mu}$-space are identical to those in Fig. 3(A). The stable region shown in the $p$-$D$ plane (hatched region) was calculated using the mean of the distribution of $\Delta$ values for each $\boldsymbol{\mu}$-cluster. The left and right panels in the second row show these distributions in $\boldsymbol{z}$-space. The axes of $\boldsymbol{z}$-space are identical to those in Fig. 5(A). Panels from the third row to the fifth row show examples of CoM time-series waveforms (third row), power spectra (fourth row left), DFA plots (fourth row right), SDA plots (fifth row left), and velocity sign-change interval histograms (fifth row right) of representative experimental participants extracted from the corresponding clusters (black: empirical data, red: synthetic data). The bottom-left and bottom-right panels show the boxplots of five clinical scores (Total, Tremor, Rigidity, Bradykinesia, and PIGD scores) among the $\boldsymbol{\mu}$- and $\boldsymbol{\zeta}$-clusters, respectively. *: $p_{bf} < 0.05$, **: $p_{bf} < 0.01$, ***: $p_{bf} < 0.001$, where $p_{bf}$ denotes the $p$-value of the Mann-Whitney U test with Bonferroni correction.

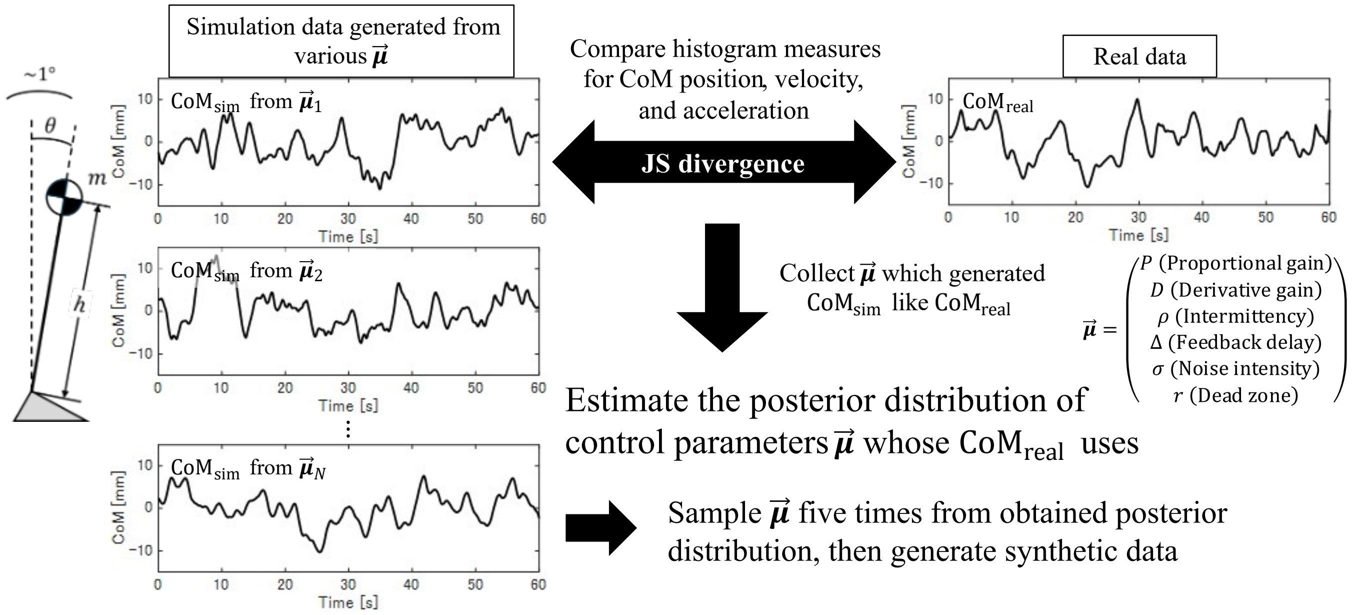


**Fig. S11.** An overview of data assimilation framework

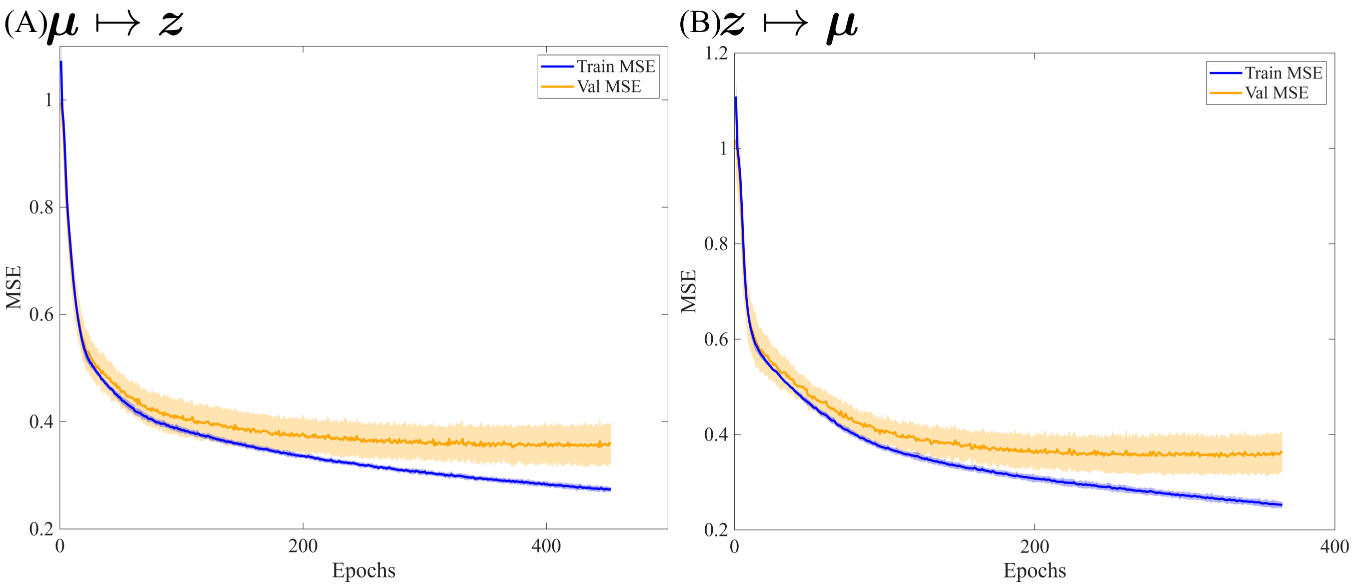


**Fig. S12.** Learning curves of the MSE obtained from 9-fold cross-validation for the three-layer neural network. (A) $\boldsymbol{\mu} \mapsto \boldsymbol{z}$, (B) $\boldsymbol{z} \mapsto \boldsymbol{\mu}$. Solid lines indicate the mean across folds, and the shaded regions represent the mean $\pm$ standard deviation. Blue denotes the MSE for the training data, and orange denotes the MSE for the validation data. The range of epochs shown in the learning curves is aligned with the fold in which training terminated at the earliest epoch due to early stopping.

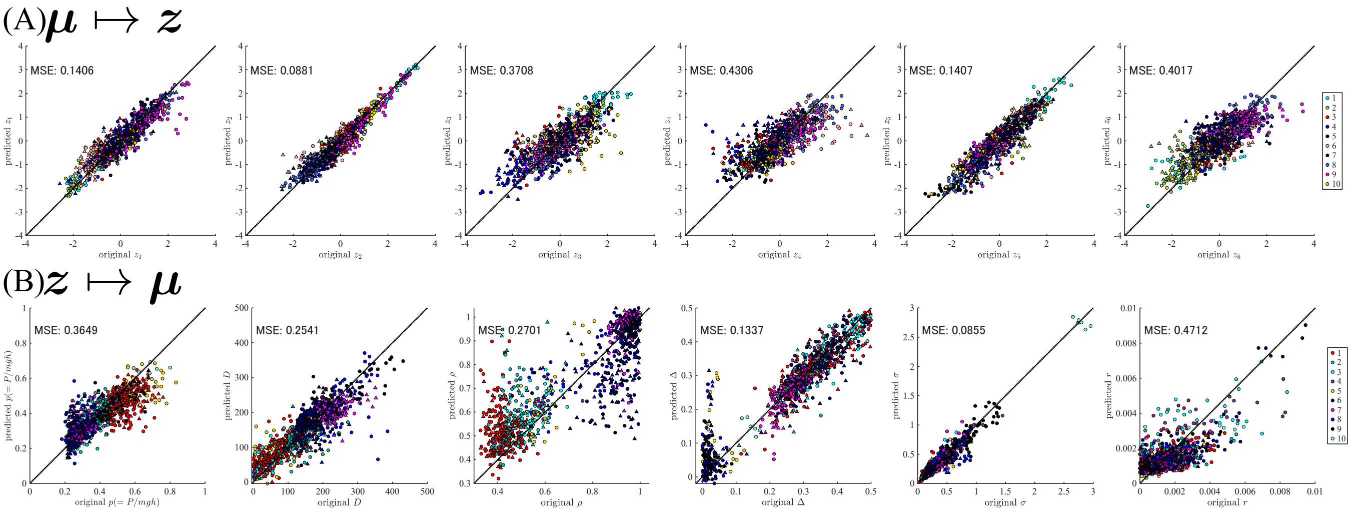


**Fig. S13.** Scatter plots of the true versus predicted values for the trained three-layer neural networks. The upper panels (A) show the network performing $\boldsymbol{\mu} \mapsto \boldsymbol{z}$, with colors indicating $\boldsymbol{\zeta}$-clusters. The lower panels (B) show the network performing $\boldsymbol{z} \mapsto \boldsymbol{\mu}$, with colors indicating $\boldsymbol{\mu}$-clusters. Circles represent patients with Parkinson's disease (PD), and triangles represent healthy older adults.

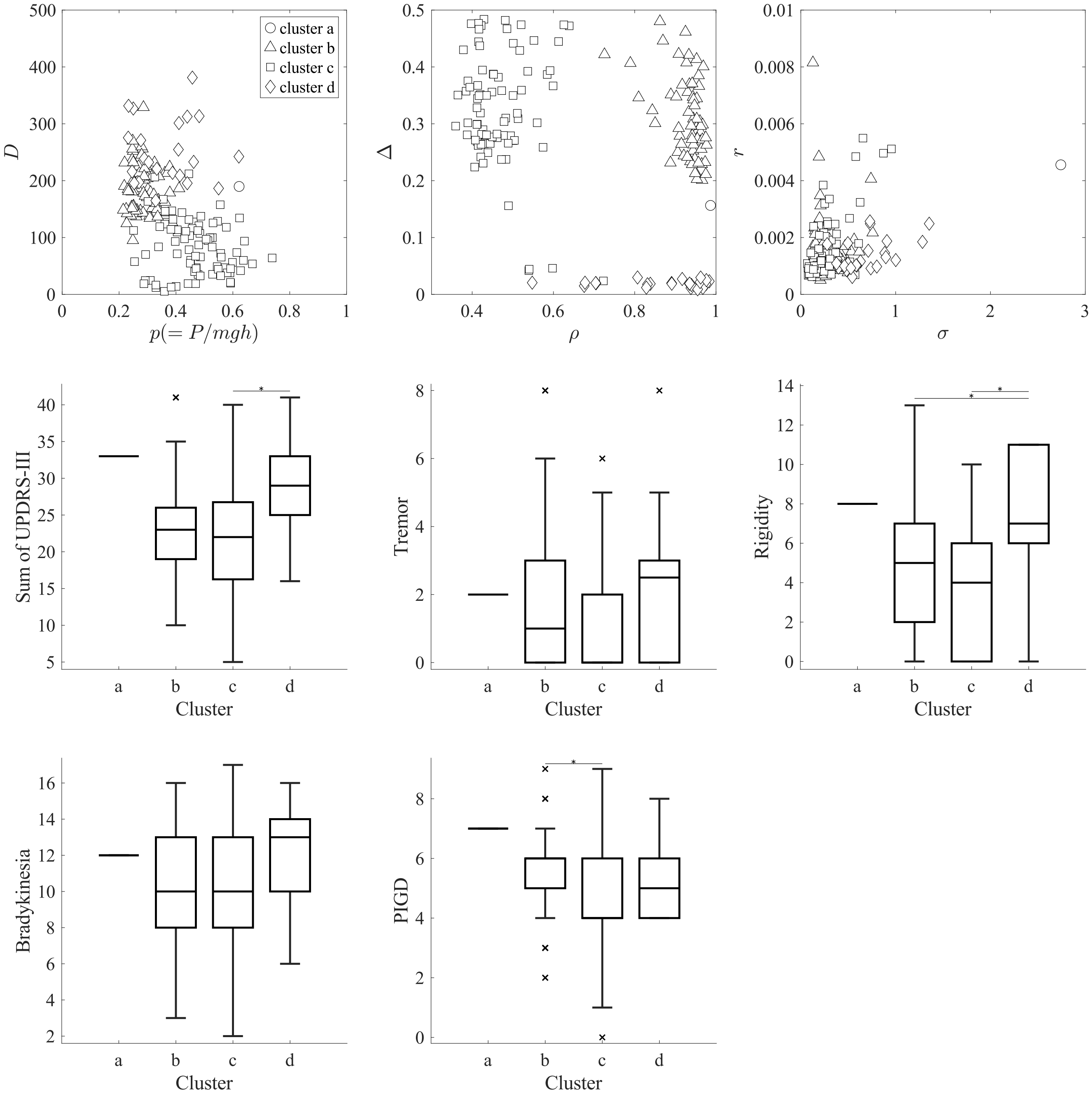


**Fig. S14.** Scatter plot of the real data in $\mu$-space (top), and comparison of clinical scores for each cluster obtained by clustering using only the real data in $\mu$-space (bottom). Cluster a contained only a single data point and was therefore excluded from multiple comparisons. *: $p_{bf} < 0.05$, $p_{bf}$: $p$-value from Mann-Whitney U test with Bonferroni correction applied.

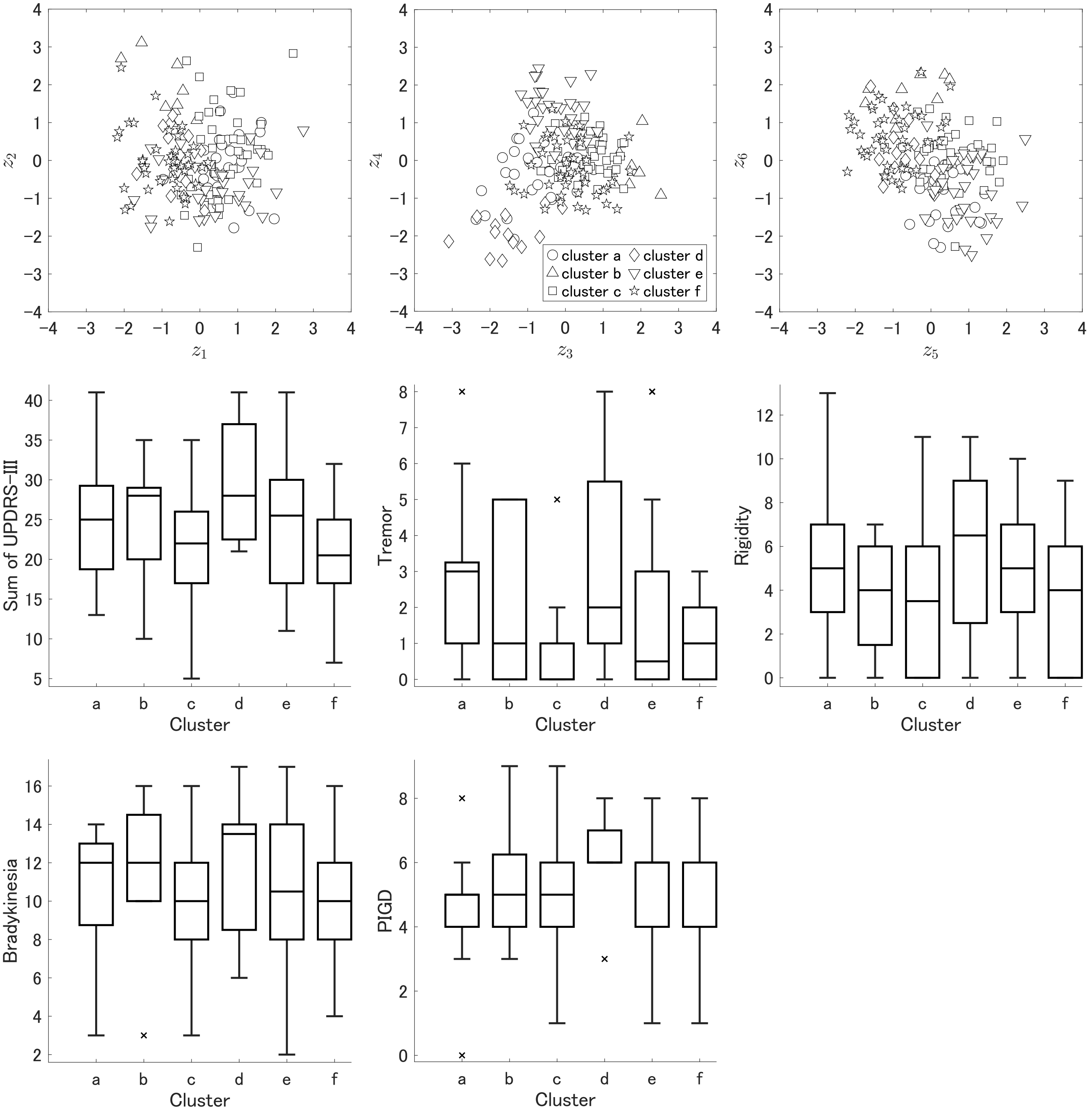


**Fig. S15.** Scatter plot of the real data in $z$-space (top), and comparison of clinical scores for each cluster obtained by clustering using only the real data in $z$-space (bottom).

**Table S1. Details of $\mu$-clusters and $\zeta$-clusters (count for real data)**

| | $\mu_1$ PD/HE | $\mu_2$ PD/HE | $\mu_3$ PD/HE | $\mu_4$ PD/HE | $\mu_5$ PD/HE | $\mu_6$ PD/HE | $\mu_7$ PD/HE | $\mu_8$ PD/HE | $\mu_9$ PD/HE | $\mu_{10}$ PD/HE | SUM PD/HE |
|---|---|---|---|---|---|---|---|---|---|---|---|
| $\zeta_3$ | 2 / 3 | 1 / 0 | 0 / 0 | 0 / 0 | 1 / 0 | 2 / 1 | 0 / 1 | 1 / 0 | 0 / 0 | 0 / 0 | 7 / 5 |
| $\zeta_7$ | 4 / 2 | 4 / 3 | 0 / 0 | 0 / 0 | 1 / 0 | 16 / 6 | 3 / 1 | 0 / 1 | 0 / 2 | 0 / 0 | 28 / 15 |
| $\zeta_1$ | 1 / 0 | 1 / 0 | 5 / 0 | 1 / 0 | 0 / 0 | 1 / 0 | 0 / 0 | 0 / 0 | 0 / 0 | 0 / 0 | 9 / 0 |
| $\zeta_2$ | 10 / 6 | 2 / 3 | 5 / 0 | 2 / 0 | 1 / 0 | 0 / 2 | 0 / 0 | 2 / 3 | 0 / 0 | 0 / 0 | 22 / 14 |
| $\zeta_{10}$ | 0 / 0 | 0 / 0 | 0 / 0 | 0 / 0 | 1 / 0 | 0 / 0 | 0 / 0 | 0 / 0 | 0 / 0 | 0 / 0 | 1 / 0 |
| $\zeta_8$ | 1 / 0 | 0 / 0 | 0 / 0 | 0 / 0 | 0 / 0 | 4 / 1 | 4 / 1 | 2 / 0 | 1 / 1 | 0 / 0 | 12 / 3 |
| $\zeta_9$ | 1 / 0 | 2 / 0 | 0 / 0 | 1 / 0 | 0 / 0 | 4 / 0 | 6 / 2 | 0 / 0 | 1 / 0 | 1 / 0 | 16 / 2 |
| $\zeta_4$ | 0 / 2 | 0 / 0 | 0 / 0 | 0 / 0 | 0 / 0 | 1 / 0 | 3 / 3 | 1 / 2 | 0 / 0 | 0 / 0 | 5 / 7 |
| $\zeta_5$ | 0 / 0 | 0 / 0 | 0 / 0 | 0 / 0 | 0 / 0 | 0 / 0 | 1 / 0 | 0 / 0 | 4 / 0 | 0 / 0 | 5 / 0 |
| $\zeta_6$ | 8 / 3 | 3 / 0 | 1 / 0 | 0 / 0 | 0 / 0 | 1 / 4 | 0 / 0 | 2 / 0 | 0 / 0 | 0 / 0 | 15 / 7 |
| SUM | 27 / 16 | 13 / 6 | 11 / 0 | 4 / 0 | 4 / 0 | 29 / 14 | 17 / 8 | 8 / 6 | 6 / 3 | 1 / 0 | 120 / 53 |

Rows represent $\boldsymbol{\zeta}$-clusters and columns represent $\boldsymbol{\mu}$-clusters. Values to the left and right of the slash indicate the numbers of real data for patients with Parkinson's disease (PD) and healthy older adults, respectively, in the corresponding cluster pair.

**Table S2. Details of $\mu$-clusters and $\zeta$-clusters (including synthetic data)**

| | $\mu_1$ PD/HE | $\mu_2$ PD/HE | $\mu_3$ PD/HE | $\mu_4$ PD/HE | $\mu_5$ PD/HE | $\mu_6$ PD/HE | $\mu_7$ PD/HE | $\mu_8$ PD/HE | $\mu_9$ PD/HE | $\mu_{10}$ PD/HE | SUM PD/HE |
|---|---|---|---|---|---|---|---|---|---|---|---|
| $\zeta_3$ | 33 / 30 | 12 / 3 | 1 / 0 | 3 / 0 | 1 / 0 | 5 / 2 | 1 / 1 | 1 / 0 | 0 / 0 | 0 / 0 | 57 / 36 |
| $\zeta_7$ | 36 / 28 | 31 / 19 | 0 / 0 | 0 / 0 | 7 / 0 | 85 / 37 | 7 / 2 | 2 / 6 | 2 / 8 | 0 / 0 | 170 / 100 |
| $\zeta_1$ | 9 / 0 | 1 / 0 | 25 / 0 | 13 / 0 | 0 / 0 | 7 / 0 | 0 / 0 | 0 / 0 | 0 / 0 | 0 / 0 | 55 / 0 |
| $\zeta_2$ | 14 / 17 | 6 / 6 | 24 / 1 | 2 / 0 | 6 / 0 | 3 / 7 | 0 / 0 | 7 / 13 | 3 / 0 | 0 / 0 | 65 / 44 |
| $\zeta_{10}$ | 6 / 0 | 2 / 0 | 0 / 0 | 1 / 0 | 18 / 0 | 0 / 1 | 0 / 0 | 10 / 0 | 7 / 0 | 0 / 0 | 44 / 1 |
| $\zeta_8$ | 2 / 0 | 4 / 0 | 0 / 0 | 0 / 0 | 0 / 0 | 23 / 6 | 29 / 7 | 9 / 0 | 4 / 6 | 0 / 0 | 71 / 19 |
| $\zeta_9$ | 5 / 0 | 9 / 0 | 0 / 0 | 6 / 0 | 0 / 0 | 34 / 4 | 48 / 19 | 0 / 0 | 2 / 0 | 6 / 0 | 110 / 23 |
| $\zeta_4$ | 3 / 10 | 2 / 2 | 0 / 0 | 0 / 0 | 0 / 0 | 7 / 1 | 9 / 16 | 15 / 16 | 1 / 0 | 0 / 0 | 37 / 45 |
| $\zeta_5$ | 0 / 0 | 0 / 0 | 0 / 0 | 0 / 0 | 0 / 0 | 0 / 2 | 2 / 1 | 3 / 1 | 34 / 5 | 0 / 0 | 39 / 9 |
| $\zeta_6$ | 42 / 13 | 15 / 4 | 5 / 0 | 0 / 0 | 0 / 0 | 8 / 24 | 0 / 0 | 2 / 0 | 0 / 0 | 0 / 0 | 72 / 41 |
| SUM | 150 / 98 | 82 / 34 | 55 / 1 | 25 / 0 | 32 / 0 | 172 / 84 | 96 / 46 | 49 / 36 | 53 / 19 | 6 / 0 | 720 / 318 |

Rows represent $\boldsymbol{\zeta}$-clusters and columns represent $\boldsymbol{\mu}$-clusters. Values to the left and right of the slash indicate the numbers of patients with Parkinson's disease (PD) and healthy older adults, respectively, in the corresponding cluster pair (including both of real and synthetic data).

**Table S3. Results of the Kruskal-Wallis test for UPDRS scores among the $\mu$-clusters**

| | $H$ | $p$ |
|---|---|---|
| Total | 105.96 | $p < 0.001$ |
| Tremor | 29.30 | $p < 0.001$ |
| Rigidity | 104.96 | $p < 0.001$ |
| Bradykinesia | 63.66 | $p < 0.001$ |
| PIGD | 109.34 | $p < 0.001$ |

**Table S4. Comparison of Total scores among the $\mu$-clusters**

| | | $\boldsymbol{\mu}_2$ | $\boldsymbol{\mu}_3$ | $\boldsymbol{\mu}_4$ | $\boldsymbol{\mu}_5$ | $\boldsymbol{\mu}_6$ | $\boldsymbol{\mu}_7$ | $\boldsymbol{\mu}_8$ | $\boldsymbol{\mu}_9$ | $\boldsymbol{\mu}_{10}$ |
|---|---|---|---|---|---|---|---|---|---|---|
| | $U$ | | | | | | 5189.5 | 1208.5 | 2654.0 | 78.0 |
| $\boldsymbol{\mu}_1$ | $p_{bf}$ | n.s. | n.s. | n.s. | n.s. | n.s. | 0.010 | $< 0.001$ | 0.015 | 0.029 |
| | $\|\delta\|$ | | | | | | 0.279 | 0.671 | 0.332 | 0.827 |
| | $U$ | | | 523.5 | | 5240.0 | 2196.0 | 465.5 | 1066.5 | 12.0 |
| $\boldsymbol{\mu}_2$ | $p_{bf}$ | — | n.s. | 0.010 | n.s. | 0.044 | $< 0.001$ | $< 0.001$ | $< 0.001$ | 0.005 |
| | $\|\delta\|$ | | | 0.489 | | 0.257 | 0.442 | 0.768 | 0.509 | 0.951 |
| | $U$ | | | | | | | 437.5 | | 0.0 |
| $\boldsymbol{\mu}_3$ | $p_{bf}$ | — | — | n.s. | n.s. | n.s. | n.s. | $< 0.001$ | n.s. | 0.003 |
| | $\|\delta\|$ | | | | | | | 0.675 | | 1.000 |
| | $U$ | | | | | | | | | |
| $\boldsymbol{\mu}_4$ | $p_{bf}$ | — | — | — | n.s. | n.s. | n.s. | n.s. | n.s. | n.s. |
| | $\|\delta\|$ | | | | | | | | | |
| | $U$ | | | | | | | 286.5 | | 6.0 |
| $\boldsymbol{\mu}_5$ | $p_{bf}$ | — | — | — | — | n.s. | n.s. | $< 0.001$ | n.s. | 0.014 |
| | $\|\delta\|$ | | | | | | | 0.635 | | 0.938 |
| | $U$ | | | | | | | 1327.0 | 3150.5 | 54.0 |
| $\boldsymbol{\mu}_6$ | $p_{bf}$ | — | — | — | — | — | n.s. | $< 0.001$ | 0.032 | 0.009 |
| | $\|\delta\|$ | | | | | | | 0.685 | 0.309 | 0.895 |
| | $U$ | | | | | | | 1335.0 | | |
| $\boldsymbol{\mu}_7$ | $p_{bf}$ | — | — | — | — | — | — | $< 0.001$ | n.s. | n.s. |
| | $\|\delta\|$ | | | | | | | 0.432 | | |
| | $U$ | | | | | | | | | |
| $\boldsymbol{\mu}_8$ | $p_{bf}$ | — | — | — | — | — | — | — | n.s. | n.s. |
| | $\|\delta\|$ | | | | | | | | | |
| | $U$ | | | | | | | | | |
| $\boldsymbol{\mu}_9$ | $p_{bf}$ | — | — | — | — | — | — | — | — | n.s. |
| | $\|\delta\|$ | | | | | | | | | |

Rows and columns correspond to each $\boldsymbol{\mu}$-cluster. n.s.: not significant. $U$: test statistic of the Mann-Whitney U test. $p_{bf}$: $p$-value after Bonferroni correction. $\delta$: Cliff's $\delta$.

**Table S5. Three group comparison of Total scores among the $\mu$-clusters**

| | | $\boldsymbol{M}_{\text{CONT}}$ ($\boldsymbol{\mu}_6 \cup \boldsymbol{\mu}_7$) | $\boldsymbol{M}_{\text{STIFF}}$ ($\boldsymbol{\mu}_8 \cup \boldsymbol{\mu}_9$) |
|---|---|---|---|
| | $U$ | 24560 | 5394.5 |
| $\boldsymbol{M}_{\text{INT}}$ | $p_{bf}$ | 0.002 | $< 0.001$ |
| ($\boldsymbol{\mu}_1 \cup \boldsymbol{\mu}_2$) | $\|\delta\|$ | 0.210 | 0.544 |
| | $U$ | | 8064.5 |
| $\boldsymbol{M}_{\text{CONT}}$ | $p_{bf}$ | — | $< 0.001$ |
| ($\boldsymbol{\mu}_6 \cup \boldsymbol{\mu}_7$) | $\|\delta\|$ | | 0.410 |

Rows and columns correspond to each $\boldsymbol{\mu}$-cluster. n.s.: not significant. $U$: test statistic of the Mann-Whitney U test. $p_{bf}$: $p$-value after Bonferroni correction. $\delta$: Cliff's $\delta$.

**Table S6. Comparison of Tremor scores among the $\mu$-clusters**

| | | $\mu_2$ | $\mu_3$ | $\mu_4$ | $\mu_5$ | $\mu_6$ | $\mu_7$ | $\mu_8$ | $\mu_9$ | $\mu_{10}$ |
|---|---|---|---|---|---|---|---|---|---|---|
| $\mu_1$ | $U$<br>$p_{bf}$<br>$\|\delta\|$ | n.s. | n.s. | n.s. | n.s. | n.s. | n.s. | 2495.0<br>0.011<br>0.321 | n.s. | n.s. |
| $\mu_2$ | $U$<br>$p_{bf}$<br>$\|\delta\|$ | — | n.s. | n.s. | n.s. | n.s. | n.s. | n.s. | n.s. | n.s. |
| $\mu_3$ | $U$<br>$p_{bf}$<br>$\|\delta\|$ | — | — | n.s. | n.s. | n.s. | n.s. | n.s. | n.s. | n.s. |
| $\mu_4$ | $U$<br>$p_{bf}$<br>$\|\delta\|$ | — | — | — | n.s. | n.s. | n.s. | n.s. | n.s. | n.s. |
| $\mu_5$ | $U$<br>$p_{bf}$<br>$\|\delta\|$ | — | — | — | — | n.s. | n.s. | n.s. | n.s. | 18.0<br>0.046<br>0.812 |
| $\mu_6$ | $U$<br>$p_{bf}$<br>$\|\delta\|$ | — | — | — | — | — | n.s. | n.s. | n.s. | n.s. |
| $\mu_7$ | $U$<br>$p_{bf}$<br>$\|\delta\|$ | — | — | — | — | — | — | n.s. | n.s. | n.s. |
| $\mu_8$ | $U$<br>$p_{bf}$<br>$\|\delta\|$ | — | — | — | — | — | — | — | n.s. | n.s. |
| $\mu_9$ | $U$<br>$p_{bf}$<br>$\|\delta\|$ | — | — | — | — | — | — | — | — | n.s. |

Rows and columns correspond to each $\mu$-cluster. n.s.: not significant. $U$: test statistic of the Mann-Whitney U test. $p_{bf}$: $p$-value after Bonferroni correction. $\delta$: Cliff's $\delta$.

**Table S7. Three group comparison of Tremor scores among the $\mu$-clusters**

| | | $M_{\mathrm{CONT}}$ ($\mu_6 \cup \mu_7$) | $M_{\mathrm{STIFF}}$ ($\mu_8 \cup \mu_9$) |
|---|---|---|---|
| $M_{\mathrm{INT}}$<br>($\mu_1 \cup \mu_2$) | $U$<br>$p_{bf}$<br>$\|\delta\|$ | n.s. | 8586.0<br>$< 0.001$<br>0.274 |
| $M_{\mathrm{CONT}}$<br>($\mu_6 \cup \mu_7$) | $U$<br>$p_{bf}$<br>$\|\delta\|$ | — | n.s. |

Rows and columns correspond to each $\mu$-cluster. n.s.: not significant. $U$: test statistic of the Mann-Whitney U test. $p_{bf}$: $p$-value after Bonferroni correction. $\delta$: Cliff's $\delta$.

**Table S8. Comparison of Rigidity scores among the $\mu$-clusters**

| | | $\boldsymbol{\mu}_2$ | $\boldsymbol{\mu}_3$ | $\boldsymbol{\mu}_4$ | $\boldsymbol{\mu}_5$ | $\boldsymbol{\mu}_6$ | $\boldsymbol{\mu}_7$ | $\boldsymbol{\mu}_8$ | $\boldsymbol{\mu}_9$ | $\boldsymbol{\mu}_{10}$ |
|---|---|---|---|---|---|---|---|---|---|---|
| $\boldsymbol{\mu}_1$ | $U$ | | | | | | | 1113.0 | | |
| | $p_{bf}$ | n.s. | n.s. | n.s. | n.s. | n.s. | n.s. | $< 0.001$ | n.s. | n.s. |
| | $\|\delta\|$ | | | | | | | 0.697 | | |
| $\boldsymbol{\mu}_2$ | $U$ | | | | | | | 482.5 | 1313.5 | 36.0 |
| | $p_{bf}$ | — | n.s. | n.s. | n.s. | n.s. | n.s. | $< 0.001$ | 0.004 | 0.016 |
| | $\|\delta\|$ | | | | | | | 0.760 | 0.396 | 0.854 |
| $\boldsymbol{\mu}_3$ | $U$ | | | | | | | 207.5 | 881.0 | 0.0 |
| | $p_{bf}$ | — | — | n.s. | n.s. | n.s. | n.s. | $< 0.001$ | 0.016 | 0.002 |
| | $\|\delta\|$ | | | | | | | 0.846 | 0.396 | 1.000 |
| $\boldsymbol{\mu}_4$ | $U$ | | | | | | | 33.0 | | 0.0 |
| | $p_{bf}$ | — | — | — | n.s. | n.s. | n.s. | $< 0.001$ | n.s. | 0.007 |
| | $\|\delta\|$ | | | | | | | 0.946 | | 1.000 |
| $\boldsymbol{\mu}_5$ | $U$ | | | | | | | 296.0 | | |
| | $p_{bf}$ | — | — | — | — | n.s. | n.s. | $< 0.001$ | n.s. | n.s. |
| | $\|\delta\|$ | | | | | | | 0.622 | | |
| $\boldsymbol{\mu}_6$ | $U$ | | | | | | | 1300.0 | | |
| | $p_{bf}$ | — | — | — | — | — | n.s. | $< 0.001$ | n.s. | n.s. |
| | $\|\delta\|$ | | | | | | | 0.692 | | |
| $\boldsymbol{\mu}_7$ | $U$ | | | | | | | 683.0 | | 42.0 |
| | $p_{bf}$ | — | — | — | — | — | — | $< 0.001$ | n.s. | 0.020 |
| | $\|\delta\|$ | | | | | | | 0.710 | | 0.854 |
| $\boldsymbol{\mu}_8$ | $U$ | | | | | | | | | |
| | $p_{bf}$ | — | — | — | — | — | — | — | n.s. | n.s. |
| | $\|\delta\|$ | | | | | | | | | |
| $\boldsymbol{\mu}_9$ | $U$ | | | | | | | | | |
| | $p_{bf}$ | — | — | — | — | — | — | — | — | n.s. |
| | $\|\delta\|$ | | | | | | | | | |

Rows and columns correspond to each $\boldsymbol{\mu}$-cluster. n.s.: not significant. $U$: test statistic of the Mann-Whitney U test. $p_{bf}$: $p$-value after Bonferroni correction. $\delta$: Cliff's $\delta$.

**Table S9. Three group comparison of Rigidity scores among the $\mu$-clusters**

| | | $\boldsymbol{M}_{\text{CONT}}$ ($\boldsymbol{\mu}_6 \cup \boldsymbol{\mu}_7$) | $\boldsymbol{M}_{\text{STIFF}}$ ($\boldsymbol{\mu}_8 \cup \boldsymbol{\mu}_9$) |
|---|---|---|---|
| $\boldsymbol{M}_{\text{INT}}$ ($\boldsymbol{\mu}_1 \cup \boldsymbol{\mu}_2$) | $U$ | | 5768.0 |
| | $p_{bf}$ | n.s. | $< 0.001$ |
| | $\|\delta\|$ | | 0.513 |
| $\boldsymbol{M}_{\text{CONT}}$ ($\boldsymbol{\mu}_6 \cup \boldsymbol{\mu}_7$) | $U$ | | 7204.5 |
| | $p_{bf}$ | — | $< 0.001$ |
| | $\|\delta\|$ | | 0.473 |

Rows and columns correspond to each $\boldsymbol{\mu}$-cluster. n.s.: not significant. $U$: test statistic of the Mann-Whitney U test. $p_{bf}$: $p$-value after Bonferroni correction. $\delta$: Cliff's $\delta$.

**Table S10. Comparison of Bradykinesia scores among the $\mu$-clusters**

| | | $\boldsymbol{\mu}_2$ | $\boldsymbol{\mu}_3$ | $\boldsymbol{\mu}_4$ | $\boldsymbol{\mu}_5$ | $\boldsymbol{\mu}_6$ | $\boldsymbol{\mu}_7$ | $\boldsymbol{\mu}_8$ | $\boldsymbol{\mu}_9$ | $\boldsymbol{\mu}_{10}$ |
|---|---|---|---|---|---|---|---|---|---|---|
| $\boldsymbol{\mu}_1$ | $U$<br>$p_{bf}$<br>$\|\delta\|$ | n.s. | n.s. | n.s. | n.s. | n.s. | n.s. | 2416.5<br>0.014<br>0.342 | 2648.0<br>0.013<br>0.334 | n.s. |
| $\boldsymbol{\mu}_2$ | $U$<br>$p_{bf}$<br>$\|\delta\|$ | — | n.s. | 534.5<br>0.013<br>0.479 | n.s. | n.s. | 2643.0<br>0.007<br>0.329 | 1058.0<br>$< 0.001$<br>0.473 | 1271.0<br>0.002<br>0.415 | n.s. |
| $\boldsymbol{\mu}_3$ | $U$<br>$p_{bf}$<br>$\|\delta\|$ | — | — | n.s. | n.s. | n.s. | n.s. | n.s. | n.s. | n.s. |
| $\boldsymbol{\mu}_4$ | $U$<br>$p_{bf}$<br>$\|\delta\|$ | — | — | — | n.s. | 1140.0<br>0.006<br>0.470 | n.s. | n.s. | n.s. | n.s. |
| $\boldsymbol{\mu}_5$ | $U$<br>$p_{bf}$<br>$\|\delta\|$ | — | — | — | — | n.s. | n.s. | 418.0<br>0.017<br>0.467 | 443.5<br>0.010<br>0.477 | n.s. |
| $\boldsymbol{\mu}_6$ | $U$<br>$p_{bf}$<br>$\|\delta\|$ | — | — | — | — | — | 5863.0<br>0.004<br>0.290 | 2459.5<br>$< 0.001$<br>0.416 | 2752.5<br>$< 0.001$<br>0.396 | n.s. |
| $\boldsymbol{\mu}_7$ | $U$<br>$p_{bf}$<br>$\|\delta\|$ | — | — | — | — | — | — | n.s. | n.s. | n.s. |
| $\boldsymbol{\mu}_8$ | $U$<br>$p_{bf}$<br>$\|\delta\|$ | — | — | — | — | — | — | — | n.s. | n.s. |
| $\boldsymbol{\mu}_9$ | $U$<br>$p_{bf}$<br>$\|\delta\|$ | — | — | — | — | — | — | — | — | n.s. |

Rows and columns correspond to each $\boldsymbol{\mu}$-cluster. n.s.: not significant. $U$: test statistic of the Mann-Whitney U test. $p_{bf}$: $p$-value after Bonferroni correction. $\delta$: Cliff's $\delta$.

**Table S11. Three group comparison of Bradykinesia scores among the $\mu$-clusters**

| | | $\boldsymbol{M}_{\text{CONT}}$ $(\boldsymbol{\mu}_6 \cup \boldsymbol{\mu}_7)$ | $\boldsymbol{M}_{\text{STIFF}}$ $(\boldsymbol{\mu}_8 \cup \boldsymbol{\mu}_9)$ |
|---|---|---|---|
| $\boldsymbol{M}_{\text{INT}}$<br>$(\boldsymbol{\mu}_1 \cup \boldsymbol{\mu}_2)$ | $U$<br>$p_{bf}$<br>$\|\delta\|$ | n.s. | 7393.5<br>$< 0.001$<br>0.375 |
| $\boldsymbol{M}_{\text{CONT}}$<br>$(\boldsymbol{\mu}_6 \cup \boldsymbol{\mu}_7)$ | $U$<br>$p_{bf}$<br>$\|\delta\|$ | — | 9601.5<br>$< 0.001$<br>0.298 |

Rows and columns correspond to each $\boldsymbol{\mu}$-cluster. n.s.: not significant. $U$: test statistic of the Mann-Whitney U test. $p_{bf}$: $p$-value after Bonferroni correction. $\delta$: Cliff's $\delta$.

**Table S12. Comparison of PIGD scores among the $\mu$-clusters**

| | | $\boldsymbol{\mu}_2$ | $\boldsymbol{\mu}_3$ | $\boldsymbol{\mu}_4$ | $\boldsymbol{\mu}_5$ | $\boldsymbol{\mu}_6$ | $\boldsymbol{\mu}_7$ | $\boldsymbol{\mu}_8$ | $\boldsymbol{\mu}_9$ | $\boldsymbol{\mu}_{10}$ |
|---|---|---|---|---|---|---|---|---|---|---|
| | $U$ | | | 853.0 | | 9066.0 | 3968.5 | 2068.0 | | 45.0 |
| $\boldsymbol{\mu}_1$ | $p_{bf}$ | n.s. | n.s. | $< 0.001$ | n.s. | $< 0.001$ | $< 0.001$ | $< 0.001$ | n.s. | 0.006 |
| | $\|\delta\|$ | | | 0.545 | | 0.297 | 0.449 | 0.437 | | 0.900 |
| | $U$ | | | 364.5 | | 4006.0 | 1706.0 | 857.0 | 1364.0 | 0.0 |
| $\boldsymbol{\mu}_2$ | $p_{bf}$ | — | n.s. | $< 0.001$ | n.s. | $< 0.001$ | $< 0.001$ | $< 0.001$ | 0.007 | 0.001 |
| | $\|\delta\|$ | | | 0.644 | | 0.432 | 0.567 | 0.573 | 0.372 | 1.000 |
| | $U$ | | | 331.0 | | | 1538.5 | 830.5 | | 12.0 |
| $\boldsymbol{\mu}_3$ | $p_{bf}$ | — | — | 0.008 | n.s. | n.s. | $< 0.001$ | 0.025 | n.s. | 0.008 |
| | $\|\delta\|$ | | | 0.519 | | | 0.417 | 0.384 | | 0.927 |
| | $U$ | | | | | | | | | |
| $\boldsymbol{\mu}_4$ | $p_{bf}$ | — | — | — | n.s. | n.s. | n.s. | n.s. | n.s. | n.s. |
| | $\|\delta\|$ | | | | | | | | | |
| | $U$ | | | | | | | | | |
| $\boldsymbol{\mu}_5$ | $p_{bf}$ | — | — | — | — | n.s. | n.s. | n.s. | n.s. | n.s. |
| | $\|\delta\|$ | | | | | | | | | |
| | $U$ | | | | | | | | | |
| $\boldsymbol{\mu}_6$ | $p_{bf}$ | — | — | — | — | — | n.s. | n.s. | n.s. | n.s. |
| | $\|\delta\|$ | | | | | | | | | |
| | $U$ | | | | | | | | 1615.5 | |
| $\boldsymbol{\mu}_7$ | $p_{bf}$ | — | — | — | — | — | — | n.s. | 0.008 | n.s. |
| | $\|\delta\|$ | | | | | | | | 0.365 | |
| | $U$ | | | | | | | | | |
| $\boldsymbol{\mu}_8$ | $p_{bf}$ | — | — | — | — | — | — | — | n.s. | n.s. |
| | $\|\delta\|$ | | | | | | | | | |
| | $U$ | | | | | | | | | 15.0 |
| $\boldsymbol{\mu}_9$ | $p_{bf}$ | — | — | — | — | — | — | — | — | 0.008 |
| | $\|\delta\|$ | | | | | | | | | 0.906 |

Rows and columns correspond to each $\boldsymbol{\mu}$-cluster. n.s.: not significant. $U$: test statistic of the Mann-Whitney U test. $p_{bf}$: $p$-value after Bonferroni correction. $\delta$: Cliff's $\delta$.

**Table S13. Three group comparison of PIGD scores among the $\mu$-clusters**

| | | $\boldsymbol{M}_{\text{CONT}}$ ($\boldsymbol{\mu}_6 \cup \boldsymbol{\mu}_7$) | $\boldsymbol{M}_{\text{STIFF}}$ ($\boldsymbol{\mu}_8 \cup \boldsymbol{\mu}_9$) |
|---|---|---|---|
| | $U$ | 18746.5 | 7458.5 |
| $\boldsymbol{M}_{\text{INT}}$ | $p_{bf}$ | $< 0.001$ | $< 0.001$ |
| ($\boldsymbol{\mu}_1 \cup \boldsymbol{\mu}_2$) | $\|\delta\|$ | 0.397 | 0.370 |
| | $U$ | | |
| $\boldsymbol{M}_{\text{CONT}}$ | $p_{bf}$ | — | n.s. |
| ($\boldsymbol{\mu}_6 \cup \boldsymbol{\mu}_7$) | $\|\delta\|$ | | |

Rows and columns correspond to each $\boldsymbol{\mu}$-cluster. n.s.: not significant. $U$: test statistic of the Mann-Whitney U test. $p_{bf}$: $p$-value after Bonferroni correction. $\delta$: Cliff's $\delta$.

**Table S14. Results of the Kruskal-Wallis test for UPDRS scores among the $\zeta$-clusters**

| | $H$ | $p$ |
|---|---|---|
| Total | 68.43 | $p < 0.001$ |
| Tremor | 53.44 | $p < 0.001$ |
| Rigidity | 62.32 | $p < 0.001$ |
| Bradykinesia | 63.66 | $p < 0.001$ |
| PIGD | 77.70 | $p < 0.001$ |

**Table S15. Comparison of Total scores among the $\zeta$-clusters**

| | | $\zeta_2$ | $\zeta_3$ | $\zeta_4$ | $\zeta_5$ | $\zeta_6$ | $\zeta_7$ | $\zeta_8$ | $\zeta_9$ | $\zeta_{10}$ |
|---|---|---|---|---|---|---|---|---|---|---|
| $\zeta_1$ | $U$<br>$p_{bf}$<br>$\|\delta\|$ | n.s. | n.s. | n.s. | n.s. | n.s. | 3211.0<br>0.021<br>0.313 | n.s. | n.s. | n.s. |
| $\zeta_2$ | $U$<br>$p_{bf}$<br>$\|\delta\|$ | — | n.s. | 682.5<br>0.013<br>0.432 | n.s. | n.s. | n.s. | 1241.5<br>$< 0.001$<br>0.462 | n.s. | n.s. |
| $\zeta_3$ | $U$<br>$p_{bf}$<br>$\|\delta\|$ | — | — | n.s. | n.s. | n.s. | n.s. | n.s. | n.s. | n.s. |
| $\zeta_4$ | $U$<br>$p_{bf}$<br>$\|\delta\|$ | — | — | — | n.s. | 718.5<br>0.004<br>0.461 | 1719.0<br>$< 0.001$<br>0.453 | n.s. | 1164.0<br>0.004<br>0.428 | n.s. |
| $\zeta_5$ | $U$<br>$p_{bf}$<br>$\|\delta\|$ | — | — | — | — | n.s. | n.s. | n.s. | n.s. | n.s. |
| $\zeta_6$ | $U$<br>$p_{bf}$<br>$\|\delta\|$ | — | — | — | — | — | n.s. | 1362.5<br>$< 0.001$<br>0.467 | n.s. | n.s. |
| $\zeta_7$ | $U$<br>$p_{bf}$<br>$\|\delta\|$ | — | — | — | — | — | — | 2931.0<br>$< 0.001$<br>0.514 | n.s. | n.s. |
| $\zeta_8$ | $U$<br>$p_{bf}$<br>$\|\delta\|$ | — | — | — | — | — | — | — | 1916.5<br>$< 0.001$<br>0.509 | n.s. |
| $\zeta_9$ | $U$<br>$p_{bf}$<br>$\|\delta\|$ | — | — | — | — | — | — | — | — | n.s. |

Rows and columns correspond to each $\zeta$-cluster. n.s.: not significant. $U$: test statistic of the Mann-Whitney U test. $p_{bf}$: $p$-value after Bonferroni correction. $\delta$: Cliff's $\delta$.

**Table S16. Comparison of Tremor scores among the $\zeta$-clusters**

| | | $\zeta_2$ | $\zeta_3$ | $\zeta_4$ | $\zeta_5$ | $\zeta_6$ | $\zeta_7$ | $\zeta_8$ | $\zeta_9$ | $\zeta_{10}$ |
|---|---|---|---|---|---|---|---|---|---|---|
| | $U$ | | | | | | 3349.0 | | 2090.0 | |
| $\zeta_1$ | $p_{bf}$ | n.s. | n.s. | n.s. | n.s. | n.s. | 0.031 | n.s. | 0.024 | n.s. |
| | $\|\delta\|$ | | | | | | 0.284 | | 0.309 | |
| | $U$ | | | 657.5 | 793.0 | | | | | |
| $\zeta_2$ | $p_{bf}$ | — | n.s. | 0.003 | 0.027 | n.s. | n.s. | n.s. | n.s. | n.s. |
| | $\|\delta\|$ | | | 0.453 | 0.374 | | | | | |
| | $U$ | | | | | | | | | |
| $\zeta_3$ | $p_{bf}$ | — | — | n.s. | n.s. | n.s. | n.s. | n.s. | n.s. | n.s. |
| | $\|\delta\|$ | | | | | | | | | |
| | $U$ | | | | | | 1769.0 | | 1098.0 | 463.5 |
| $\zeta_4$ | $p_{bf}$ | — | — | — | n.s. | n.s. | $< 0.001$ | n.s. | $< 0.001$ | 0.023 |
| | $\|\delta\|$ | | | | | | 0.438 | | 0.460 | 0.431 |
| | $U$ | | | | | | | | 1395.0 | |
| $\zeta_5$ | $p_{bf}$ | — | — | — | — | n.s. | n.s. | n.s. | 0.020 | n.s. |
| | $\|\delta\|$ | | | | | | | | 0.350 | |
| | $U$ | | | | | | | | | |
| $\zeta_6$ | $p_{bf}$ | — | — | — | — | — | n.s. | n.s. | n.s. | n.s. |
| | $\|\delta\|$ | | | | | | | | | |
| | $U$ | | | | | | | 4489.0 | | |
| $\zeta_7$ | $p_{bf}$ | — | — | — | — | — | — | 0.032 | n.s. | n.s. |
| | $\|\delta\|$ | | | | | | | 0.256 | | |
| | $U$ | | | | | | | | 2757.0 | |
| $\zeta_8$ | $p_{bf}$ | — | — | — | — | — | — | — | 0.015 | n.s. |
| | $\|\delta\|$ | | | | | | | | 0.294 | |
| | $U$ | | | | | | | | | |
| $\zeta_9$ | $p_{bf}$ | — | — | — | — | — | — | — | — | n.s. |
| | $\|\delta\|$ | | | | | | | | | |

Rows and columns correspond to each $\zeta$-cluster. n.s.: not significant. $U$: test statistic of the Mann-Whitney U test. $p_{bf}$: $p$-value after Bonferroni correction. $\delta$: Cliff's $\delta$.

**Table S17. Comparison of Rigidity scores among the $\zeta$-clusters**

| | | $\zeta_2$ | $\zeta_3$ | $\zeta_4$ | $\zeta_5$ | $\zeta_6$ | $\zeta_7$ | $\zeta_8$ | $\zeta_9$ | $\zeta_{10}$ |
|---|---|---|---|---|---|---|---|---|---|---|
| | $U$ | | | 287.0 | | | | 782.5 | | |
| $\zeta_1$ | $p_{bf}$ | n.s. | n.s. | $< 0.001$ | n.s. | n.s. | n.s. | $< 0.001$ | n.s. | n.s. |
| | $\|\delta\|$ | | | 0.718 | | | | 0.599 | | |
| | $U$ | | | | | | | | | |
| $\zeta_2$ | $p_{bf}$ | — | n.s. | n.s. | n.s. | n.s. | n.s. | n.s. | n.s. | n.s. |
| | $\|\delta\|$ | | | | | | | | | |
| | $U$ | | | 546.5 | | | | 1248.5 | | |
| $\zeta_3$ | $p_{bf}$ | — | — | 0.003 | n.s. | n.s. | n.s. | 0.008 | n.s. | n.s. |
| | $\|\delta\|$ | | | 0.482 | | | | 0.383 | | |
| | $U$ | | | | | 725.5 | 1696.5 | | 922.5 | |
| $\zeta_4$ | $p_{bf}$ | — | — | — | n.s. | 0.004 | $< 0.001$ | n.s. | $< 0.001$ | n.s. |
| | $\|\delta\|$ | | | | | 0.455 | 0.461 | | 0.547 | |
| | $U$ | | | | | | | | | |
| $\zeta_5$ | $p_{bf}$ | — | — | — | — | n.s. | n.s. | n.s. | n.s. | n.s. |
| | $\|\delta\|$ | | | | | | | | | |
| | $U$ | | | | | | | | | |
| $\zeta_6$ | $p_{bf}$ | — | — | — | — | — | n.s. | n.s. | n.s. | n.s. |
| | $\|\delta\|$ | | | | | | | | | |
| | $U$ | | | | | | | 4016.5 | | |
| $\zeta_7$ | $p_{bf}$ | — | — | — | — | — | — | 0.002 | n.s. | n.s. |
| | $\|\delta\|$ | | | | | | | 0.334 | | |
| | $U$ | | | | | | | | 2170.0 | |
| $\zeta_8$ | $p_{bf}$ | — | — | — | — | — | — | — | $< 0.001$ | n.s. |
| | $\|\delta\|$ | | | | | | | | 0.444 | |
| | $U$ | | | | | | | | | |
| $\zeta_9$ | $p_{bf}$ | — | — | — | — | — | — | — | — | n.s. |
| | $\|\delta\|$ | | | | | | | | | |

Rows and columns correspond to each $\zeta$-cluster. n.s.: not significant. $U$: test statistic of the Mann-Whitney U test. $p_{bf}$: $p$-value after Bonferroni correction. $\delta$: Cliff's $\delta$.

**Table S18. Comparison of Bradykinesia scores among the $\zeta$-clusters**

| | | $\zeta_2$ | $\zeta_3$ | $\zeta_4$ | $\zeta_5$ | $\zeta_6$ | $\zeta_7$ | $\zeta_8$ | $\zeta_9$ | $\zeta_{10}$ |
|---|---|---|---|---|---|---|---|---|---|---|
| | $U$ | | | | | 1113.5 | 2439.0 | | 1871.5 | |
| $\zeta_1$ | $p_{bf}$ | n.s. | n.s. | n.s. | n.s. | 0.001 | $< 0.001$ | n.s. | 0.002 | n.s. |
| | $\lvert\delta\rvert$ | | | | | 0.438 | 0.478 | | 0.381 | |
| | $U$ | | | | | | | 1556.0 | | |
| $\zeta_2$ | $p_{bf}$ | — | n.s. | n.s. | n.s. | n.s. | n.s. | 0.039 | n.s. | n.s. |
| | $\lvert\delta\rvert$ | | | | | | | 0.326 | | |
| | $U$ | | | | | | 3353.5 | | | |
| $\zeta_3$ | $p_{bf}$ | — | — | n.s. | n.s. | n.s. | 0.021 | n.s. | n.s. | n.s. |
| | $\lvert\delta\rvert$ | | | | | | 0.308 | | | |
| | $U$ | | | | | | | | | |
| $\zeta_4$ | $p_{bf}$ | — | — | — | n.s. | n.s. | n.s. | n.s. | n.s. | n.s. |
| | $\lvert\delta\rvert$ | | | | | | | | | |
| | $U$ | | | | | | | | | |
| $\zeta_5$ | $p_{bf}$ | — | — | — | — | n.s. | n.s. | n.s. | n.s. | n.s. |
| | $\lvert\delta\rvert$ | | | | | | | | | |
| | $U$ | | | | | | | 1525.5 | | |
| $\zeta_6$ | $p_{bf}$ | — | — | — | — | — | n.s. | 0.001 | n.s. | n.s. |
| | $\lvert\delta\rvert$ | | | | | | | 0.403 | | |
| | $U$ | | | | | | | 3263.5 | | |
| $\zeta_7$ | $p_{bf}$ | — | — | — | — | — | — | $< 0.001$ | n.s. | n.s. |
| | $\lvert\delta\rvert$ | | | | | | | 0.459 | | |
| | $U$ | | | | | | | | 2451.5 | |
| $\zeta_8$ | $p_{bf}$ | — | — | — | — | — | — | — | $< 0.001$ | n.s. |
| | $\lvert\delta\rvert$ | | | | | | | | 0.372 | |
| | $U$ | | | | | | | | | |
| $\zeta_9$ | $p_{bf}$ | — | — | — | — | — | — | — | — | n.s. |
| | $\lvert\delta\rvert$ | | | | | | | | | |

Rows and columns correspond to each $\zeta$-cluster. n.s.: not significant. $U$: test statistic of the Mann-Whitney U test. $p_{bf}$: $p$-value after Bonferroni correction. $\delta$: Cliff's $\delta$.

#### Table S19. Comparison of PIGD scores among the $\zeta$-clusters

| | | $\zeta_2$ | $\zeta_3$ | $\zeta_4$ | $\zeta_5$ | $\zeta_6$ | $\zeta_7$ | $\zeta_8$ | $\zeta_9$ | $\zeta_{10}$ |
|---|---|---|---|---|---|---|---|---|---|---|
| $\zeta_1$ | $U$<br>$p_{bf}$<br>$\lvert\delta\rvert$ | n.s. | n.s. | n.s. | n.s. | n.s. | n.s. | n.s. | n.s. | n.s. |
| $\zeta_2$ | $U$<br>$p_{bf}$<br>$\lvert\delta\rvert$ | — | n.s. | n.s. | n.s. | n.s. | n.s. | 1161.0<br>$< 0.001$<br>0.497 | n.s. | n.s. |
| $\zeta_3$ | $U$<br>$p_{bf}$<br>$\lvert\delta\rvert$ | — | — | n.s. | n.s. | n.s. | n.s. | 1023.0<br>$< 0.001$<br>0.494 | n.s. | n.s. |
| $\zeta_4$ | $U$<br>$p_{bf}$<br>$\lvert\delta\rvert$ | — | — | — | n.s. | n.s. | n.s. | 752.5<br>0.009<br>0.427 | n.s. | n.s. |
| $\zeta_5$ | $U$<br>$p_{bf}$<br>$\lvert\delta\rvert$ | — | — | — | — | n.s. | n.s. | 411.5<br>$< 0.001$<br>0.703 | n.s. | n.s. |
| $\zeta_6$ | $U$<br>$p_{bf}$<br>$\lvert\delta\rvert$ | — | — | — | — | — | n.s. | 996.5<br>$< 0.001$<br>0.610 | 2455.0<br>$< 0.001$<br>0.380 | 945.0<br>0.008<br>0.403 |
| $\zeta_7$ | $U$<br>$p_{bf}$<br>$\lvert\delta\rvert$ | — | — | — | — | — | — | 2472.5<br>$< 0.001$<br>0.590 | 6988.0<br>0.011<br>0.253 | n.s. |
| $\zeta_8$ | $U$<br>$p_{bf}$<br>$\lvert\delta\rvert$ | — | — | — | — | — | — | — | 2214.5<br>$< 0.001$<br>0.433 | 963.5<br>0.016<br>0.383 |
| $\zeta_9$ | $U$<br>$p_{bf}$<br>$\lvert\delta\rvert$ | — | — | — | — | — | — | — | — | n.s. |

Rows and columns correspond to each $\zeta$-cluster. n.s.: not significant. $U$: test statistic of the Mann-Whitney U test. $p_{bf}$: $p$-value after Bonferroni correction. $\delta$: Cliff's $\delta$.

**Table S20. Performance evaluation of the five-layer neural network for $\mu \mapsto z$.**

| | |
|---|---|
| MSE for test data | $0.3597$ |
| Mean MSE for 9-fold cross-validation | $0.3645 \pm 0.0436$ |
| Mean prediction error of the $\boldsymbol{\mu}$-cluster centroids | $0.0361 \pm 0.0433$ |

**Table S21. Inter-centroid distances of the $\zeta$-clusters**

| | $\zeta_2$ | $\zeta_3$ | $\zeta_4$ | $\zeta_5$ | $\zeta_6$ | $\zeta_7$ | $\zeta_8$ | $\zeta_9$ | $\zeta_{10}$ |
|---|---|---|---|---|---|---|---|---|---|
| $\zeta_1$ | 1.1663 | 1.4567 | 3.8162 | 4.0225 | 2.3780 | 2.0255 | 4.7998 | 2.7670 | 1.9816 |
| $\zeta_2$ | | 0.4374 | 1.3554 | 2.2953 | 1.0981 | 0.8917 | 2.6349 | 2.2754 | 1.1157 |
| $\zeta_3$ | | | 0.6017 | 1.2272 | 0.8703 | 0.6143 | 1.7294 | 1.1800 | 1.1184 |
| $\zeta_4$ | | | | 0.8527 | 1.5416 | 1.4137 | 1.5039 | 1.8509 | 1.9133 |
| $\zeta_5$ | | | | | 2.7255 | 1.4922 | 1.4702 | 1.0969 | 1.1413 |
| $\zeta_6$ | | | | | | 0.6390 | 1.0968 | 1.3232 | 1.7049 |
| $\zeta_7$ | | | | | | | 0.7636 | 0.6528 | 0.6467 |
| $\zeta_8$ | | | | | | | | 0.7284 | 1.6150 |
| $\zeta_9$ | | | | | | | | | 1.0696 |

Distances were calculated using the mean squared distance. All values were larger than the MSE obtained when test data were input into the 5-layer NN performing $\boldsymbol{\mu} \mapsto \boldsymbol{z}$.

**Table S22. Performance evaluation of the five-layer neural network for $z \mapsto \mu$.**

| | |
|---|---|
| MSE for test data | $0.3595$ |
| Mean MSE for 9-fold cross-validation | $0.3742 \pm 0.0406$ |
| Mean prediction error of the $\zeta$-cluster centroids | $0.0466 \pm 0.0360$ |

**Table S23. Inter-centroid distances of the $\mu$-clusters**

| | $\mu_2$ | $\mu_3$ | $\mu_4$ | $\mu_5$ | $\mu_6$ | $\mu_7$ | $\mu_8$ | $\mu_9$ | $\mu_{10}$ |
|---|---|---|---|---|---|---|---|---|---|
| $\mu_1$ | 0.5270 | 0.7028 | 1.0972 | 1.4693 | 1.9211 | 2.5834 | 3.6056 | 3.9169 | 17.2729 |
| $\mu_2$ | | 0.4630 | 1.0218 | 2.2696 | 0.8027 | 1.1520 | 1.9649 | 2.9229 | 16.9470 |
| $\mu_3$ | | | 0.8871 | 2.9170 | 0.9634 | 1.8344 | 3.3531 | 4.4244 | 17.6902 |
| $\mu_4$ | | | | 1.5591 | 1.7903 | 2.0209 | 2.4997 | 2.6252 | 11.2882 |
| $\mu_5$ | | | | | 3.3835 | 3.3445 | 2.8791 | 2.1604 | 12.8684 |
| $\mu_6$ | | | | | | **0.2513** | 1.5405 | 2.4896 | 16.5418 |
| $\mu_7$ | | | | | | | 0.7396 | 1.3954 | 14.3102 |
| $\mu_8$ | | | | | | | | 0.6325 | 13.0535 |
| $\mu_9$ | | | | | | | | | 9.2846 |

Distances were calculated using the mean squared distance. Values in bold indicate distances smaller than the MSE obtained when test data were input into the 5-layer NN performing $z \mapsto \mu$.

**Table S24. Number of data used for training and prediction performance ($\mu \mapsto z$)**

| Training data | MSE for test data | MSE for mixed dataset | | | | | |
|---|---|---|---|---|---|---|---|
| | | $z_1$ | $z_2$ | $z_3$ | $z_4$ | $z_5$ | $z_6$ |
| Mixed dataset ($N = 1038$) | 0.3597 | 0.1406 | 0.0881 | 0.3708 | 0.4306 | 0.1407 | 0.4017 |
| Only real data ($N = 173$) | 0.4893 | 0.2743 | 0.2079 | 0.6521 | 0.6939 | 0.4825 | 0.7215 |

**Table S25. Number of data used for training and prediction performance ($z \mapsto \mu$)**

| Training data | MSE for test data | MSE for mixed dataset | | | | | |
|---|---|---|---|---|---|---|---|
| | | $p$ | $D$ | $\rho$ | $\Delta$ | $\sigma$ | $r$ |
| Mixed dataset ($N = 1038$) | 0.3595 | 0.3649 | 0.2541 | 0.2701 | 0.1337 | 0.0855 | 0.4712 |
| Only real data ($N = 173$) | 0.5157 | 0.8420 | 0.5722 | 0.6509 | 0.2366 | 0.3897 | 0.6776 |
